# Bacterial contribution to genesis of the novel germ line determinant *oskar*

**DOI:** 10.1101/453514

**Authors:** Leo Blondel, Tamsin E. M. Jones, Cassandra G. Extavour

## Abstract

New cellular functions and developmental processes can evolve by modifying existing genes or creating novel genes. Novel genes can arise not only via duplication or mutation but also by acquiring foreign DNA, also called horizontal gene transfer (HGT). Here we show that HGT likely contributed to the creation of a novel gene indispensable for reproduction in some insects. Long considered a novel gene with unknown origin, *oskar* has evolved to fulfil a crucial role in insect germ cell formation. Our analysis of over 100 insect Oskar sequences suggests that Oskar arose de novo *via* fusion of eukaryotic and prokaryotic sequences. This work shows that highly unusual gene origin processes can give rise to novel genes that can facilitate evolution of novel developmental mechanisms.

**One Sentence Summary:** Our research shows that gene origin processes often considered highly unusual, including HGT and de novo coding region evolution, can give rise to novel genes that can both participate in pre-existing gene regulatory networks, and also facilitate the evolution of novel developmental mechanisms.

## Main Text

Heritable variation is the raw material of evolutionary change. Genetic variation can arise from mutation and gene duplication of existing genes (*1*), or through *de novo* processes (*2*), but the extent to which such novel, or “orphan” genes participate significantly in the evolutionary process is unclear. Mutation of existing cis-regulatory (*3*) or protein coding regions (*4*) can drive evolutionary change in developmental processes. However, recent studies in animals and fungi suggest that novel genes can also drive phenotypic change (*5*). Although counterintuitive, novel genes may be integrating continuously into otherwise conserved gene networks, with a higher rate of partner acquisition than subtler variations on preexisting genes (*6*). Moreover, in humans and fruit flies, a large proportion of novel genes are expressed in the brain, suggesting their participation in the evolution of major organ systems (*7, 8*). However, while next generation sequencing has improved their discovery, the developmental and evolutionary significance of novel genes remains understudied.

The mechanism of formation of a novel gene may have implications for its function. Novel genes that arise by duplication, thus possessing the same biophysical properties as their parent genes, have innate potential to participate in preexisting cellular and molecular mechanisms (*1*). However, orphan genes lacking sequence similarity to existing genes must form novel functional molecular relationships with extant genes, in order to persist in the genome. When such genes arise by introduction of foreign DNA into a host genome through horizontal gene transfer (HGT), they may introduce novel, already functional sequence information into a genome. Whether genes created by HGT show a greater propensity to contribute to or enable novel processes is unclear. Endosymbionts in the host germ line cytoplasm (germ line symbionts) could increase the occurrence of evolutionarily relevant HGT events, as foreign DNA integrated into the germ line genome is transferred to the next generation. HGT from bacterial endosymbionts into insect genomes appears widespread, involving transfer of metabolic genes or even larger genomic fragments to the host genome (see for example *9, 10–12*).

Here we examined the evolutionary origins of the *oskar* (*osk*) gene, long considered a novel gene that evolved to be indispensable for insect reproduction (*13*). First discovered in *Drosophila melanogaster* (*14*), *osk* is necessary and sufficient for assembly of germ plasm, a cytoplasmic determinant that specifies the germ line in the embryo. Germ plasm-based germ line specification appears derived within insects, confined to insects that undergo metamorphosis (Holometabola) (*15, 16*). Initially thought exclusive to Diptera (flies and mosquitoes), its discovery in a wasp, another holometabolous insect with germ plasm (*17*), led to the hypothesis that *oskar* originated as a novel gene at the base of the Holometabola approximately 300 Mya, facilitating the evolution of insect germ plasm as a novel developmental mechanism (*17*). However, its subsequent discovery in a cricket (*15*), a hemimetabolous insect without germ plasm (*18*), implied that *osk* was instead at least 50 My older, and that its germ plasm role was derived rather than ancestral (*19*). Despite its orphan gene status, *osk* plays major developmental roles, interacting with the products of many genes highly conserved across animals (*13, 20, 21*). *osk* thus represents an example of a novel gene that not only functions within pre-existing gene networks in the nervous system (*15*), but has also evolved into the only animal gene known to be both necessary and sufficient for germ line specification (*22, 23*).

The evolutionary origins of this remarkable gene are unknown. Osk contains two biophysically conserved domains, an N-terminal LOTUS domain and a C-terminal hydrolase-like domain called OSK (*20, 24*) (Figure 1a). An initial BLASTp search using the full-length *D. melanogaster osk* sequence as a query yielded either other holometabolous insect *osk* genes, or partial hits for the LOTUS or OSK domains (E-value < 0.01; Supplementary files: BLAST search results). This suggested that full length *osk* was unlikely to be a duplication of any other known gene. This prompted us to perform two more BLASTp searches, one using each of the two conserved Osk protein domains individually as query sequences. Strikingly, in this BLASTp search, although we recovered several eukaryotic hits for the LOTUS domain, we recovered no eukaryotic sequences that resembled the OSK domain, even with very low E-value stringency (E-value < 10; see Methods section “*BLAST searches of oskar”* for an explanation of E-value threshold choices; Supplementary files: BLAST search results).

**Figure 1.**
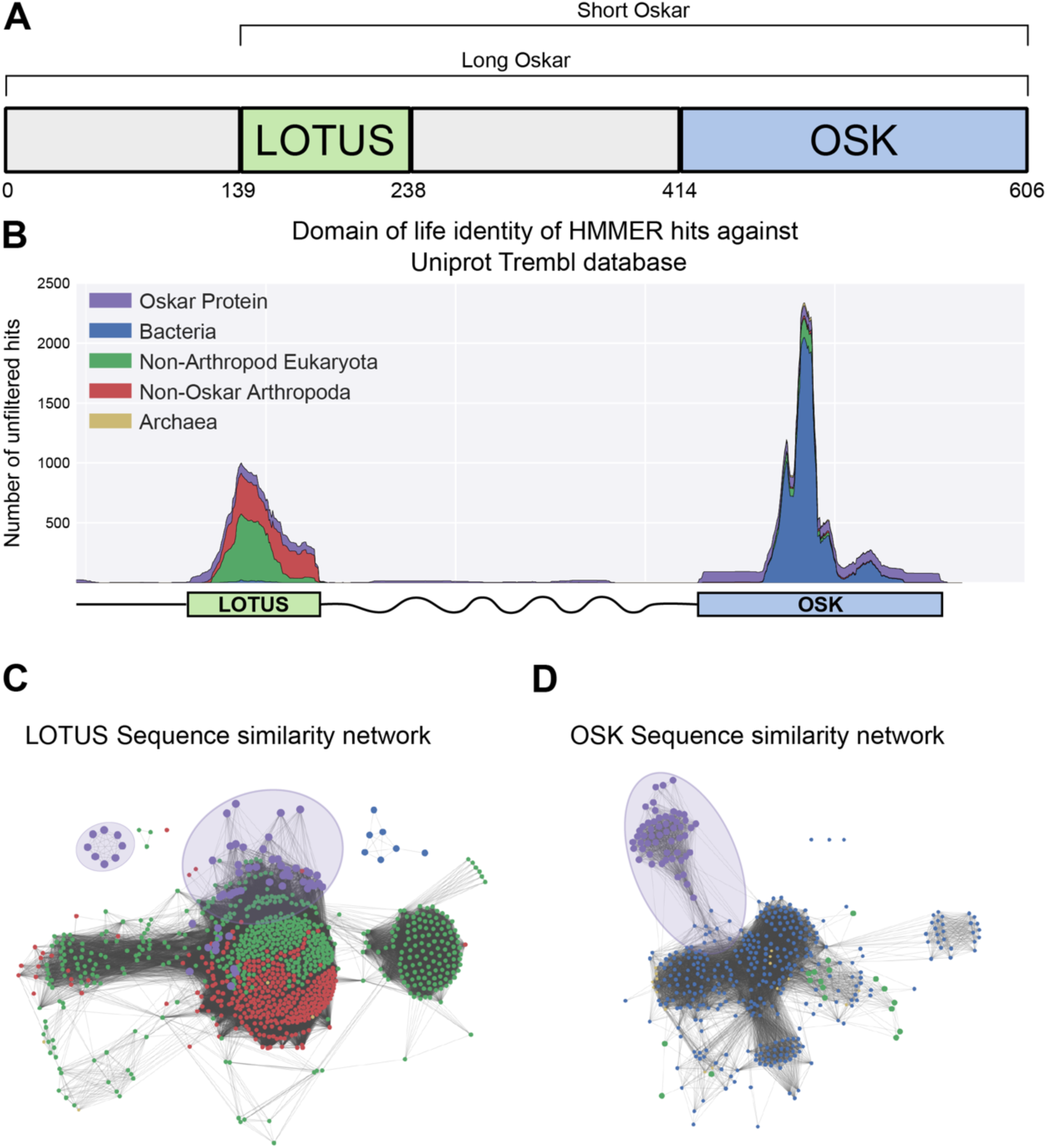
Sequence analysis of the Oskar gene. **a**, Schematic representation of the Oskar gene. The LOTUS and OSK hydrolase-like domains are separated by a poorly conserved region of predicted high disorder and variable length between species. In some dipterans, a region 5’ to the LOTUS domain is translated to yield a second isoform, called Long Oskar. Residue numbers correspond to the *D. melanogaster* Osk sequence. **b**, Stackplot of domain of life identity of HMMER hits across the protein sequence. For a sliding window of 60 Amino Acids across the protein sequence (X axis), the number of hits in the Trembl (UniProt) database (Y axis) is represented and color coded by domain of life origin (see Methods: Iterative HMMER search of OSK and LOTUS domains), stacked on top of each other. **c, d** EFI-EST-generated graphs of the sequence similarity network of the LOTUS (**c**) and OSK (**d**) domains of Oskar (*73*). Sequences were obtained using HMMER against the UniProtKB database. Most Oskar LOTUS sequences cluster within eukaryotes and arthropods. In contrast, Oskar OSK sequences cluster most strongly with a small subset of bacterial sequences.

To understand this anomaly, we built an alignment of 95 Oskar sequences (Supplementary files: Alignments>OSKAR_MUSCLE_FINAL.fasta; Supplementary Table S2) and used a custom iterative HMMER sliding window search tool to compare each domain with protein sequences from all domains of life. Sequences most similar to the LOTUS domain were almost exclusively eukaryotic sequences (Supplementary Table 3). In contrast, those most similar to the OSK domain were bacterial, specifically sequences similar to SGNH-like hydrolases (*20, 24*) (Pfam Clan: SGNH_hydrolase - CL0264; Supp. Table 4; Figure 1b). To visualize their relationships, we graphed the sequence similarity network for the sequences of these domains and their closest hits. We observed that the majority of LOTUS domain sequences clustered within eukaryotic sequences (Figure 1c). In contrast, OSK domain sequences formed an isolated cluster, a small subset of which formed a connection to bacterial sequences (Figure 1d). These data are consistent with a previous suggestion, based on BLAST results (*17*), that HGT from a bacterium into an ancestral insect genome may have contributed to the evolution of *osk*. However, this possibility was not adequately addressed by previous analyses, which were based on alignments of full length Osk containing only eukaryotic sequences as outgroups (*15*). To rigorously test this hypothesis, we therefore performed phylogenetic analyses of the two domains independently. A finding that LOTUS sequences were nested within eukaryotes, while OSK sequences were nested within bacteria, would provide support for the HGT hypothesis.

Both Maximum likelihood and Bayesian approaches confirmed this prediction (Figure 2a, Figure 2–figure supplements 1 and 2), and these results were robust to changes in the methods of sequence alignment (Figure 2-figure supplements 6-10). As expected, LOTUS sequences from Osk proteins were related to other eukaryotic LOTUS domains, to the exclusion of the only three bacterial sequences that met our E-value cutoff for inclusion in the analyses (Figs. 2a, Figure 2–figure supplements 1 and 2; see Methods and Supplemental Text). LOTUS sequences from non-Oskar proteins were almost exclusively eukaryotic. (Supplementary Table 3); only three bacterial sequences matched the LOTUS domain with an E-value < 0.01. Osk LOTUS domains clustered into two distinct clades, one comprising all Dipteran sequences, and the other comprising all other Osk LOTUS domains examined from both holometabolous and hemimetabolous orders (Figure 2a). Dipteran Osk LOTUS sequences formed a monophyletic group that branched sister to a clade of LOTUS domains from Tud5 family proteins of non-arthropod animals (NAA). NAA LOTUS domains from Tud7 family members were polyphyletic, but most of them formed a clade branching sister to (Osk LOTUS + NAA Tud5 LOTUS). Non-Dipteran Osk LOTUS domains formed a monophyletic group that was related in a polytomy to the aforementioned (NAA Tud7 LOTUS + (Dipteran Osk LOTUS + NAA Tud5 LOTUS)) clade, and to various arthropod Tud7 family LOTUS domains.

**Figure 2.**
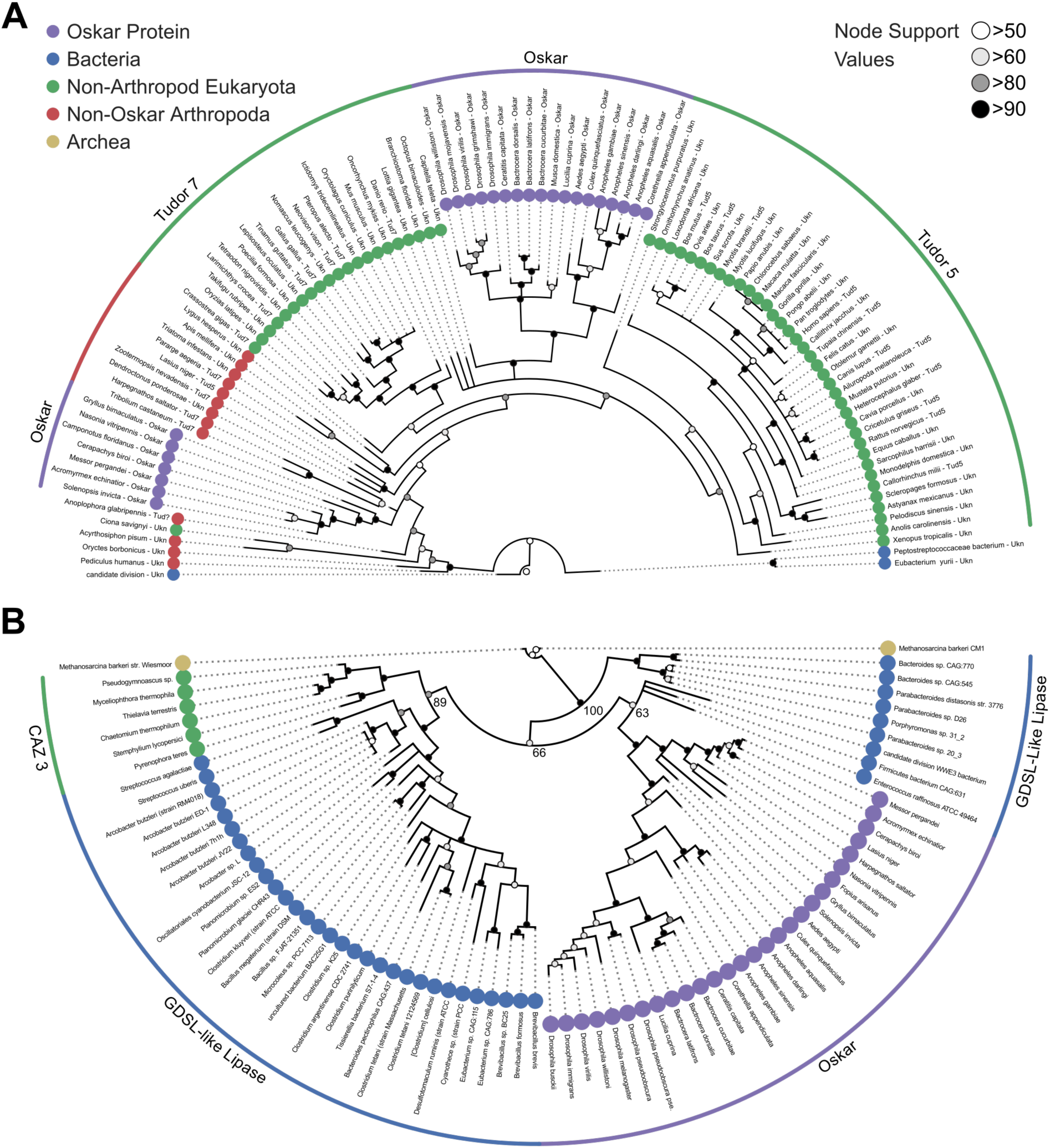
Phylogenetic analysis of the LOTUS and OSK domains. **a**, Bayesian consensus tree for the LOTUS domain. Three major LOTUS-containing protein families are represented within the tree: Tudor 5, Tudor 7, and Oskar. Oskar LOTUS domains form two clades, one containing only dipterans and one containing all other represented insects (hymenopterans and orthopterans). The tree was rooted to the three bacterial sequences added in the dataset. **b**, Bayesian consensus tree for the OSK domain. The OSK domain is nested within GDSL-like domains of bacterial species from phyla known to contain germ line symbionts in insects. The ten non-Oskar eukaryotic sequences in the analysis form one clade comprising fungal Carbohydrate Active Enzyme 3 (CAZ3) proteins. For Bayesian and RaxML trees with all accession numbers and node support values see Figure 2–figure supplement 1-4.

The fact that Tud7 LOTUS domains are polyphyletic suggests that arthropod domains in this family may have evolved differently than their homologues in other animals. The relationships of Dipteran LOTUS sequences were consistent with the current hypothesis for interrelationships between Dipteran species (*25*) Similarly, among the non-Dipteran Osk LOTUS sequences, the hymenopteran sequences form a clade to the exclusion of the single hemimetabolous sequence (from the cricket *Gryllus bimaculatus*), consistent with the monophyly of Hymenoptera (*26*). It is unclear why Dipteran Osk LOTUS domains cluster separately from those of other insect Osk proteins. We speculate that the evolution of the Long Oskar domain (*27, 28*), which appears to be a novelty within Diptera (Supplementary Files: Alignments>OSKAR_MUSCLE_FINAL.fasta), may have influenced the evolution of the Osk LOTUS domain in at least some of these insects. Consistent with this hypothesis, of the 17 Dipteran *oskar* genes we examined, the seven *oskar* genes possessing a Long Osk domain clustered into two clades based on the sequences of their LOTUS domain. One of these clades comprised five Drosophila species (*D. willistoni*, *D. mojavensis*, *D. virilis*, *D. grimshawi* and *D. immigrans*), and the second was composed of two calyptrate flies from different superfamilies, *Musca domestica* (Muscoidea) and *Lucilia cuprina* (Oestroidea).

In summary, the LOTUS domain of Osk proteins is most closely related to a number of other LOTUS domains found in eukaryotic proteins, as would be expected for a gene of animal origin, and the phylogenetic interrelationships of these sequences is largely consistent with the current species or family level trees for the corresponding insects.

In contrast, OSK domain sequences were nested within bacterial sequences (Figure 2b, Figure 2–figure supplements 3, 4). This bacterial, rather than eukaryotic, affinity of the OSK domain was recovered even when different sequence alignment methods were used (Figure 2-figure supplements 7-11). The only eukaryotic proteins emerging from the iterative HMMER search for OSK domain sequences that had an E-value < 0.01 were all from fungi. All five of these sequences were annotated as Carbohydrate Active Enzyme 3 (CAZ3), and all CAZ3 sequences formed a clade that was sister to a clade of primarily Firmicutes. Most bacterial sequences used in this analysis were annotated as lipases and hydrolases, with a high representation of GDSL-like hydrolases (Supplementary Table S4). OSK sequences formed a monophyletic group but did not branch sister to the other eukaryotic sequences in the analysis. Within this OSK clade, the topology of sequence relationships was largely concordant with the species tree for insects (*29*), as we recovered monophyletic Diptera to the exclusion of other insect species. However, the single orthopteran OSK sequence (from the cricket *Gryllus bimaculatus*) grouped within the Hymenoptera, rather than branching as sister to all other insect sequences in the tree, as would be expected for this hemimetabolous sequence (*29*).

Importantly, OSK sequences did not simply form an outgroup to bacterial sequences. To formally reject the possibility that the eukaryotic OSK clade has a sister group relationship to all bacterial sequences in the analysis, we performed topology constraint analyses using the Swofford–Olsen– Waddell–Hillis (SOWH) test, which assigns statistical support to alternative phylogenetic topologies (*30*). We used the SOWHAT tool (*31*) to compare the HGT-supporting topology to two alternative topologies with constraints more consistent with vertical inheritance. The first was constrained by domain of life, disallowing paraphyletic relationships between sequences from the same domain of life (Figure 2–figure supplement 5a). The second required monophyly of Eukaryota, but allowed paraphyletic relationships between bacterial and archaeal sequences (Figure 2–figure supplement 5b). We found that the topologies of both of these constrained trees were significantly worse than the result we had recovered with our phylogenetic analysis (Figure 2–figure supplement 5), namely that the closest relatives of the OSK domain were bacterial rather than eukaryotic sequences (Figure 2b, Figure 2–figure supplements 3 and 4).

OSK sequences formed a well-supported clade nested within bacterial GDSL-like lipase sequences. The majority of these bacterial sequences were from the Firmicutes, a bacterial phylum known to include insect germline symbionts (*32, 33*). All other sequences from classified bacterial species, including a clade branching as sister to all other sequences, belonged either to the Bacteroidetes or to the Proteobacteria. Members of both of these phyla are also known germline symbionts of insects (*9, 34*) and other arthropods (*35*). In sum, the distinct phylogenetic relationships of the two domains of Oskar are consistent with a bacterial origin for the OSK domain. Further, the specific bacterial clades close to OSK suggest that an ancient arthropod germ line endosymbiont could have been the source of a GDSL-like sequence that was transferred into an ancestral insect genome, and ultimately gave rise to the OSK domain of *oskar*.

We then asked if two additional sequence characteristics, GC3 content and codon use, were consistent with distinct domain of life origins for the two Oskar domains (*36*). Under our hypothesis, the HGT event that contributed to *oskar*’s formation would have occurred at least 480 Mya, in a common insect ancestor (*29*). This means that we would be highly unlikely to be able to definitively identify any extant bacterial species from which the OSK sequence were putatively derived, as has been possible, for example, with much more recent HGT events on the order of a few million years ago (see for example *9*). However, we reasoned that if evolutionary time had not completely erased the original GC3 content and codon use signatures from the putative bacterially donated sequence (OSK), we might detect some differences in these parameters both from the LOTUS domain, and from the host genome. Thus, we performed a parametric analysis of these parameters for 17 well annotated insect genomes (Supplementary Table 5).

To quantify the null hypothesis, we calculated an “Intra-Gene distribution” for all genes in the genome, which assessed the correlation between codon use in the 5’ and 3’ halves of a given gene. To determine whether the AT3/GC3 content correlation between the OSK and LOTUS domains of *oskar* was different from that between the 5’ and 3’ ends of an average gene in the genome, we then calculated the residuals of the Intra-Gene distribution and the LOTUS-OSK domain pair. Pooling the residuals together revealed that the GC3 content was significantly different between the LOTUS and OSK domains, compared to what would be expected within an average gene in that genome (Figure 3a-c).

**Figure 3.**
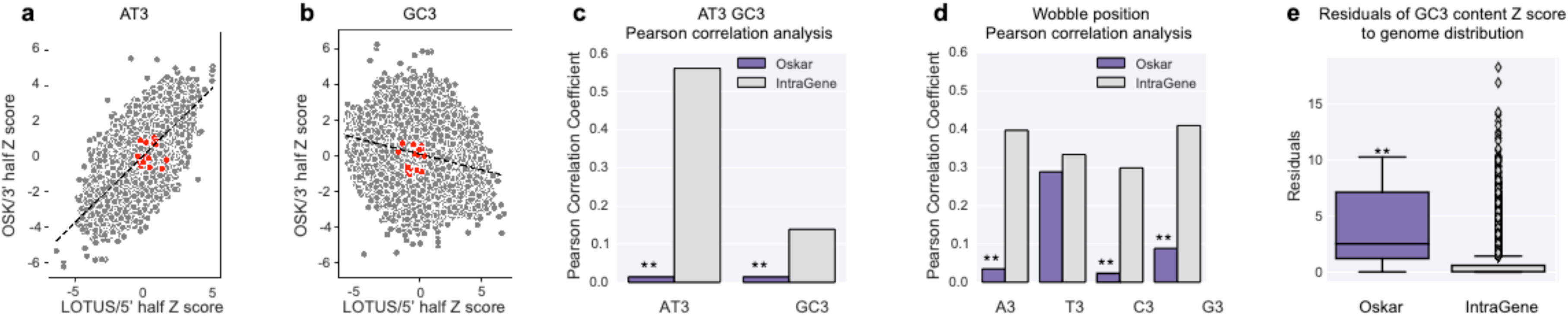
Parametric analysis of codon use for the LOTUS and OSK domains. **a**, Scatter plot representing the full distribution of correlations from Pearson correlation analysis for AT3 use in 3’ and 5’ halves of all non-*oskar* genes in the 17 genomes analyzed (grey), compared to the correlation for the OSK and LOTUS domains of *oskar* (red). The Z score ranges of correlations for the *oskar* domains are within the distribution for all genes in the genome, although correlations of use across the gene are lower for *oskar* than for non-*oskar* genes (see also **c**). **b**, Scatter plot representing the full distribution of correlations from Pearson correlation analysis for GC3 use in 3’ and 5’ halves of all non-*oskar* genes in the 17 genomes analyzed (grey), compared to the correlation for the OSK and LOTUS domains of *oskar* (red). The Z score ranges of correlations for the *oskar* domains are within the distribution for all genes in the genome, although correlations of use across the gene are lower for *oskar* than for non-*oskar* genes (see also **c**). **c**, Pearson correlation analysis of AT3 and GC3 content for Oskar vs other genes. AT3 and GC3 content are correlated across the 5’ and 3’ halves of a gene for all genes in a given genome (grey), but not between the LOTUS and OSK domains of Oskar (purple). (**: Pearson correlation p-value > 0.1, AT3 r=0.014 p=0.646 and GC3 r=0.014 and p=0.646) **d**, Pearson correlation analysis of wobble position identity for the Oskar gene vs other genes. Wobble position identity content is correlated across the 5’ and 3’ halves of a gene for all genes in a given genome (grey) but not between the LOTUS and OSK domains of Oskar (purple), with the exception of T3. (**: Pearson correlation p-value > 0.1, A3 r=0.034 p=0.476, T3 r=0.289 p=0.026, C3 r=0.023 p=0.560 and G3 r=0.088 p=0.247) **e**, Analysis of GC3 content. Measure of the residuals of Z scores for Oskar gene GC3 content (LOTUS vs OSK: purple) and the IntraGene GC3 content (grey). The GC3 content of the LOTUS and OSK domains does not follow a linear relationship, and the residuals are significantly higher (purple) than those observed across the 5’ and 3’ halves of other genes within a given genome (grey). (** : Mann-Whitney U test p-value < 10^-5^ p=1.927.10^-7^).

To ask if codon use was different between the OSK domain and other genes in the genome, we used a modified “codon adaptation index” (CAI) approach (37). We calculated the frequency of codon use for every gene in the available transcriptomes, and calculated a “global CAI” for all genes in each of these species (rather than only for highly expressed genes, since we cannot be confident which ones those are based on the available data for these species). We compared the genome-wide CAI distribution with the CAI values of the full length oskar mRNA, the LOTUS domain and the OSK domain, and found that the latter three CAI values fell well within the ranges of the genome-wide CAI distribution, for all 17 genomes studied (Figure 3 Supplement 6A). Similarly, the differences between the LOTUS and OSK domain CAI values were not significantly different from those between the 5’ and 3’ ends of each of the other genes in the genome (Figure 3 Supplement 6B), and the OSK and LOTUS CAI values were similar to median values for the 5’ and 3’ ends of other genes in the same genome (Figure 3 Supplement 6B). In sum, while the overall codon use of the OSK and LOTUS domains of *oskar* appears similar to each other and to the codon use of other genes in the same genome, the GC3 content is significantly different between the two domains, and from that of an average gene in the genome.

While the results of our GC3 content analyses are consistent with the hypothesis of an HGT origin for *oskar*, we cannot exclude the possibility that other mechanisms might impact this codon use difference between two different regions of *oskar*. For example, it is possible that a form of translational selection might be acting to promote differential codon use in distinct regions of *oskar* (*38*), as a consequence of different folding requirements for different regions of the protein (*39, 40*). Codon use may influence differential translation of ordered versus disordered regions of the same protein (*41*), and Oskar contains regions of high predicted disorder in addition to the LOTUS and OSK domains (*20, 42*). However, in the present study we have examined only the latter two domains, and to our knowledge there is no evidence of differential translational regulation of these two well-folded domains of Oskar.

An alternative parameter that could impact codon use across the *oskar* gene is a possible requirement for specific structures of the *oskar* mRNA. Given that the *oskar* transcript plays a translation-independent role in the *Drosophila* germ line (*43, 44*), it is conceivable that structural constraints might be different on different regions of its mRNA. Moreover, specific stem-loop structures within the oskar 3’ UTR are required for its localization within the oocyte (*45–47*), consistent with structural constraints on *oskar* sequence evolution. However, we are unaware of any evidence suggesting specific roles for the structure of the coding region of the *oskar* transcript. Thus, taken together with the sequence similarity and strong phylogenetic evidence presented above, we propose that the results of these analyses support an HGT origin for the OSK domain (Figure 4).

**Figure 4.**
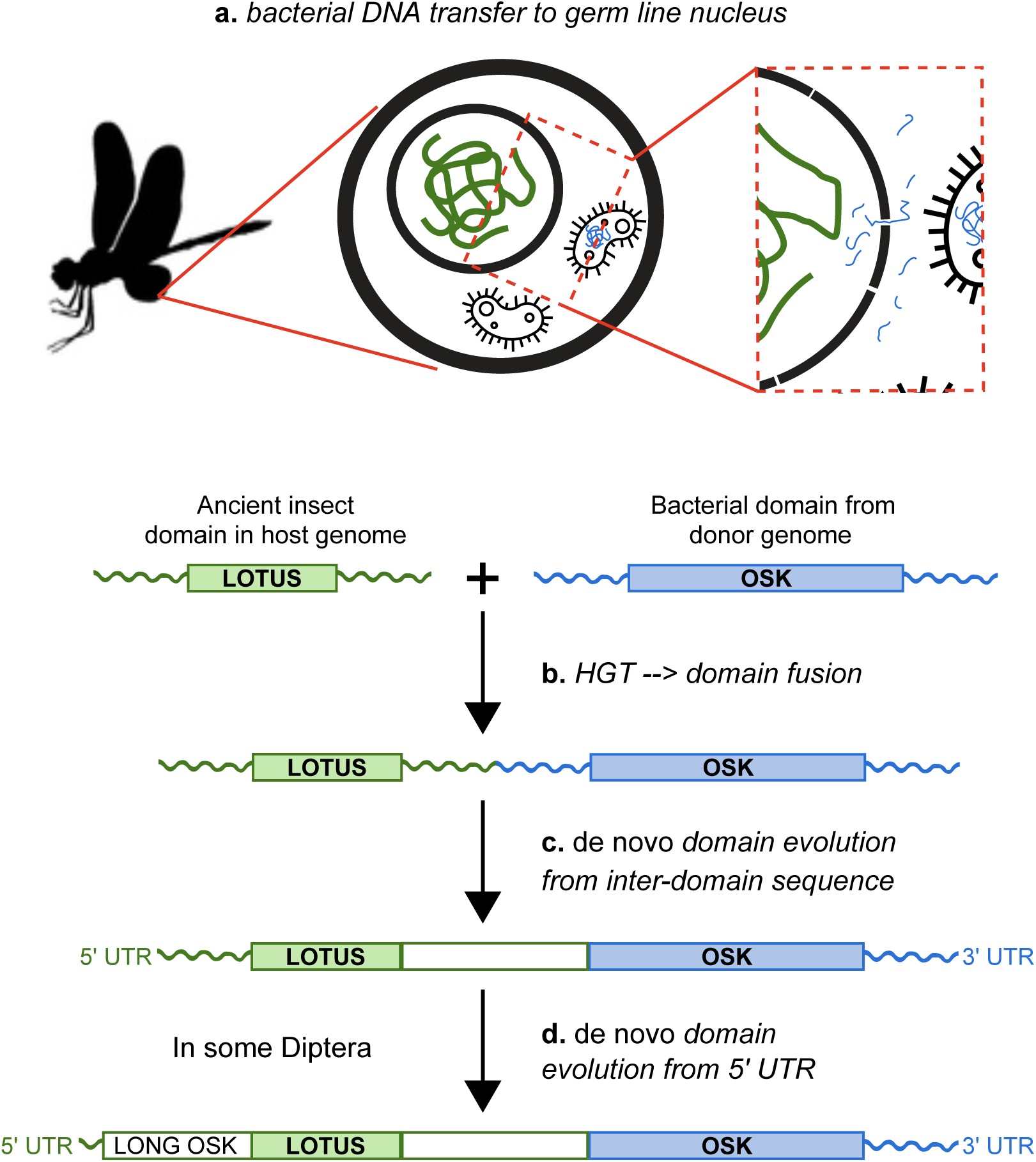
Hypothesis for the origin of *oskar*. Integration of the OSK domain close to a LOTUS domain in an ancestral insect genome. **a**, DNA containing a GDSL-like domain from an endosymbiotic germ line bacterium is transferred to the nucleus of a germ cell in an insect common ancestor. **b**, DNA damage or transposable element activity induces an integration event in the host genome, close to a pre-existing LOTUS-like domain. **c**, The region between the two domains undergoes *de novo* coding evolution, creating an open reading frame with a unique, chimeric domain structure. **d**, In some Diptera, including *D. melanogaster*, part of the 5’ UTR of *oskar* has undergone *de novo* coding evolution to form the Long Oskar domain.

While multiple mechanisms can give rise to novel genes, HGT is arguably among the least well understood, as it involves multiple genomes and ancient biotic interactions between donor and host organisms that are often difficult to reconstruct. In the case of *oskar*, however, the fact that both germline symbionts (*48*) and HGT events (*9*) are widespread in insects, provides a plausible biological mechanism consistent with our hypothesis that fusion of eukaryotic and bacterial domain sequences led to the birth of this novel gene. Under this hypothesis, this fusion would have taken place before the major diversification of insects, nearly 500 million years ago (*29*).

Once arisen, novel genes might be expected to disappear rapidly, given that pre-existing gene regulatory networks operated successfully without them (*1*). However, it is clear that novel genes can evolve functional connections with existing networks, become essential (*49*), and in some cases lead to new functions (*50*) and contribute to phenotypic diversity (*5*). Even given the growing number of convincing examples of HGT from both prokaryotic and eukaryotic origins (see for example *51, 52–54*), some authors suspect that the contribution of horizontal gene transfer to the acquisition of novel traits has been underestimated across animals (*55*). Moreover, the functional contribution of genes horizontally transferred specifically from bacteria to insects has been documented for a range of adaptive phenotypes (see for example *56, 57, 58*), including digestive metabolism (*10, 11, 59*), glycolysis (*60*) complex symbiosis (*12*) and endosymbiont cell wall construction (*61*). *oskar* plays multiple critical roles in insect development, from neural patterning (*15, 62*) to oogenesis (*43*). In the Holometabola, a clade of nearly one million extant species (*63*), *oskar*’s co-option to become necessary and sufficient for germ plasm assembly is likely the cell biological mechanism underlying the evolution of this derived mode of insect germ line specification (*15, 17, 19*). Our study thus provides evidence that HGT can not only introduce functional genes into a host genome, but also, by contributing sequences of individual domains, generate genes with entirely novel domain structures that may facilitate the evolution of novel developmental mechanisms.

## Methods

### BLAST searches of Oskar

All BLAST (*64*) searches were performed using the NCBI BLASTp tool suite on the non-redundant (nr) database. Amino Acid (AA) sequences of *D. melanogaster* full length Oskar (EMBL ID AAF54306.1), as well as the AA sequences for the *D. melanogaster* Oskar LOTUS (AA 139-238) and OSK (AA 414-606) domains were used for the BLAST searches. We used the default NCBI cut-off parameters (E-value cut-off of 10) for searches using OSK and LOTUS as queries, and a more stringent E-value threshhold of 0.01 for the search using full length *D. melanogaster* Oskar as a query. We chose an E-value threshold of 10 for LOTUS and OSK to capture potentially highly divergent homologs of the two domains, especially for the OSK domain, where we were looking for any viable candidate for a homologous eukaryotic domain. All BLAST searches results are included in the Supplementary files: BLAST search results.

### Hidden Markov Model (HMM) generation and alignments of the OSK and LOTUS domains

101 1KITE transcriptomes (*65*) (Supplementary Table 1) were downloaded and searched using the local BLAST program (BLAST+) using the tblastn algorithm with default parameters, with Oskar protein sequences of *Drosophila melanogaster, Aedes aegypti, Nasonia vitripennis* and *Gryllus bimaculatus* as queries (EntrezIDs: NP_731295.1, ABC41128.1, NP_001234884.1 and AFV31610.1 respectively). For all of these 1KITE transcriptome searches, predicted protein sequences from transcript data were obtained by in silico translation using the online ExPASy translate tool (https://web.expasy.org/translate/), taking the longest open reading frame. Publicly available sequences in the non-redundant (nr), TSA databases at NCBI, and a then-unpublished transcriptome (*66*) (kind gift of Matthew Benton and Siegfried Roth, University of Cologne) were subsequently searched using the web-based BLAST tool hosted at NCBI, using the tblastn algorithm with default parameters. Sequences used for queries were the four Oskar proteins described above, and newfound *oskar* sequences from the 1KITE transcriptomes of *Baetis pumilis, Cryptocercus wright,* and *Frankliniella cephalica*. For both searches, *oskar* orthologs were identified by the presence of BLAST hits on the same transcript to both the LOTUS (N-terminal) and OSK (C-terminal) regions of any of the query *oskar* sequences, regardless of E-values. The sequences found were aligned using MUSCLE (8 iterations) (*67*) into a 46-sequence alignment (Supplementary files: Alignments>OSKAR_MUSCLE_INITIAL.fasta). From this alignment, the LOTUS and OSK domains were extracted (Supplementary files: Alignments>LOTUS_MUSCLE_INITIAL.fasta and Alignments>OSK_MUSCLE_INITIAL.fasta) to define the initial Hidden Markov Models (HMM) using the hmmbuild tool from the HMMER tool suite with default parameters (*68*). 126 insect genomes and 128 insect transcriptomes (from the Transcriptome Shotgun Assembly TSA database: https://www.ncbi.nlm.nih.gov/Traces/wgs/?view=TSA) were subsequently downloaded from NCBI (download date September 29, 2015; Supplementary table 1). Genomes were submitted to Augustus v2.5.5 (*69*) (using the *D. melanogaster* exon HMM predictor) and SNAP v2006-07-28 (*70*) (using the default ‘fly’ HMM) for gene discovery. The resulting nucleotide sequence database comprising all 309 downloaded and annotated genomes and transcriptomes, was then translated in six frames to generate a non-redundant amino acid database (where all sequences with the same amino acid content are merged into one). This process was automated using a series of custom scripts available here: https://github.com/Xqua/Genomes. The non-redundant amino acid database was searched using the HMMER v3.1 tool suite (*68*) and the HMM for the LOTUS and OSK domains described above. A hit was considered positive if it consisted of a contiguous sequence containing both a LOTUS domain and an OSK domain, with the two domains separated by an inter-domain sequence. We imposed no length, alignment or conservation criteria on the inter-domain sequence, as this is a rapidly-evolving region of Oskar protein with predicted high disorder (*20, 24, 71*). Positive hits were manually curated and added to the main alignment, and the search was performed iteratively until no more new sequences meeting the above criteria were discovered. This resulted in a total of 95 Oskar protein sequences, (see Supplementary Table 2 for the complete list). Using the final resulting alignment (Supplementary Files: Alignments>OSKAR_MUSCLE_FINAL.fasta), the LOTUS and OSK domains were extracted from these sequences (Supplementary Files: Alignments>LOTUS_MUSCLE_FINAL.fasta and Alignments>OSK_MUSCLE_FINAL.fasta), and the final three HMM (for full-length Oskar, OSK, and LOTUS domains) used in subsequent analyses were created using hmmbuild with default parameters (Supplementary files: HMM>OSK.hmm, HMM>LOTUS.hmm and HMM>OSKAR.hmm).

### Iterative HMMER search of OSK and LOTUS domains

A reduced version of TrEMBL (*72*) (v2016-06) was created by concatenating all hits (regardless of E-value) for sequences of the LOTUS domain, the OSK domain and full-length Oskar, using hmmsearch with default parameters and the HMM models created above from the final alignment. This reduced database was created to reduce potential false positive results that might result from the limited size of the sliding window used in the search approach described here. The full-length Oskar alignment of 1133 amino acids (Supplementary files: Alignments>OSKAR_MUSCLE_FINAL.fasta) was split into 934 sub-alignments of 60 amino acids each using a sliding window of one amino acid. Each alignment was converted into a HMM using hmmbuild, and searched against the reduced TrEMBL database using hmmsearch using default parameters. Domain of life origin of every hit sequence at each position was recorded. Eukaryotic sequences were further classified as Oskar/Non-Oskar and Arthropod/Non-Arthropod. Finally, for the whole alignment, the counts for each category were saved and plotted in a stack plot representing the proportion of sequences from each category to create Figure 1b. The python code used for this search is available at https://github.com/Xqua/Iterative-HMMER.

### Sequence Similarity Networks

LOTUS and OSK domain sequences from the final alignment obtained as described above (see “*Hidden Markov Model (HMM) generation and alignments of the OSK and LOTUS domains*”; Supplementary files: Alignments>LOTUS_MUSCLE_FINAL.fasta and Alignments>OSK_MUSCLE_FINAL.fasta) were searched against TrEMBL (*72*) (v2016-06) using HMMER. All hits with E-value < 0.01 were consolidated into a fasta file that was then entered into the EFI-EST tool (*73*) using default parameters to generate a sequence similarity network. An alignment score corresponding to 30% sequence identity was chosen for the generation of the final sequence similarity network. Finally, the network was graphed using Cytoscape 3 (*74*).

### Phylogenetic Analysis Based on MUSCLE Alignment

For both the LOTUS and OSK domains, in cases where more than one sequence from the same organism was retrieved by the search described above in “*Iterative HMMER Search of OSK and LOTUS domains*”, only the sequence with the lowest E-value was used for phylogenetic analysis. For the LOTUS domain, the first 97 best hits (lowest E-value) were selected, and the only three bacterial sequences that satisfied an E-value < 0.01 were manually added. For *oskar* sequences, if more than one sequence per species was obtained by the search, only the single sequence per species with the lowest E-value was kept for analysis, generating a set of 100 sequences for the LOTUS domain, and 87 sequences for the OSK domain. Unique identifiers for all sequences used to generate alignments for phylogenetic analysis are available in Supplementary Tables S3, S4. For both datasets, the sequences were then aligned using MUSCLE (*67*) (8 iterations) and trimmed using trimAl (*75*) with 70% occupancy. The resulting alignments that were subject to phylogenetic analysis are available in Supplementary Files: Alignments>LOTUS_MUSCLE_TREE.fasta and Alignments>OSK_MUSCLE_TREE.fasta. For the maximum likelihood tree, we used RaxML v8.2.4 (*76*) with 1000 bootstraps, and the models were selected using the automatic RaxML model selection tool. The substitution model chosen for both domains was LGF. For the Bayesian tree inference, we used MrBayes V3.2.6 (*77*) with a Mixed model (prset aamodel=Mixed) and a gamma distribution (lset rates=Gamma). We ran the MonteCarlo for 4 million generations (std < 0.01) for the OSK domain, and for 3 million generations (std < 0.01) for the LOTUS domain. For the tree comparisons (Fig2 supplementary figures 8-9), the RaxML best tree output from the MUSCLE and PRANK alignments were compared using the tool Phylo.io (*78*).

### Phylogenetic analysis based on PRANK alignment

For the OSK domain, the raw full length sequences obtained from the HMMER search were aligned to each other using HMMER HMM based alignment tool: hmmalign, with the same HMM used to do the search, namely OSK.hmm (supplementary data: Data/HMM/OSK.hmm). Starting from this base alignment, we used the default alignment method option offered by PRANK (version: v.170427) ((*79*). We then used PRANK to realign those sequences, which in turn led to a usable alignment for phylogenetic analysis. This alignment was trimmed using the same parameters as described in *Hidden Markov Model (HMM) generation and alignments of the OSK and LOTUS domains* above. The final alignment is available in supplementary data: Alignment/OSK_prank_aligned.fasta. We then performed a phylogenetic analysis of this alignment using RAXML with the same parameters described in *Phylogenetic Analysis Based on MUSCLE Alignment* above. The resulting tree is presented in Figure 2 Supplementary figures 7 and 8.

For the LOTUS domain, the raw full length sequences obtained from the HMMER search were aligned to each other using the HMMER HMM based alignment tool: hmmalign, with the same HMM used to do the search, namely LOTUS.hmm (Supplementary data: Data/HMM/LOTUS.hmm). Starting from this base alignment, we then used PRANK with default options to realign those sequences. This alignment was trimmed using the same parameters as described in the *Hidden Markov Model (HMM) generation and alignments of the OSK and LOTUS domains*. The final alignment is available in supplementary data: Alignments/LOTUS_prank_aligned.fasta. We then performed a phylogenetic analysis using RAXML with the same parameters described above in *Phylogenetic Analysis Based on MUSCLE alignment*. The resulting trees are presented in Figure 2 Supplementary figures 6 and 9.

### Phylogenetic Analysis Based on T Coffee alignment

For the LOTUS and OSK domain, the raw full length sequences obtained from the HMMER search were aligned to each other using T-Coffee with its default parameters (*80*). This alignment was trimmed using the same parameters as described in *Hidden Markov Model (HMM) generation and alignments of the OSK and LOTUS domains* above. The final alignment is available in supplementary data: Alignment/LOTUS_tcoffee_aligned.fasta Alignment/OSK_tcoffee_aligned.fasta. We then performed a phylogenetic analysis of this alignment using RAXML with the same parameters described in *Phylogenetic Analysis Based on MUSCLE Alignment* above. The resulting trees are presented in Figure 2 Supplementary figures 10 and 11.

### Visual Comparison of Phylogenetic Trees

To compare the trees obtained with different alignment tools, we used Phylo.io (*78*). The trees were imported in Newick format, and the Phylo.io tool generated the mirrored and aligned versions of the trees represented in Figure 2 Supplementary figures 8, 9, 12 and 13. The color of the branches is the tree similarity score, where lighter colors represent a higher number of topological differences. Exactly, it is a custom implementation of the Jacard Index by Phylo.io.

### Statistical Analysis of Tree Topology

To statistically evaluate our best-supported topology of the OSK and LOTUS trees, we compared constrained topologies to the highest likelihood trees using the SOWHAT tool (*31*). SOWHAT automates the stringent SOWH phylogenetic topology test (*30*), and compares the log likelihood between generated trees. We defined three constrained trees to test our results, one requiring monophyly of all domains of life, a second requiring only eukaryotic monophyly, and the last one requiring monophyly of the oskar LOTUS domain (Supplementary Files: Data>Trees>constrained_kingdom_tree.tre, constrained_eukmono_tree.tre & constrained_lotus_mono_tree.tre). We then ran SOWHAT using its default parameters, 1000 bootstraps, and the two constrained trees against the OSK or LOTUS alignment used to generate the phylogenetic trees (Supplementary Files: Alignments>OSK_MUSCLE_TREE.fasta & LOTUS_MUSCLE_TREE.fasta). All best trees generated by SOWHAT are available in (Supplementary Files: Data>Trees>SOWHAT_*_test.tre).

### Selection of Sequences for Codon Use Analysis

To study the codon use of the OSK and LOTUS domains, we chose 17 well-annotated (defined as possessing at least 8,000 annotated genes) insect genomes that included a confidently annotated *oskar* orthologue from the NCBI nucleotide database. The complete list and accession numbers of the sequences used for this analysis is in Supplementary Table 5. This list contains *oskar* sequences from genomes that were either added to the databases after the first *oskar* sequence search or re-annotated after said search. Therefore the sequences coming from the following organisms are not represented in the final *oskar* alignment: *Harpegnathos saltator, Fopius arisanus, Athalia rosae, Orussus abietinus, Stomoxys calcitrans, Bactrocera oleae, Neodiprion lecontei*.

### Calculation of Codon Adaptation Index (CAI)

For each of the 17 genomes indicated in Table SX, a codon use index was calculated by measuring the Relative Synonymous Codon Use (RCSU) of all genes in each genome. Using this codon use index, the CAI for all genes, the 3’ and 5’ cuts of the intra-gene distribution, Oskar, OSK and LOTUS was calculated for each genome. All calculations were performed using the Biopython library (https://biopython.org/ version 1.70). The Delta CAI was computed by taking the CAI difference between the 5’ and 3’ or LOTUS and OSK. The Z score of the intra-gene and LOTUS OSK were computed against the distribution of CAI of all genes in a genome.

### Generation of Intra-Gene Distribution of Ccodon Use

We wished to determine whether *oskar* differed from the null hypothesis that a given gene would follow similar codon use throughout its sequence. To generate a distribution of codon use similarity across a gene for all genes in the genomes studied, we generated what we named the “Intra-Gene” sequence distribution (Figure 3 Supplement 5). Each gene was cut into two fragments at a random position “x” following the rule: 351 < x < Length_gene - 351, x modulo 3 = 0 (Corresponding Jupyter notebook file: Scripts>notebook>Codon Analysis AT3 GC3 and A3 T3 G3 C3 Section: 4). In other words, for this analysis we split each transcript into two parts, requiring that each part be a minimum of 351 bp (117 codons) long. This approach allowed us to sample at least 117 codons for each of the 3’ and 5’ parts of a transcript.

### Fitting a Linear Model of Codon Use

Using the Intra-Gene null distribution generated above, we fitted a linear model of codon use frequencies per gene for the wobble position and AT3 GC3 content. To do so, we measured the different frequencies of A3, T3, G3 and C3 (any codon ending in A was counted as A3) and AT3 GC3. Then, we fitted a linear model to the pairs of 5’ and 3’ regional codon use values for within each gene, obtained from the Intra-Gene distribution described above (conserving the 3’/5’ position information), and for the OSK and LOTUS domains, for each of the 17 genomes analyzed (Supp Table 3). We then calculated the residuals of the Intra-Gene distribution and the LOTUS-OSK distribution. Finally, we determined the Pearson correlation coefficient for all genomes pooled together, and all *oskar* genes pooled together (Corresponding Jupyter notebook file: Scripts>notebook>Codon Analysis AT3 GC3 and A3 T3 G3 C3 Section: 7 and 8).

### Calculation and Analysis of the Codon Use Z_score

For each genome, the codon use frequency for AT3/GC3 and A3/T3/G3/C3 was calculated as described above. Then, Z scores for each sequence from the Intra-Gene, OSK or LOTUS domain sequences were calculated against the corresponding genome frequency distribution. The Z scores were then used to generate the analysis of Pearson correlation coefficients shown in Figures 3, Figure 3–figure supplement 1 and 2 (Corresponding Jupyter notebook file: Scripts>notebook>Codon Analysis AT3 GC3 and A3 T3 G3 C3 Section: 3, 5 and 6).

### Data Availability

All sequences discovered using the automatic annotation pipeline described in (M&M HMM and oskar search) are annotated as such in Supplementary Table S2.

### Code Availability

All custom code generated for this study is available in the GitHub repository https://github.com/extavourlab/Oskar_HGT, commit ID 2565e2f313db9eb310f907ef37ab644985fba565.

## Supporting information

Oskar HGT Supplementary Information Files

## Acknowledgments

We thank Sean Eddy, Chuck Davis, and Extavour lab members for discussion.

## Author contributions

CGME conceived of the project and overall experimental design. TEMJ collected initial transcriptome datasets and identified *oskar* orthologues therein. LB built the HMM model, identified additional orthologues, and performed sequence, phylogenetic, cluster and codon use analyses. LB and CGME interpreted data and wrote the manuscript.

## Data and materials availability

All data is available in the main text or the supplementary materials.

## Supplementary Information

The Supplementary Information for this paper consists of the following elements:

**BLondel_Jones_Extavour_Hgt_Supplementary_Materials_ALL *(this PDF document)***

1. Supplementary Tables

a. Table S1: List of genomes and transcriptomes used for automated *oskar* search.
b. Table S2: List of *oskar* sequences used in the final alignment.
c. Table S3: List of sequences used for phylogenetic analysis of the LOTUS domain.
d. Table S4: List of sequences used for phylogenetic analysis of the OSK domain.
e. Table S5: List of genomes analyzed for codon use.

All scripts used for this analysis are hosted on GitHub at https://github.com/extavourlab/Oskar_HGT

**Oskar HGT Supplementary Information Files *(FOLDER)***

1. Subfolder **Alignments**: All sequences identified and analyzed in this study, in FASTA format and with corresponding Alignments
2. Subfolder **BLAST search results**: Results of BLASTP searches with full length Oskar, OSK or LOTUS domains as queries
3. Subfolder **Data**: Necessary files for running the different IPython notebooks:

a. Subfolder ***HMM***: HMM models used for iterative searching for sequences similar to full-length Oskar, LOTUS and OSK domains
b. Subfolder ***Taxonomy***: Conversion table for UniProt ID to taxon information. (uniprot_ID_taxa.tsv)
c. Subfolder ***Trees***: Contains the tree files obtained from

i. RaxML phylogenetic analyses of the OSK and LOTUS domains aligned with MUSCLE, T-Coffee or PRANK
ii. MrBayes phylogenetic analyses of the OSK and LOTUS domains aligned with MUSCLE
iii. SOWHAT analyses.

Please download Oskar HGT SUPPLEMENTARY INFORMATION FILES Folder here: https://www.dropbox.com/sh/zqf6kpo0kzav7xp/AAC5WPVm9lrDrZHqZg_RlsiTa?dl=0

## Supplementary Figure Legends

### Figure 2 Supplements

**Figure 2–figure supplement 1:**
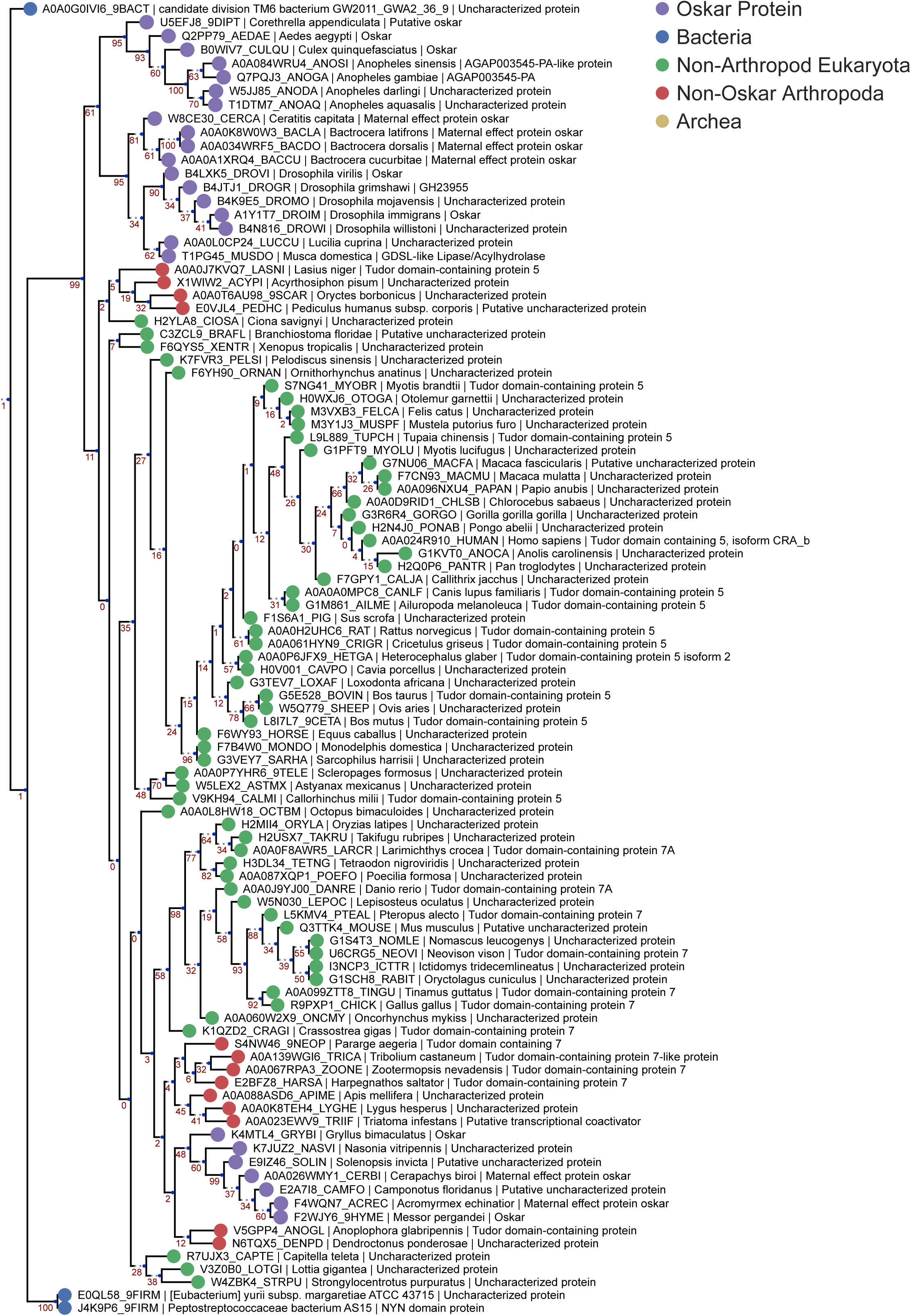
LOTUS Domain RaxML MUSCLE Tree. Phylogenetic tree of the HMMER sequences retrieved from the UniProt database using the LOTUS alignment HMM model. The top 97 hits were selected for phylogenetic analysis, and the only three bacterial sequences found to be a match were added to the alignment manually. The resulting 100 sequences were aligned using MUSCLE with default settings. The sequences were filtered to contain only one sequence per species (best E-value kept) yielding 100 sequences for analysis. Finally, the tree was created using RaxML v8.2.4, using 1000 bootstraps and model selection performed by the RaxML automatic model selection tool. See “Phylogenetic Analysis” in Methods for further detail. Sequences are color-coded as follows: Purple = Oskar; Red = Non-Oskar Arthropod; Green = Non-Arthropod Eukaryote; Blue = Bacteria. Names following leaves display the UniProt accession number followed by the species name and the UniProt protein name.

**Figure 2–figure supplement 2:**
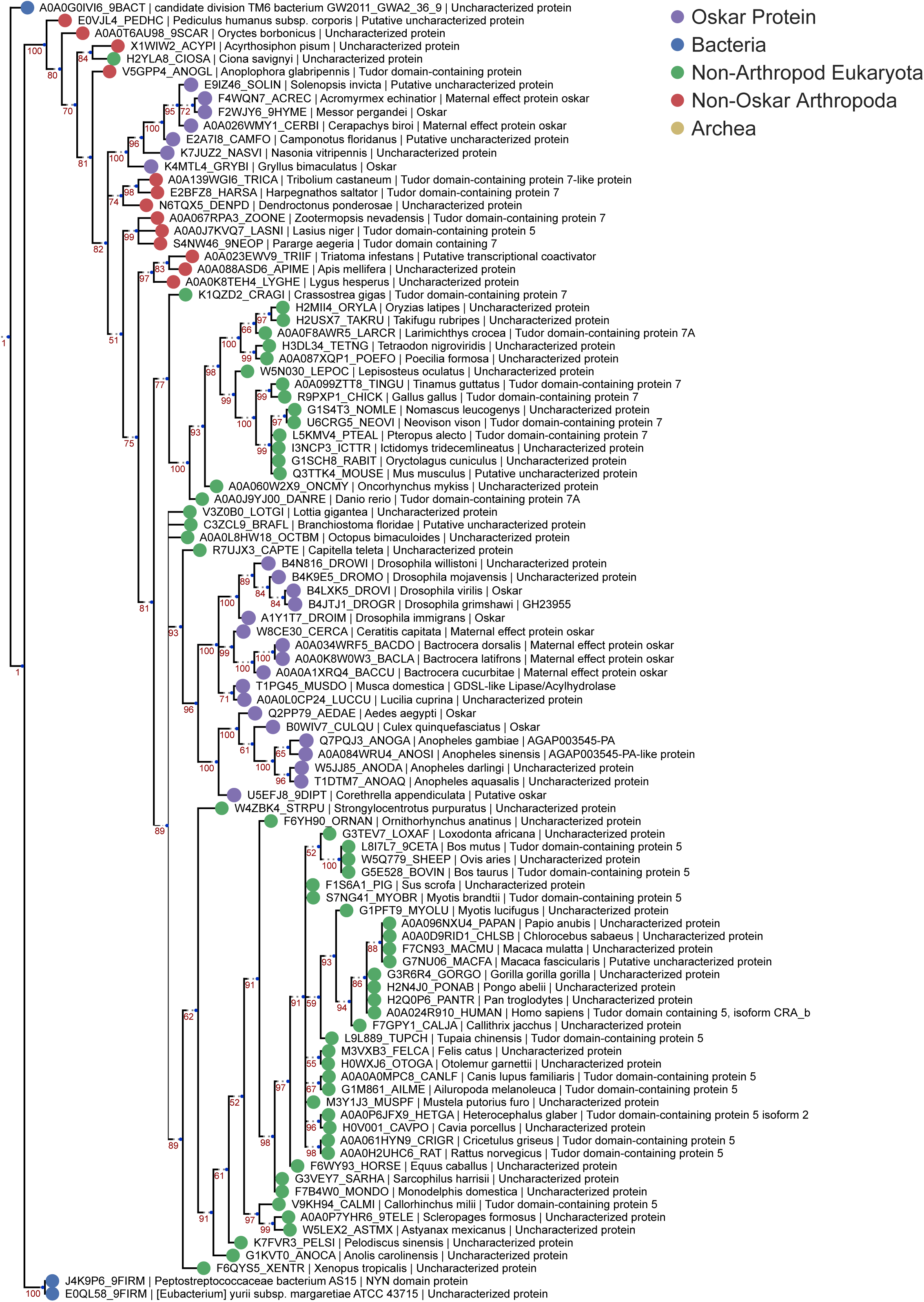
LOTUS Domain Bayesian MUSCLE Tree. Phylogenetic tree of the HMMER sequences retrieved from the UniProt database using the LOTUS alignment HMM model. 100 sequences were chosen for analysis as described for Supplementary Figure 1. The tree was created using Mr Bayes V3.2.6 using a Mixed model (prset aamodel=Mixed) and a gamma distribution (lset rates=Gamma). The algorithm was allowed to run for 3 million generations to achieve a std < 0.01. See “Phylogenetic Analysis” in Methods for further detail. Sequences are color-coded as follows: Purple = Oskar; Red = Non-Oskar Arthropod; Green = Non-Arthropod Eukaryote; Blue = Bacteria. Names following leaves display the UniProt accession number followed by the species name and the UniProt protein name.

**Figure 2–figure supplement 3:**
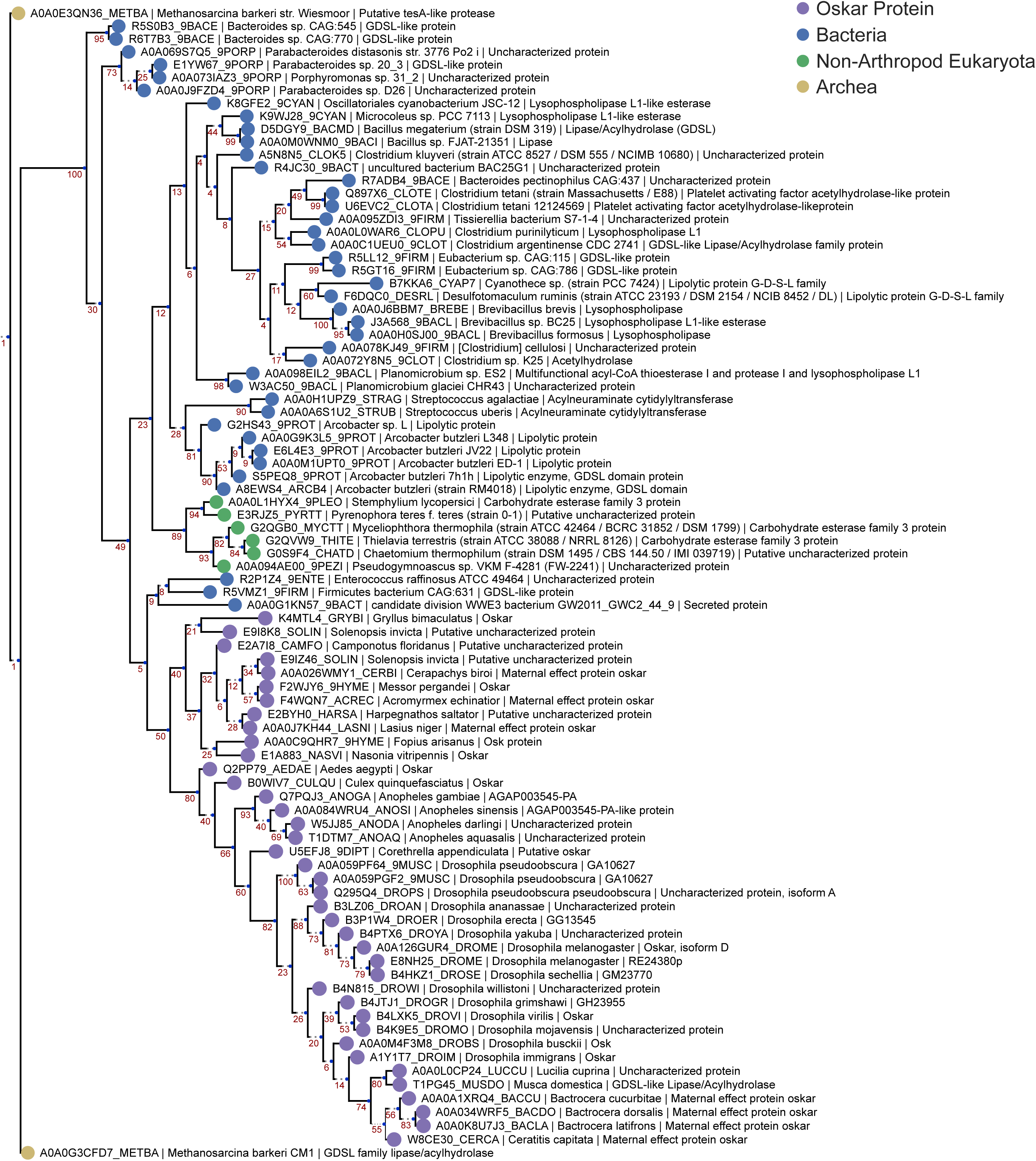
OSK Domain RaxML MUSCLE Tree. Phylogenetic tree of the HMMER sequences retrieved from the UniProt database using the OSK alignment HMM model. The top 95 hits were selected for phylogenetic analysis, and the only five non-Oskar eukaryotic sequences found to be a match were added to the alignment manually. The resulting 100 sequences were aligned using MUSCLE with default settings. The sequences were filtered to contain only one sequence per species (best E-value kept), yielding 87 sequences for analysis. Finally, the tree was created using RaxML v8.2.4, using 1000 bootstraps and model selection performed by the RaxML automatic model selection tool. See “Phylogenetic Analysis” in Methods for further detail. Sequences are color-coded as follows: Purple = Oskar; Red = Non-Oskar Arthropod; Green = Non-Arthropod Eukaryote; Blue = Bacteria. Names following leaves display the UniProt accession number followed by the species name and the UniProt protein name.

**Figure 2–figure supplement 4:**
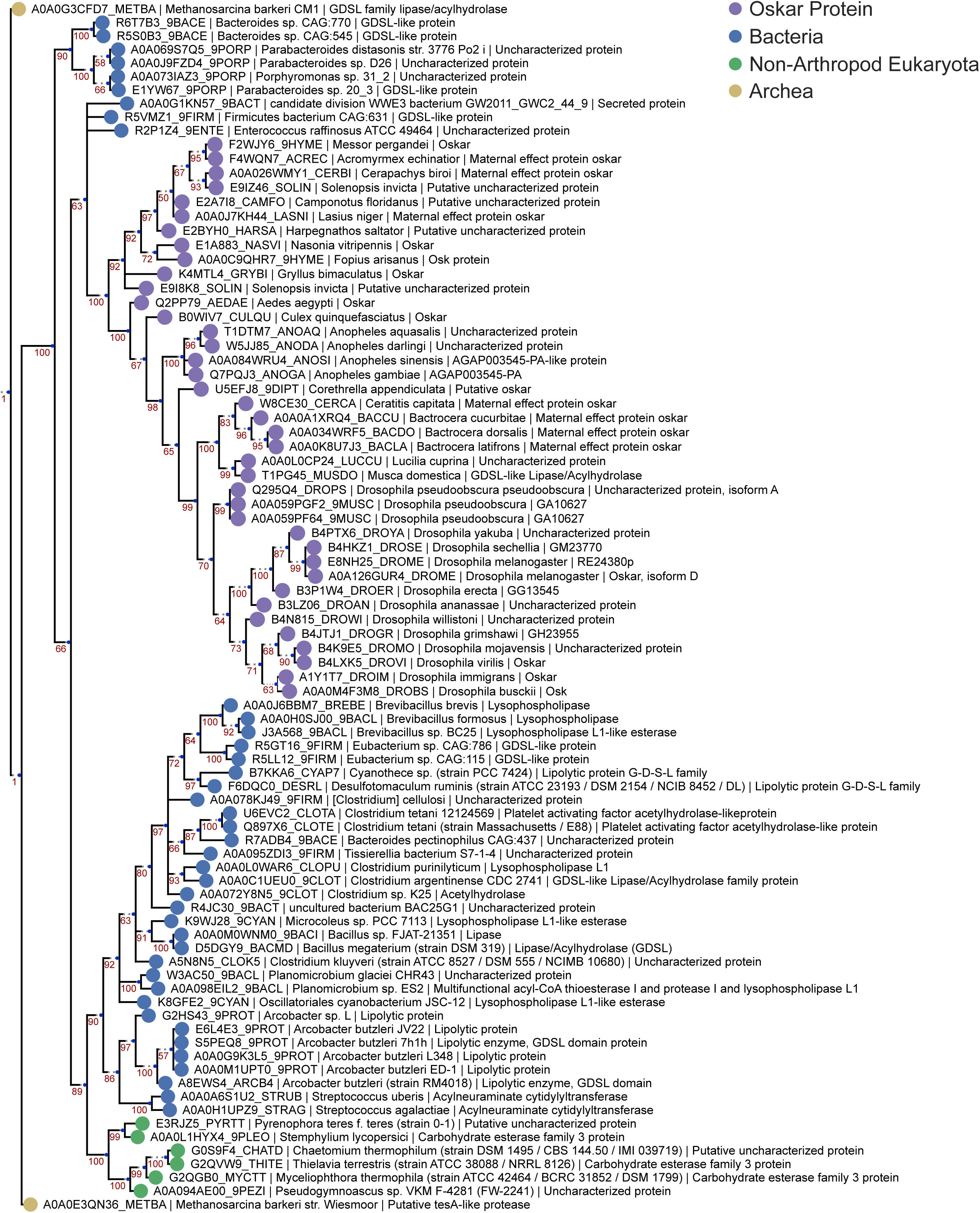
OSK Domain Bayesian MUSCLE Tree. Phylogenetic tree of the HMMER sequences hit on the UniProt database using the OSK alignment HMM model. 87 sequences were chosen for analysis as described for Supplementary Figure 3. The tree was created using Mr Bayes V3.2.6 using a Mixed model (prset aamodel=Mixed) and a gamma distribution (lset rates=Gamma). The algorithm was allowed to run for 4 million generations to achieve a std < 0.01. See “Phylogenetic Analysis” in Methods for further detail. Sequences are color-coded as follows: Purple = Oskar; Red = Non-Oskar Arthropod; Green = Non-Arthropod Eukaryote; Blue = Bacteria. Names following leaves display the UniProt accession number followed by the species name and the UniProt protein name.

**Figure 2–figure supplement 5:**
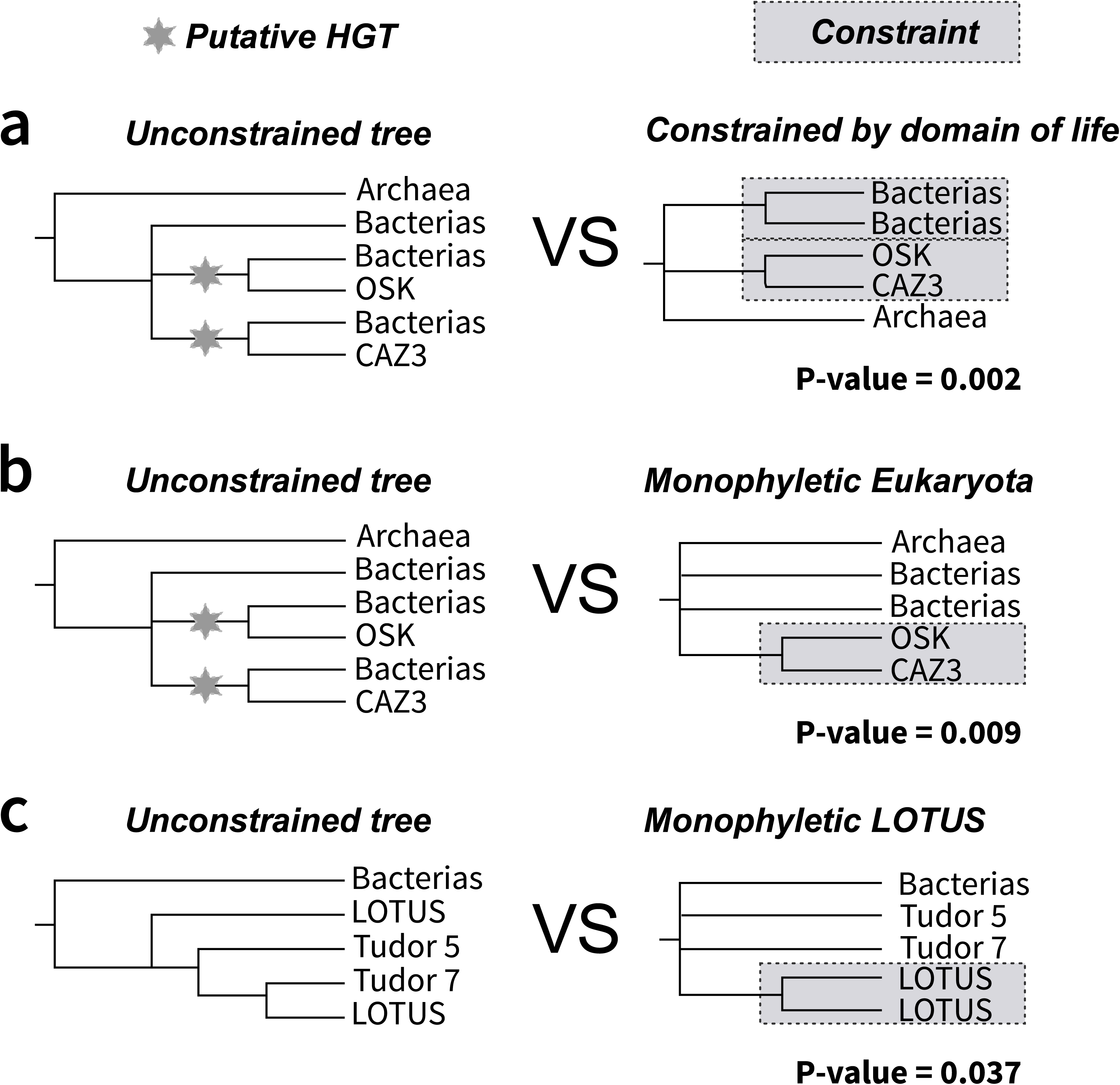
SOWHAT constrained trees and results. Two trees constrained by alternative relationships that would be expected under vertical transmission of sequences were designed and tested against our result supporting a putative HGT event of the OSK domain. (**a**) The first tree (right) is constrained by domain of life, requiring bacterial and eukaryotic sequences to be monophyletic, and disallowing sister group relationships of subsets of eukaryotic sequences and bacterial sequences. Our unconstrained tree topology (left) outperformed this topology with a p-value of 0.002 (95% confidence interval upper: 0.007 lower: 0.0002). (**b**) The second tree requires monophyly of Eukaryota. Our unconstrained tree topology (left) outperformed this topology with a p-value of 0.009 (95% confidence interval upper: 0.017 lower: 0.004). (**c**) The third tree tested whether the LOTUS domain split observed in the tree generated with the MUSCLE alignment was significantly different from a tree where the LOTUS sequences formed a monophyly. The unconstrained tree (left) outperformed this topology with a p-value of 0.037 (95% confidence interval upper: 0.05 lower: 0.026).

**Figure 2–figure supplement 6:**
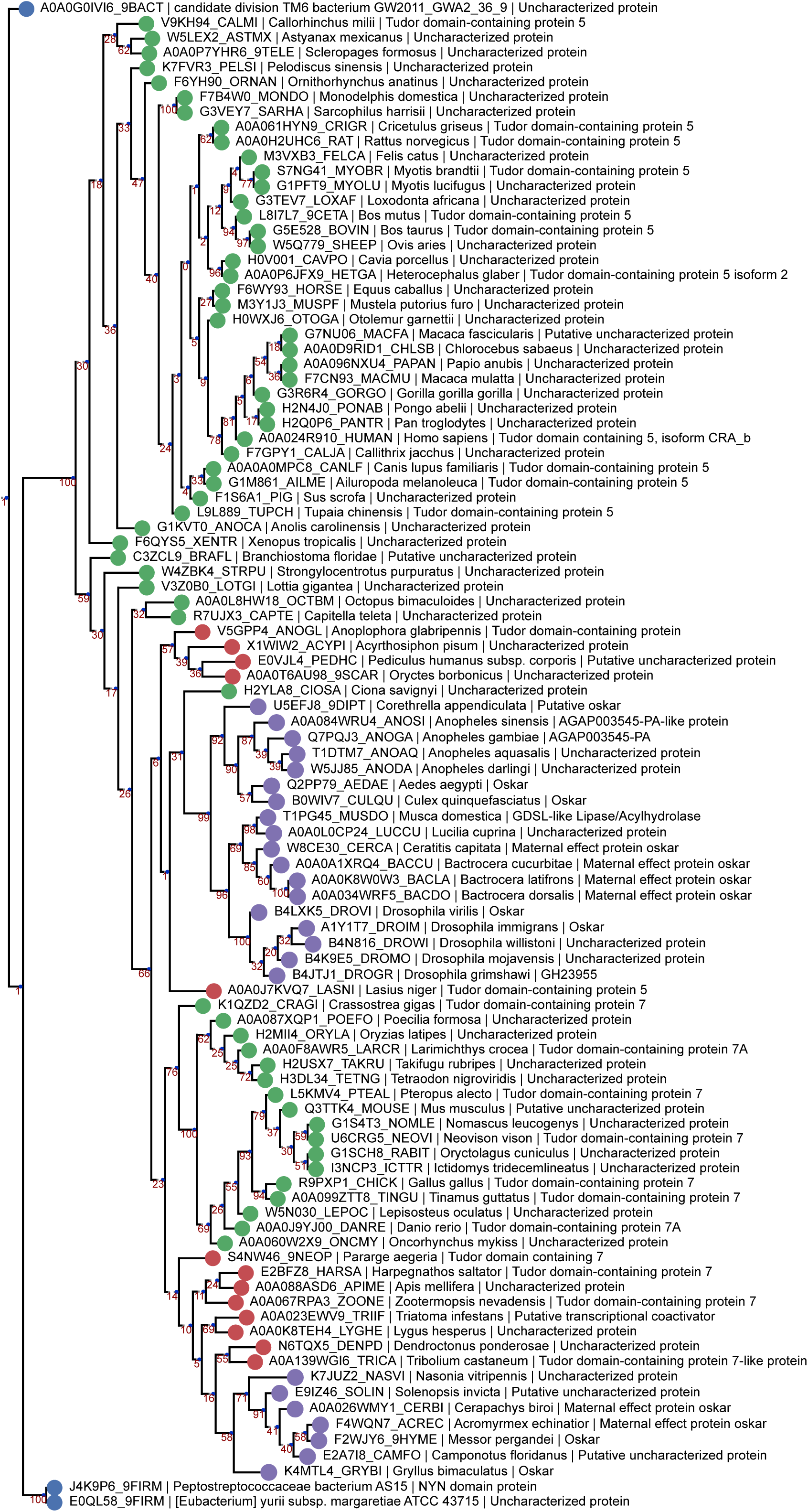
LOTUS Domain RaxML PRANK Tree. Phylogenetic tree of the same sequences used for the previous LOTUS trees. The sequences were aligned using PRANK and the tree generated with RaxML as described in ***Phylogenetic Analysis Based on PRANK alignment***. Sequences are color-coded as follows: Purple = Oskar; Red = Non-Oskar Arthropod; Green = Non-Arthropod Eukaryote; Blue = Bacteria. Names following leaves display the UniProt accession number followed by the species name and the UniProt protein name.

**Figure 2–figure supplement 7:**
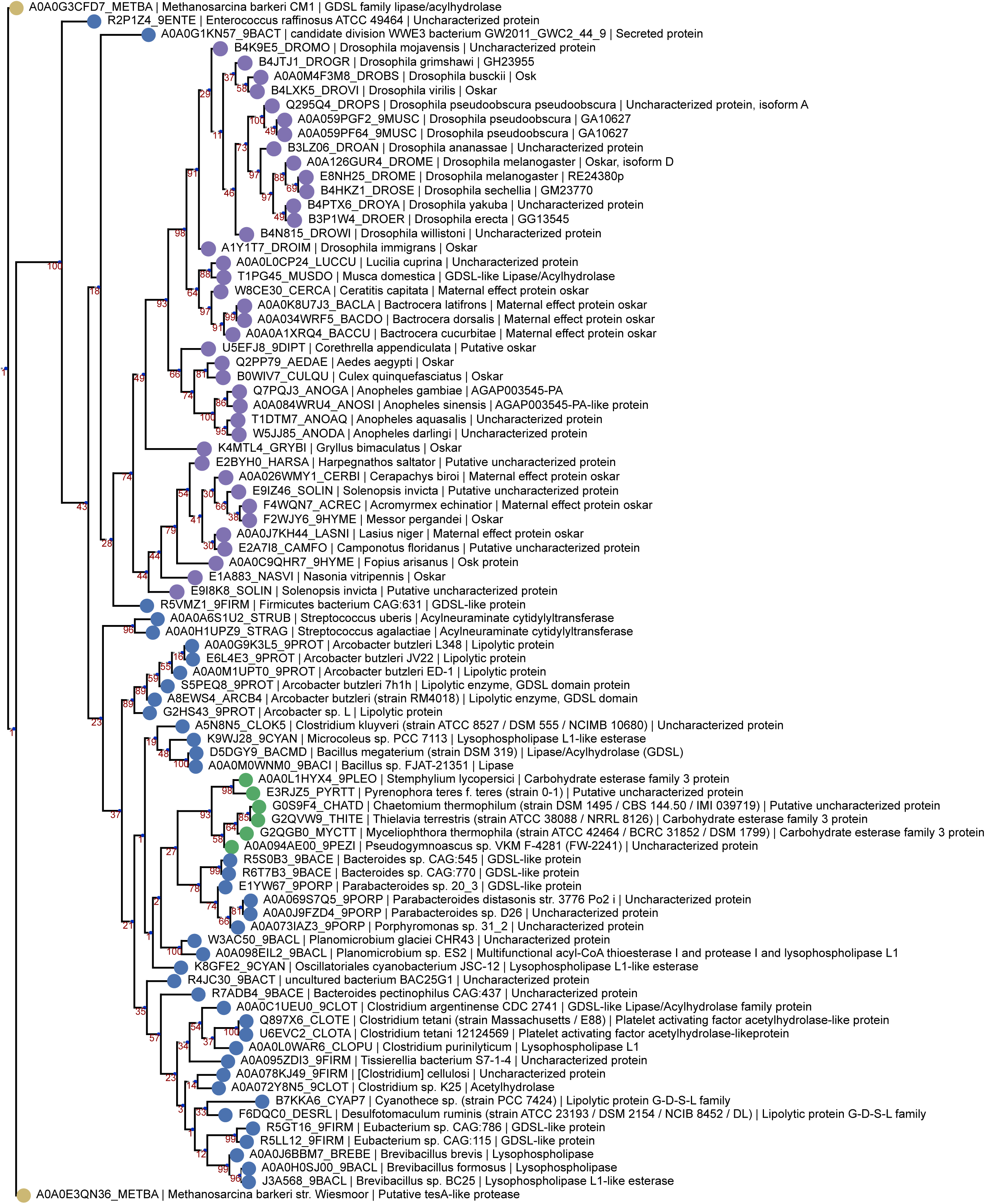
OSK Domain RaxML PRANK Tree. Phylogenetic tree of the same sequences used for the previous OSK trees. The sequences were aligned using PRANK and the tree generated with RaxML as described in ***Phylogenetic Analysis Based on PRANK alignment***. Sequences are color-coded as follows: Purple = Oskar; Red = Non-Oskar Arthropod; Green = Non-Arthropod Eukaryote; Blue = Bacteria. Names following leaves display the UniProt accession number followed by the species name and the UniProt protein name.

**Figure 2–figure supplement 8:**
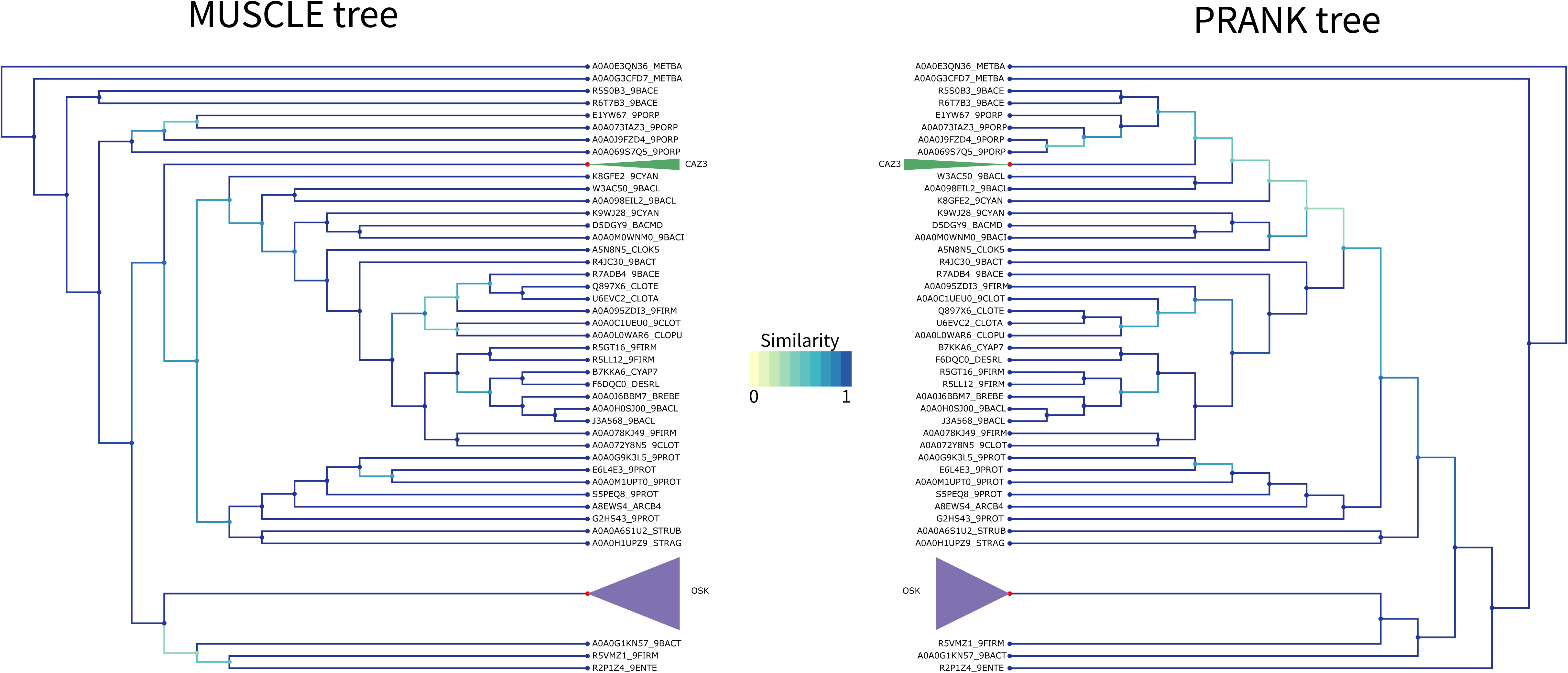
OSK Tree PRANK Comparison. Comparison of the tree obtained with RaxML starting from the MUSCLE alignment (left) versus the PRANK alignment (right) for the OSK domain. Similarity scores for the branching events are color coded from yellow to blue (see figure color bar legend). The OSK (purple) clade and CAZ3 (green) clade have been colored and compacted for readability as they do not have any internal branching changes. Node color is blue if the leaf is a sequence, and red if this is a compacted group of sequences.

**Figure 2–figure supplement 9:**
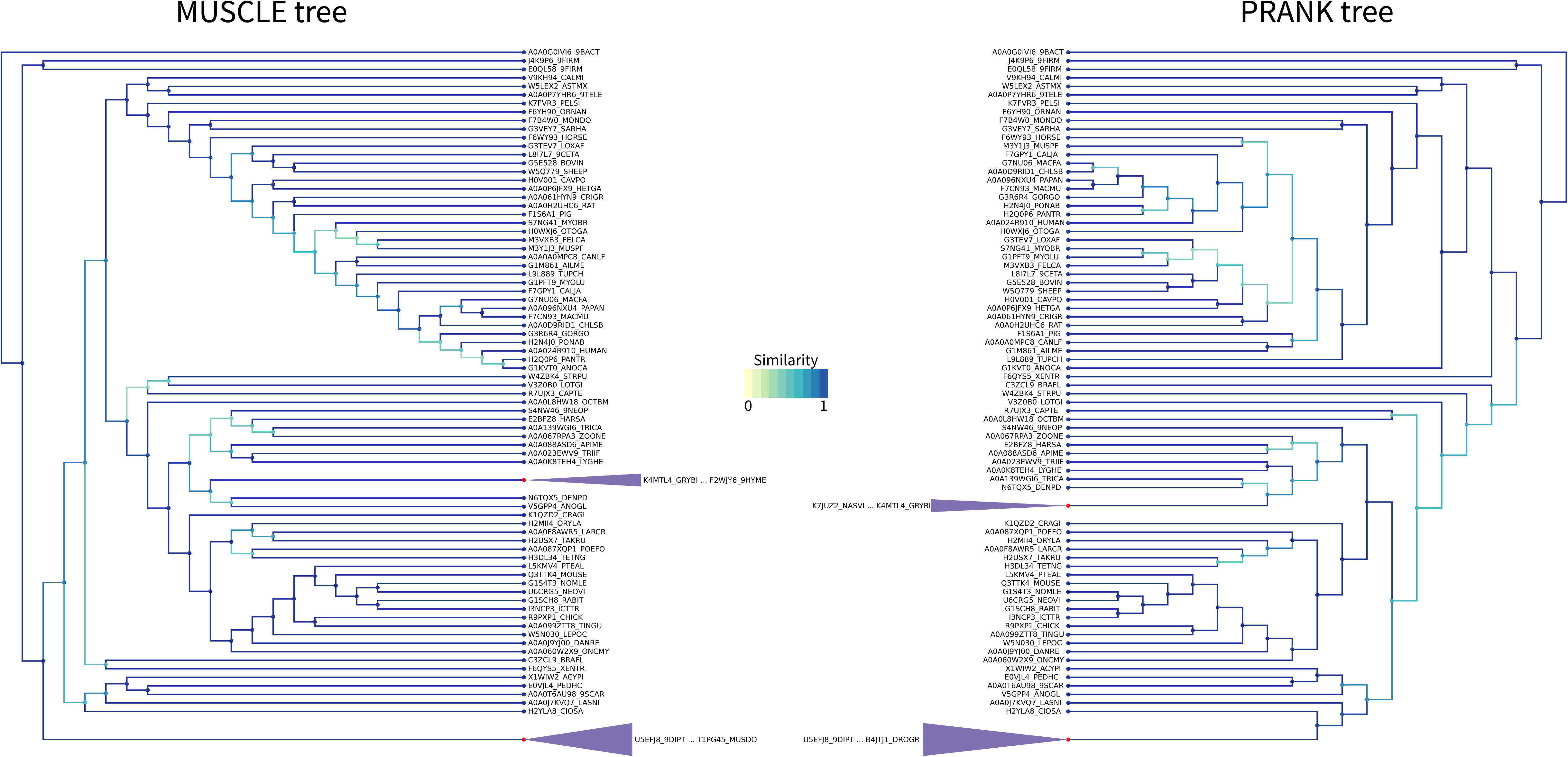
LOTUS Tree PRANK Comparison. Comparison of the tree obtained with RaxML starting from the MUSCLE alignment (left) versus the PRANK alignment (right) for the LOTUS domain. Similarity scores for the branching events are color coded from yellow to blue (see figure color bar legend). The LOTUS (purple) clades have been colored and compacted for readability as they do not have any internal branching changes. Node color is blue if the leaf is a sequence, and red if this is a compacted group of sequences.

**Figure 2–figure supplement 10:**
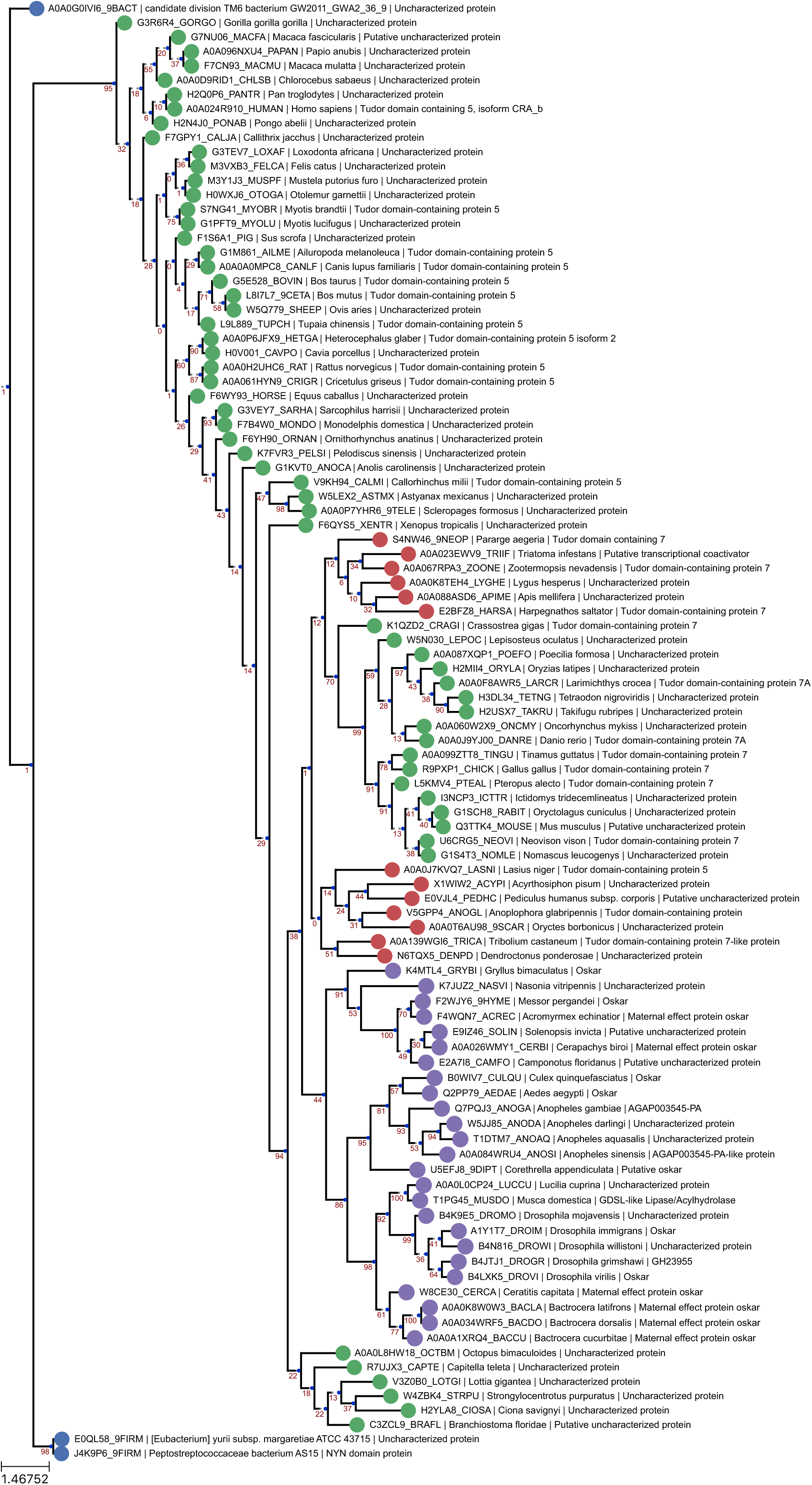
LOTUS Domain RaxML T-Coffee Tree. Phylogenetic tree of the same sequences used for the previous LOTUS trees. The sequences were aligned using T-Coffee and the tree generated with RaxML as described in ***Phylogenetic Analysis Based on* T-Coffee *alignment***. Sequences are color-coded as follows: Purple = Oskar; Red = Non-Oskar Arthropod; Green = Non-Arthropod Eukaryote; Blue = Bacteria. Names following leaves display the UniProt accession number followed by the species name and the UniProt protein name.

**Figure 2–figure supplement 11:**
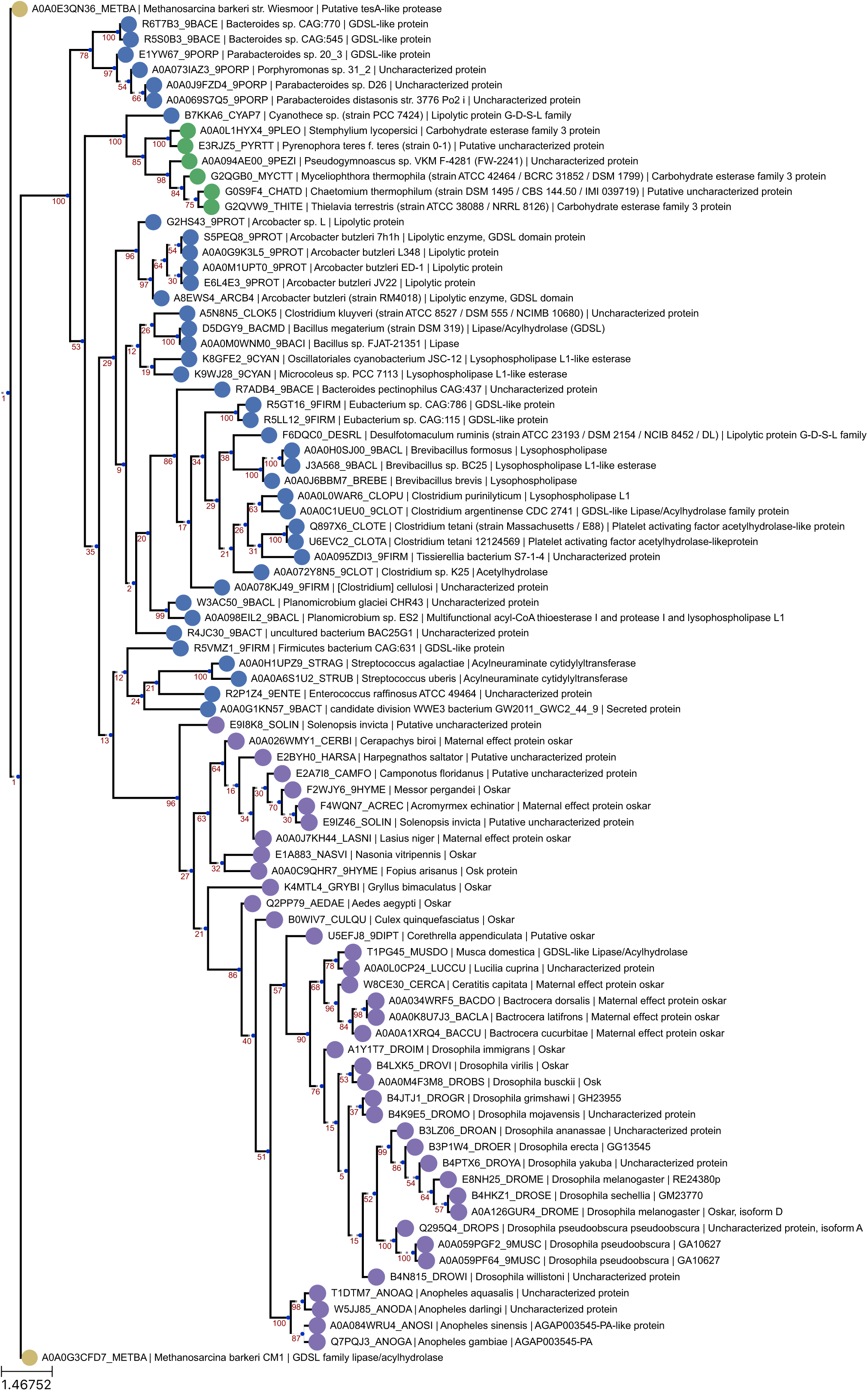
OSK Domain RaxML T-Coffee Tree. Phylogenetic tree of the same sequences used for the previous OSK trees. The sequences were aligned using T-Coffee and the tree generated with RaxML as described in ***Phylogenetic Analysis Based on* T-Coffee *alignment***. Sequences are color-coded as follows: Purple = Oskar; Red = Non-Oskar Arthropod; Green = Non-Arthropod Eukaryote; Blue = Bacteria. Names following leaves display the UniProt accession number followed by the species name and the UniProt protein name.

**Figure 2–figure supplement 12:**
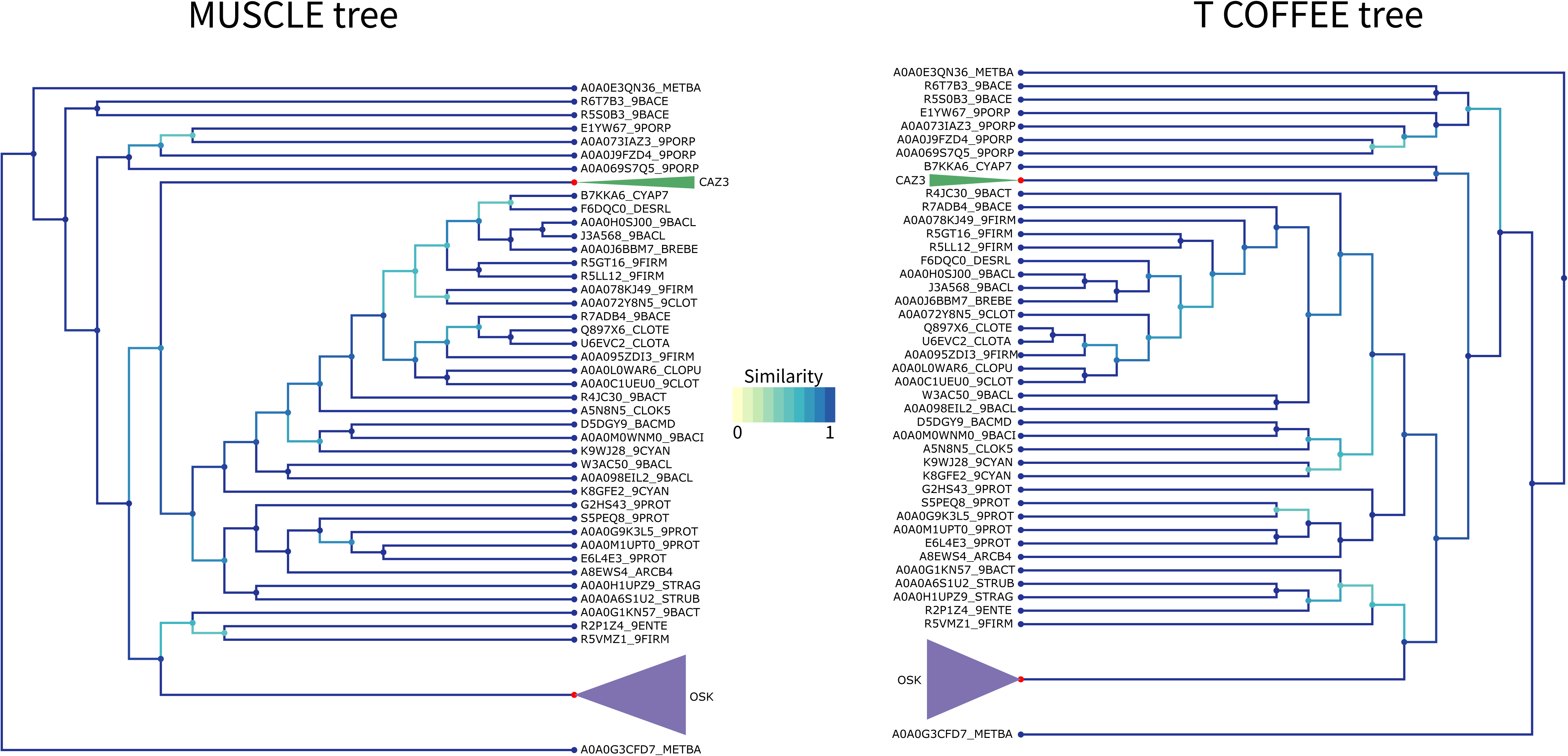
OSK Tree T-Coffee Comparison. Comparison of the tree obtained with RaxML starting from the MUSCLE alignment (left) versus the T-Coffee alignment (right) for the OSK domain. Similarity scores for the branching events are color coded from yellow to blue (see figure color bar legend). The OSK (purple) clade and CAZ3 (green) clade have been colored and compacted for readability as they do not have any internal branching changes. Node color is blue if the leaf is a sequence, and red if this is a compacted group of sequences.

**Figure 2–figure supplement 13:**
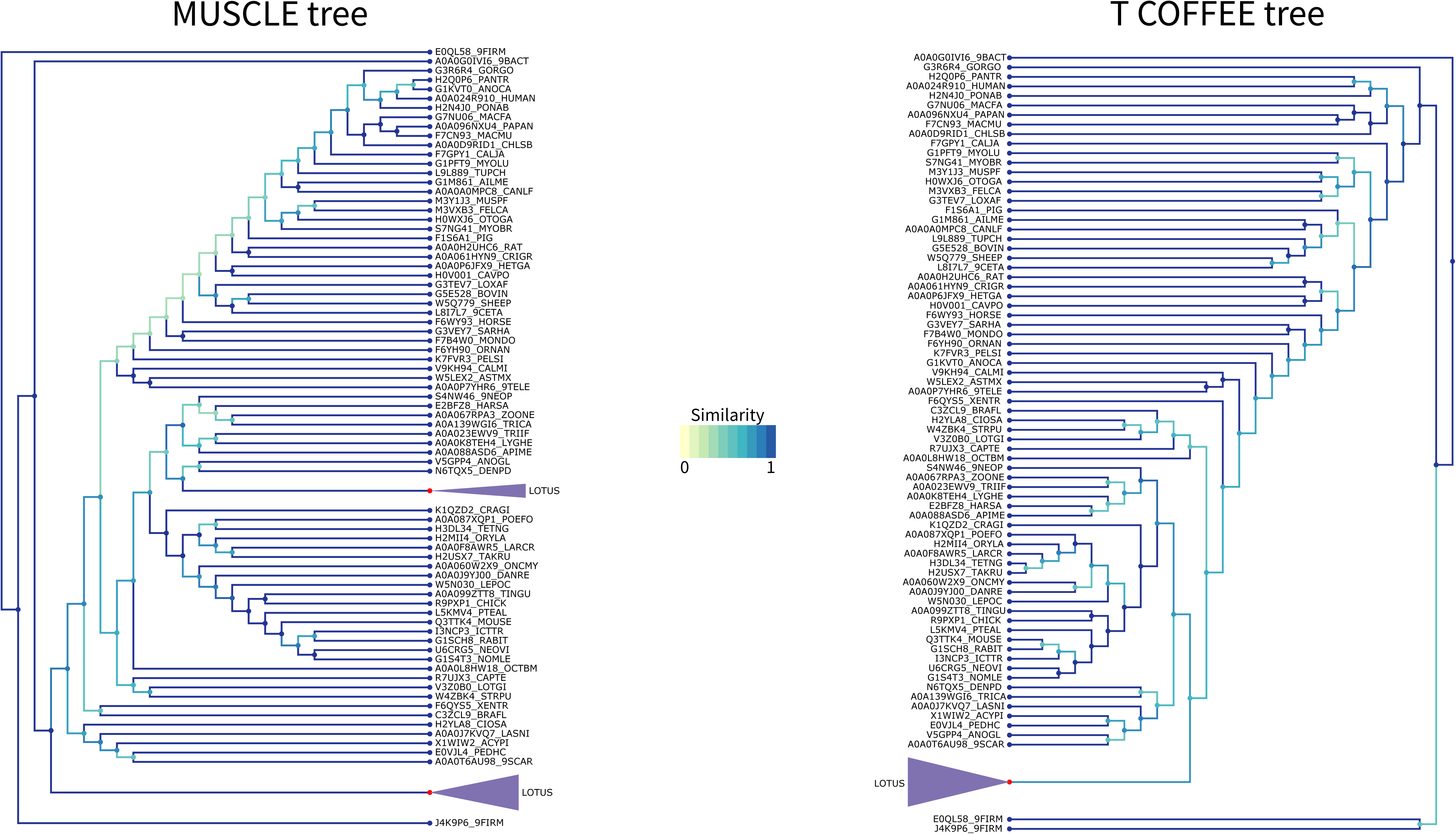
LOTUS Tree T-Coffee Comparison. Comparison of the tree obtained with RaxML starting from the MUSCLE alignment (left) versus the T-Coffee alignment (right) for the LOTUS domain. Similarity scores for the branching events are color coded from yellow to blue (see figure color bar legend). The LOTUS (purple) clades have been colored and compacted for readability as they do not have any internal branching changes. Node color is blue if the leaf is a sequence, and red if this is a compacted group of sequences.

### Figure 3 Supplements

**Figure 3–figure supplement 1:** GC3 correlations between the LOTUS and OSK domains. IntraGene distribution scatter plot for the coding sequences of the 17 genomes analyzed. Sequences were cut into two parts as per the description in Methods “Generation of IntraGene distribution of codon use”. Z-scores were calculated for the 5’ and 3’ ends against the genome distribution. Finally, the 5’ and 3’ “domain” values were plotted against each other and a linear regression was performed. The Oskar sequence was then plotted in red. A kernel density estimation is displayed as black contour lines, increasing in shades of blue as the density increases. ***Figure download link:*** https://www.dropbox.com/s/kxjypf064kano9u/Blondel_Jones_Extavour_HGT_Figure%203%E2%80%93figure%20supplement%201%3A%20GC3%20correlations%20between%20the%20LOTUS%20and%20OSK%20domains.pdf?dl=0

**Figure 3–figure supplement 2:** AT3 correlations between the LOTUS and OSK domains. Intra-Gene distribution scatter plot for the coding sequences of the 17 genomes analyzed. Sequences were cut into two parts as per the description in Methods “Generation of intra-gene distribution of codon use”. Z-scores were calculated for the 5’ and 3’ ends against the genome distribution. Finally, the 5’ and 3’ “domain” values were plotted against each other and a linear regression was performed. The Oskar sequence was then plotted in red. A kernel density estimation is displayed as black contour lines, increasing in shades of blue as the density increases. ***Figure download link:*** https://www.dropbox.com/s/1gt5xzu6eigikd5/Blondel_Jones_Extavour_HGT_Figure%203%E2%80%93figure%20supplement%202%3A%20AT3%20correlations%20between%20the%20LOTUS%20and%20OSK%20domains.pdf?dl=0

**Figure 3–figure supplement 3:**
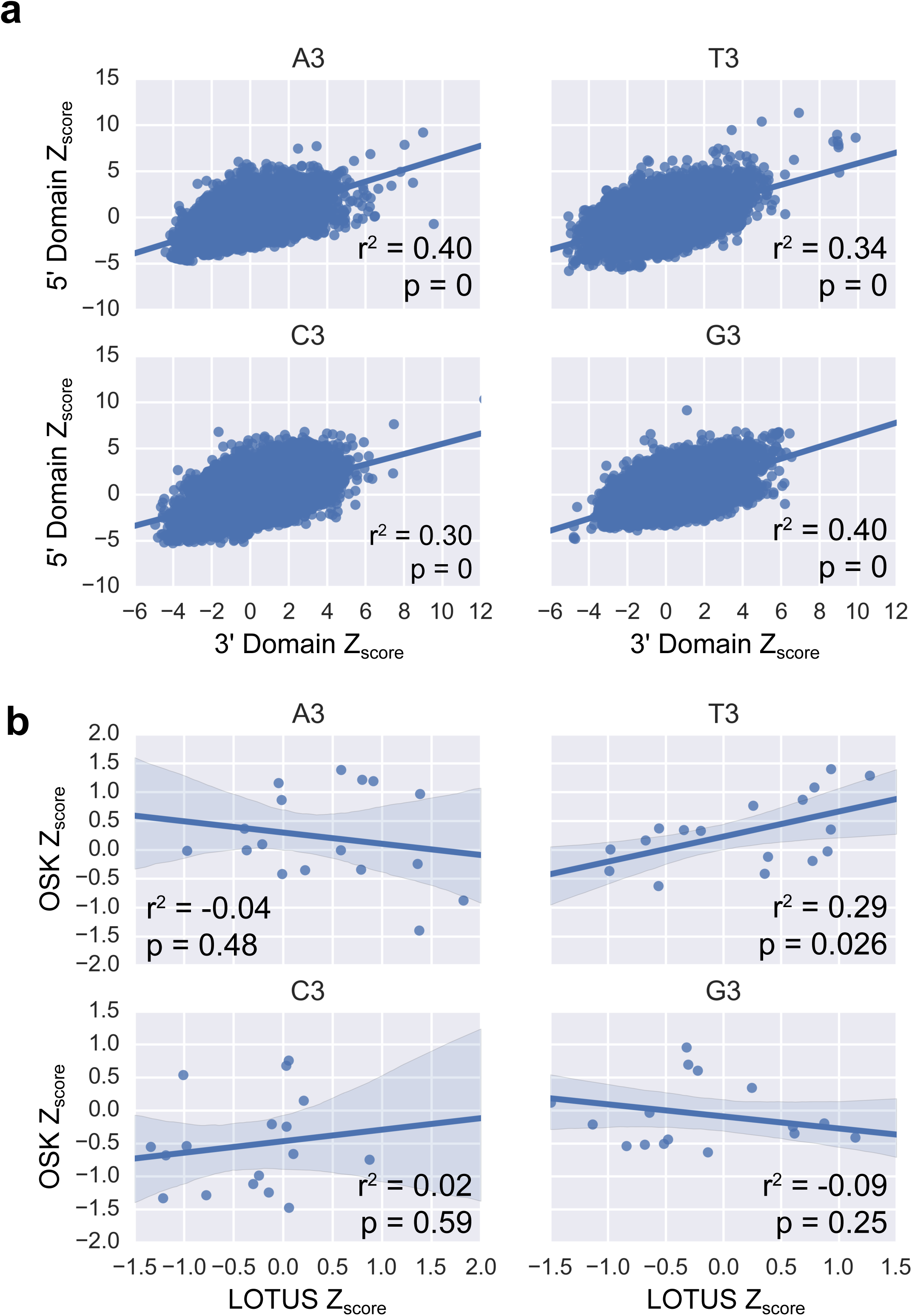
A3/T3/G3/C3 correlations between the LOTUS and OSK domains. (**a**) Intra-Gene distribution scatter plot for the coding sequences of the 17 genomes analyzed. Sequences were cut into two parts as per the description in Methods “Generation of intra-gene distribution of codon use”. The A3, T3, G3 and C3 codon use was measured, and Z-score calculations, value plots and linear regression were performed as described for Supplementary Figure 5. The A3, T3 G3 and C3 content is generally similar in the 5’ and 3’regions of all genes across the genome (A3: r^2^ = 0.40, p = 0; T3: r^2^ = 0.34, p = 0; G3: r^2^ = 0.40, p = 0; C3: r^2^ = 0.30, p = 0). (**b**) OSK vs LOTUS A3, T3, G3 and C3 use across the 17 genomes analyzed. The A3, T3, G3 and C3 content Z-score were calculated against the genome distribution. The A3, G3 and C3 content of the two domains of the Oskar gene are not correlated with each other. However, the T3 distribution follows a linear correlation similar to the one found across the Intra-Gene distribution (A3: r^2^ = −0.04, p = 0.48; T3: r^2^ = 0.29, p = 0.026; G3: r^2^ = −0.09, p = 0.25; C3: r^2^ = 0.02, p = 0.59).

**Figure 3–figure supplement 4:**
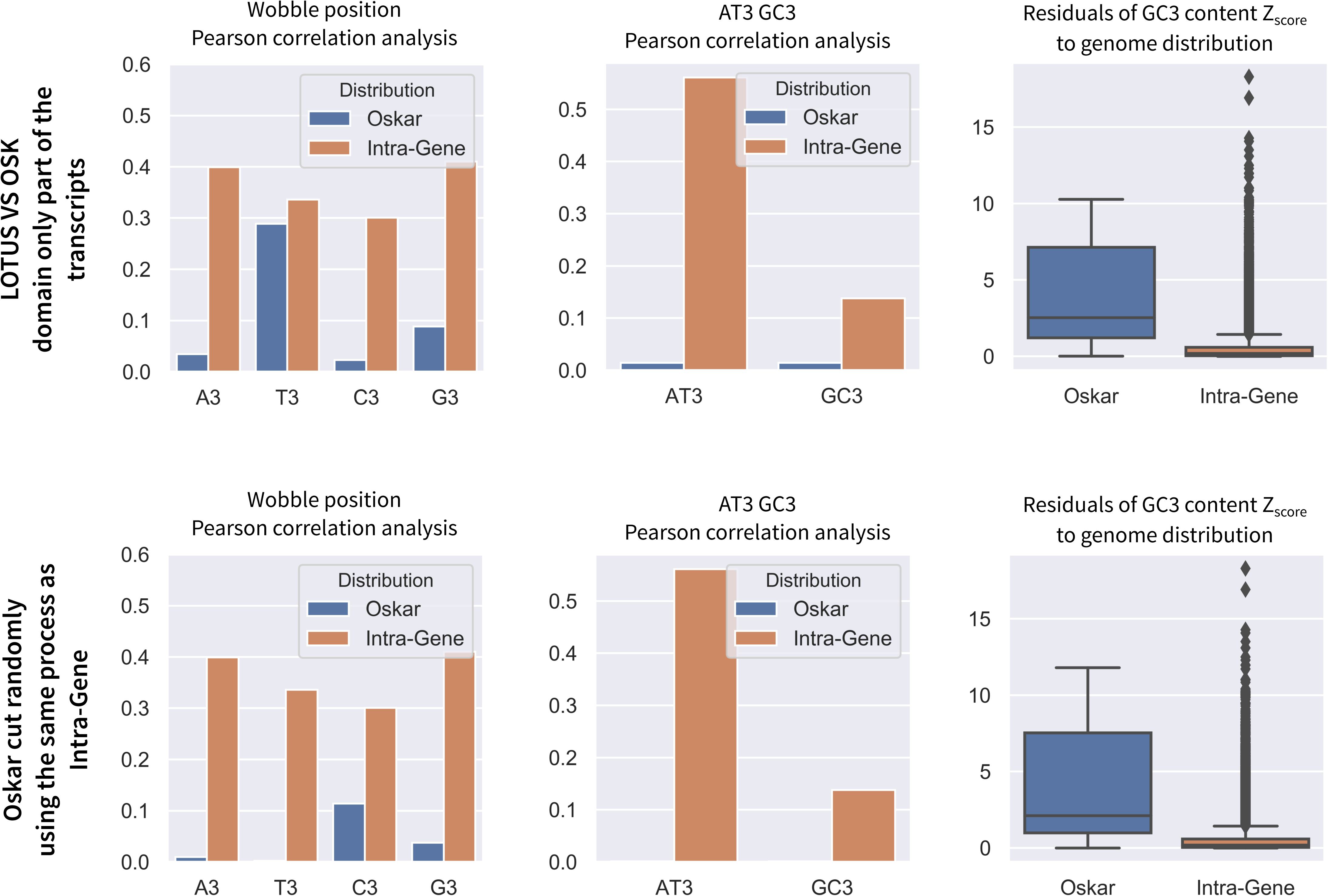
OSK-LOTUS VS Random Cut method. Comparison of Wobble position use of codon between OSK and LOTUS domains versus the Oskar gene cut at random following the same procedure as the intra-gene distribution. All analyses were performed as described in Figure 3.

**Figure 3–figure supplement 5:**
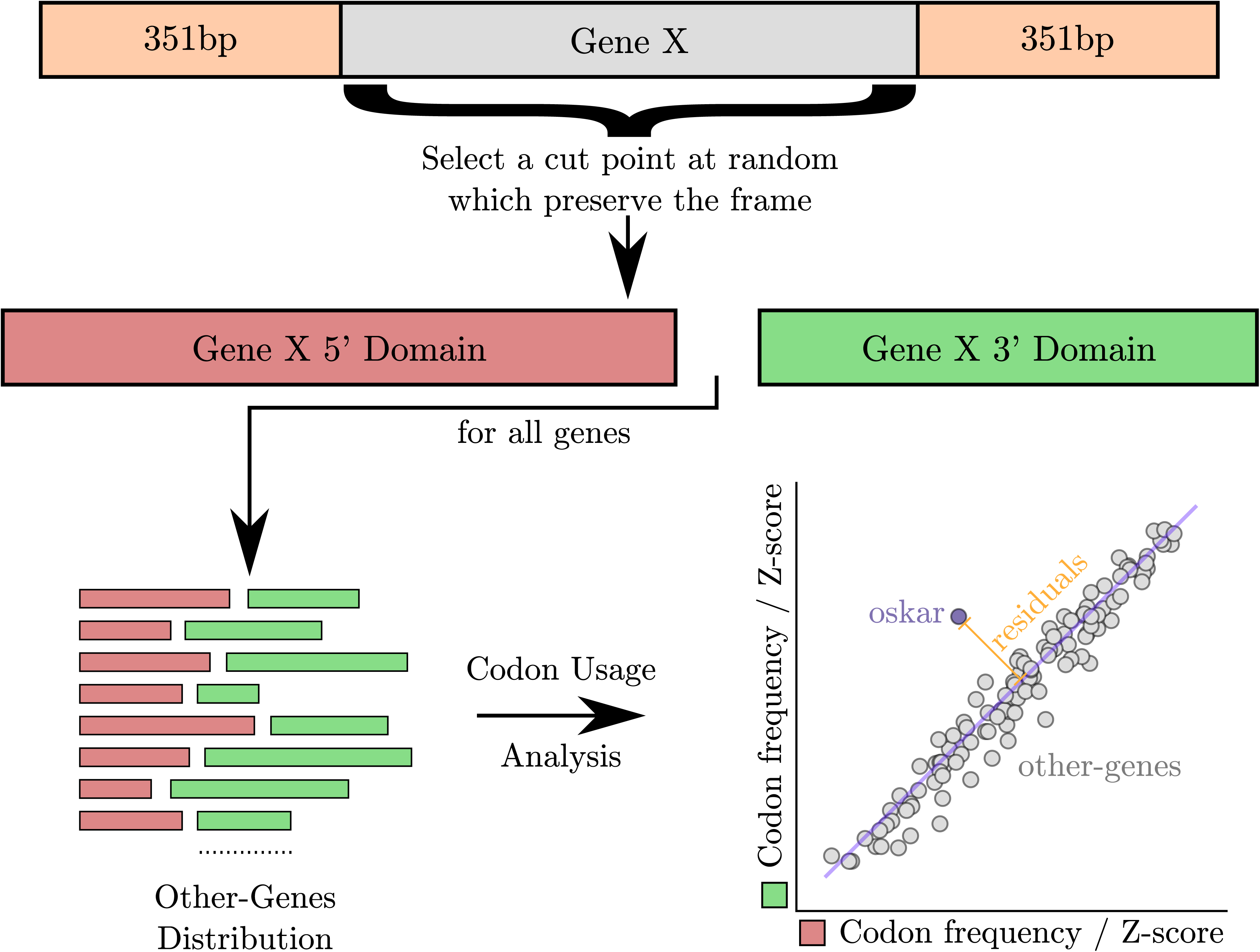
IntraGene Schematic. Schematic representation of the intra-gene distribution. For each gene, a random cut preserving the frame was chosen. The position was forced to be over 351bp and less than the length of the sequence minus 351bp. Each part of the sequence was then split into two bins, one for the 5’ and one for the 3’. Moreover, the gene identity was preserved to compare the 5’ and 3’ end of the same sequence in all following analysis. For more information see *Generation of Intra-Gene distribution of codon use*.

**Figure 3–figure supplement 6:**
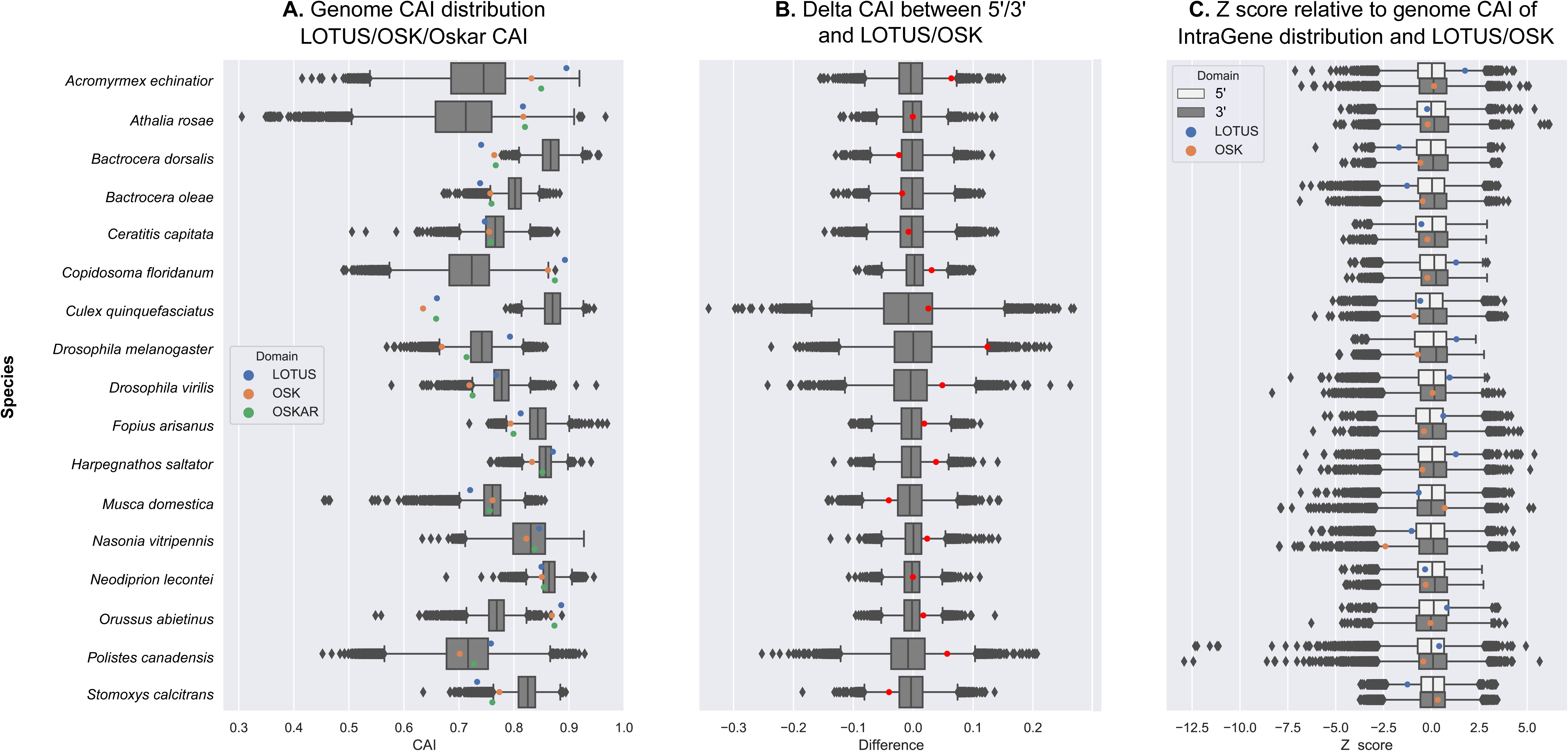
Genome and Oskar CAI. This Figure presents the results of the Codon Adaptation Index (CAI) analysis of *oskar*. The CAI were computed against a codon index measured for each genome (see *Calculation of Codon Adaptation Index (CAI)*). (A) The distribution of CAI for each gene in the genome (grey box whisker plot). Colored dots represent are the CAI values for full length Oskar (green), OSK domain (orange) and LOTUS domain (blue) sequences. (B) The distribution of the Delta CAI, or the difference in CAI between two halves of the same gene sequence. Grey = intra-gene distribution, where for each gene, the 5’ and 3’ CAI were computed and the difference calculated. Red = Delta CAI between the LOTUS and OSK domains of Oskar (C). The Z score distribution of the CAI scores for the 3’ (dark grey) and 5’ (light grey) halves of a gene. Colored dots represent the CAI scores for the LOTUS domain (blue) and OSK domain (orange) of Oskar.

**Supplementary Table S1:**
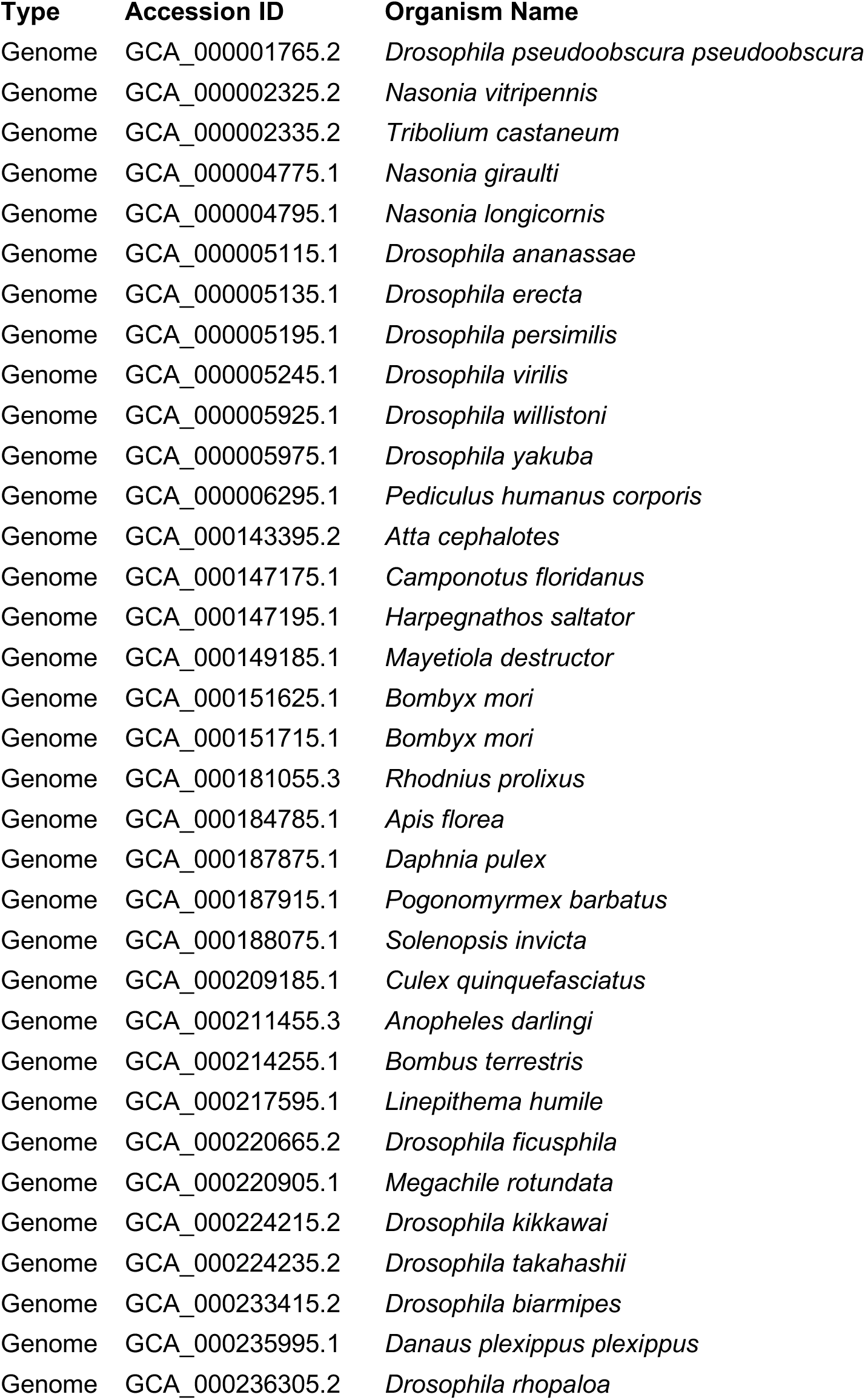
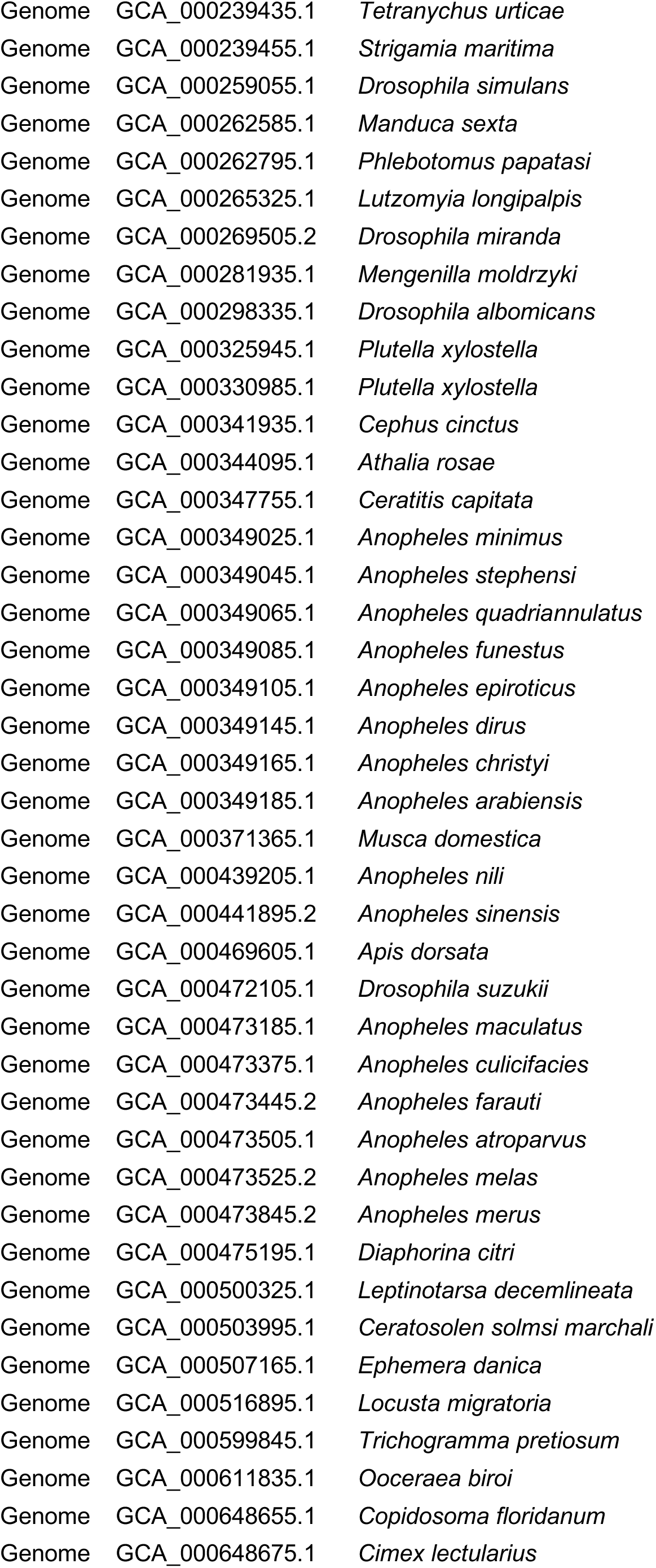
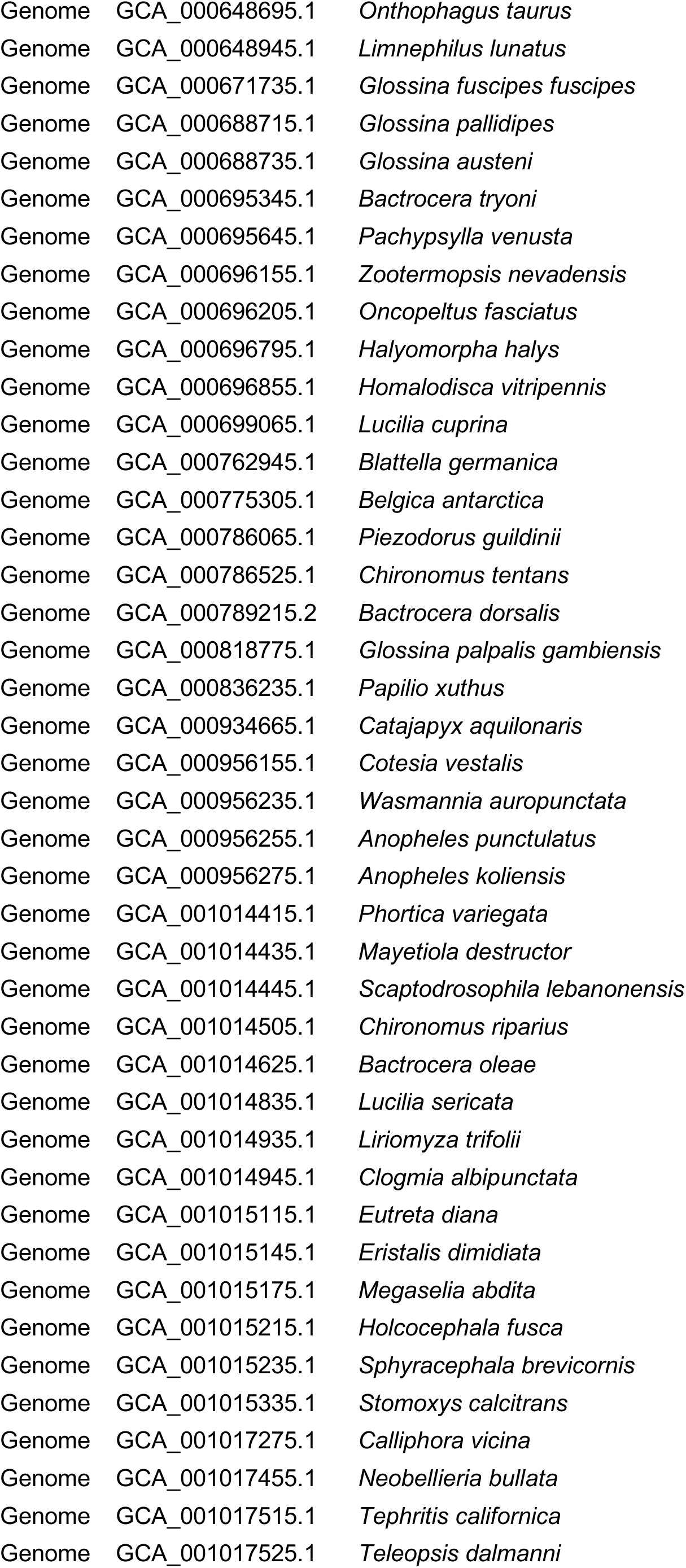
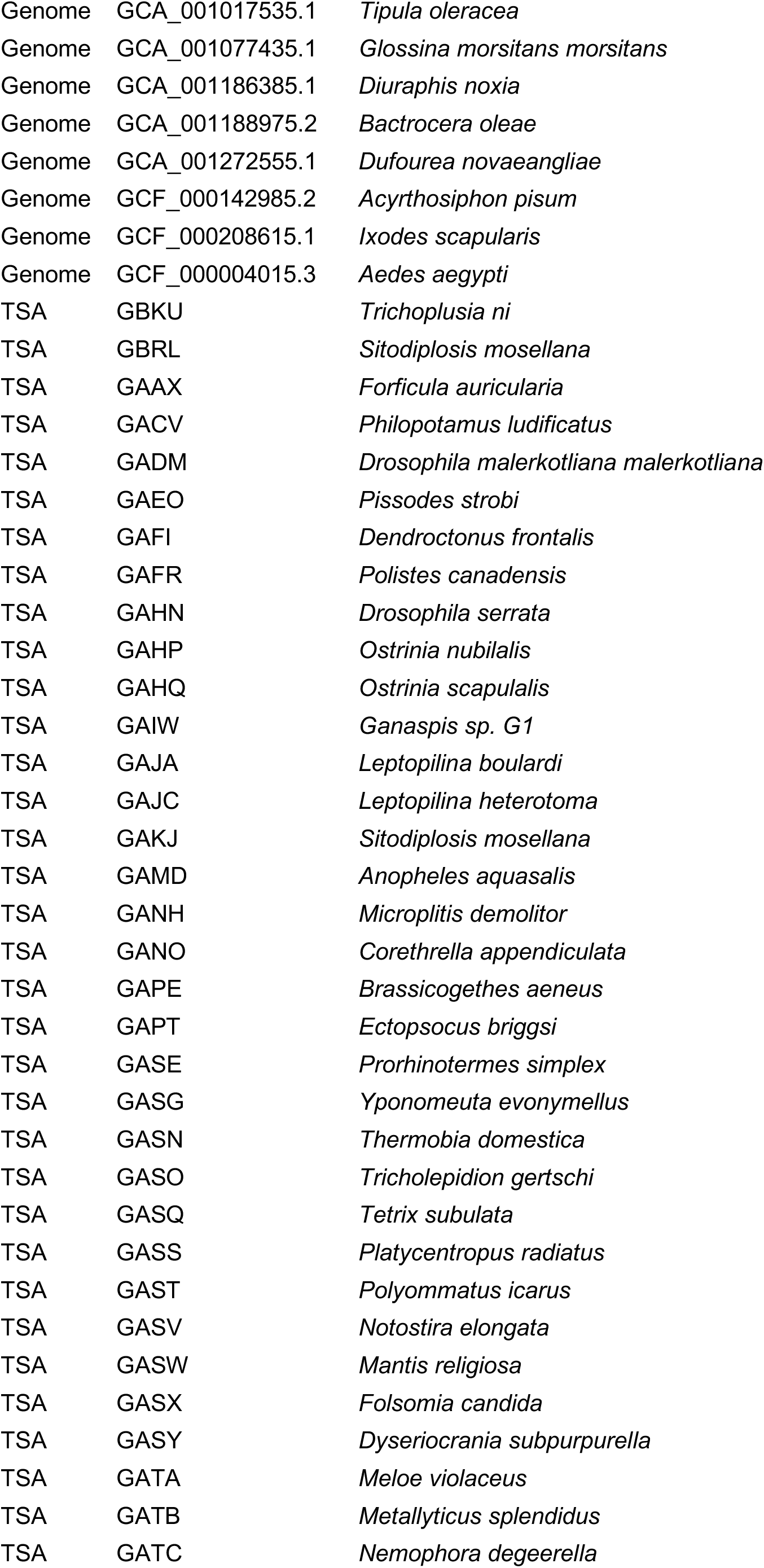
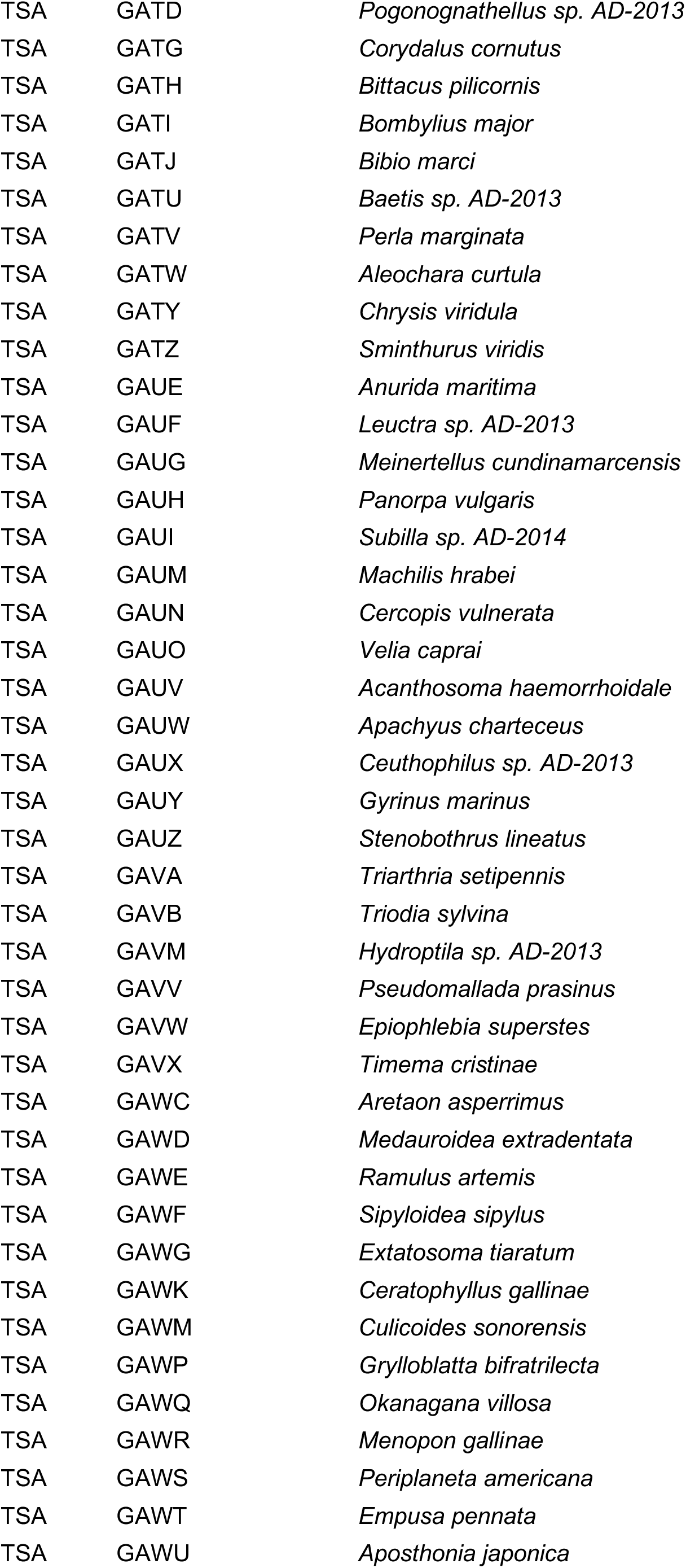
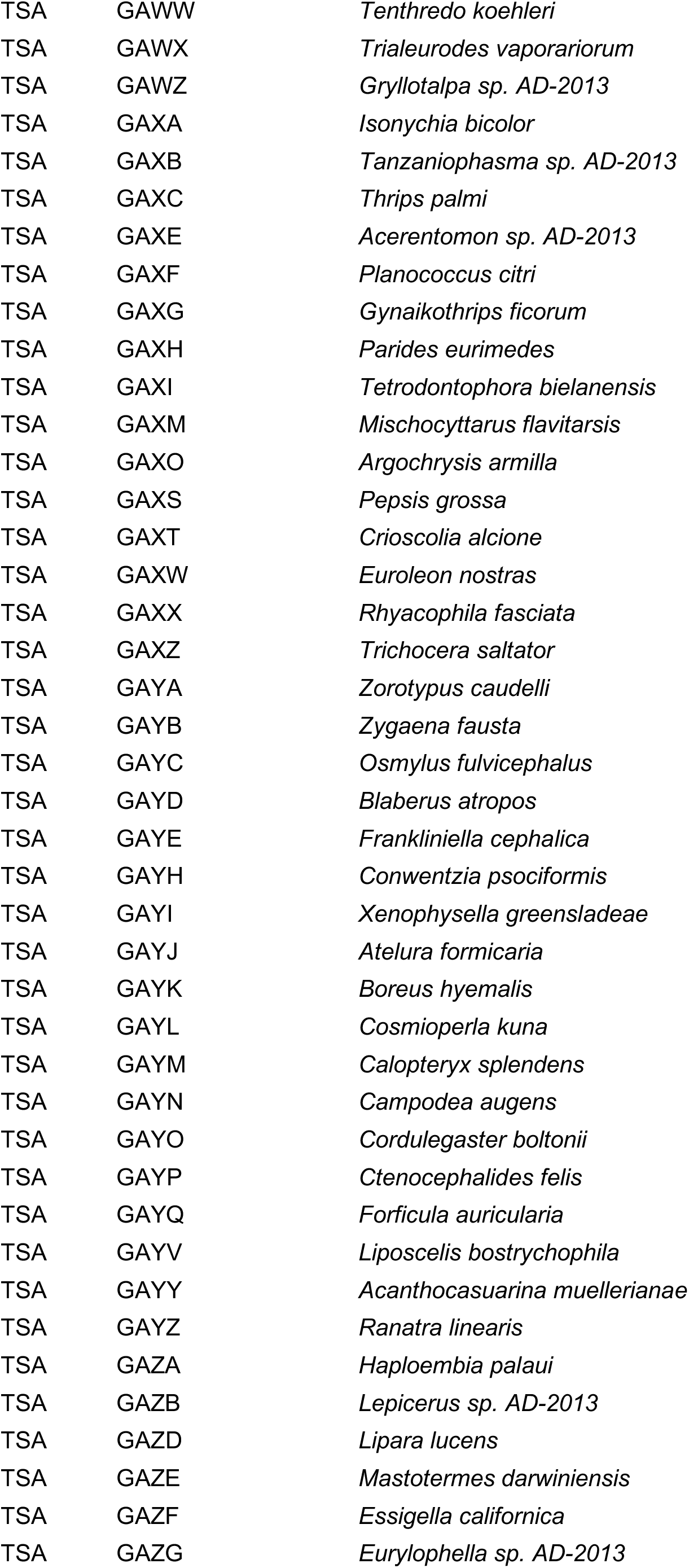
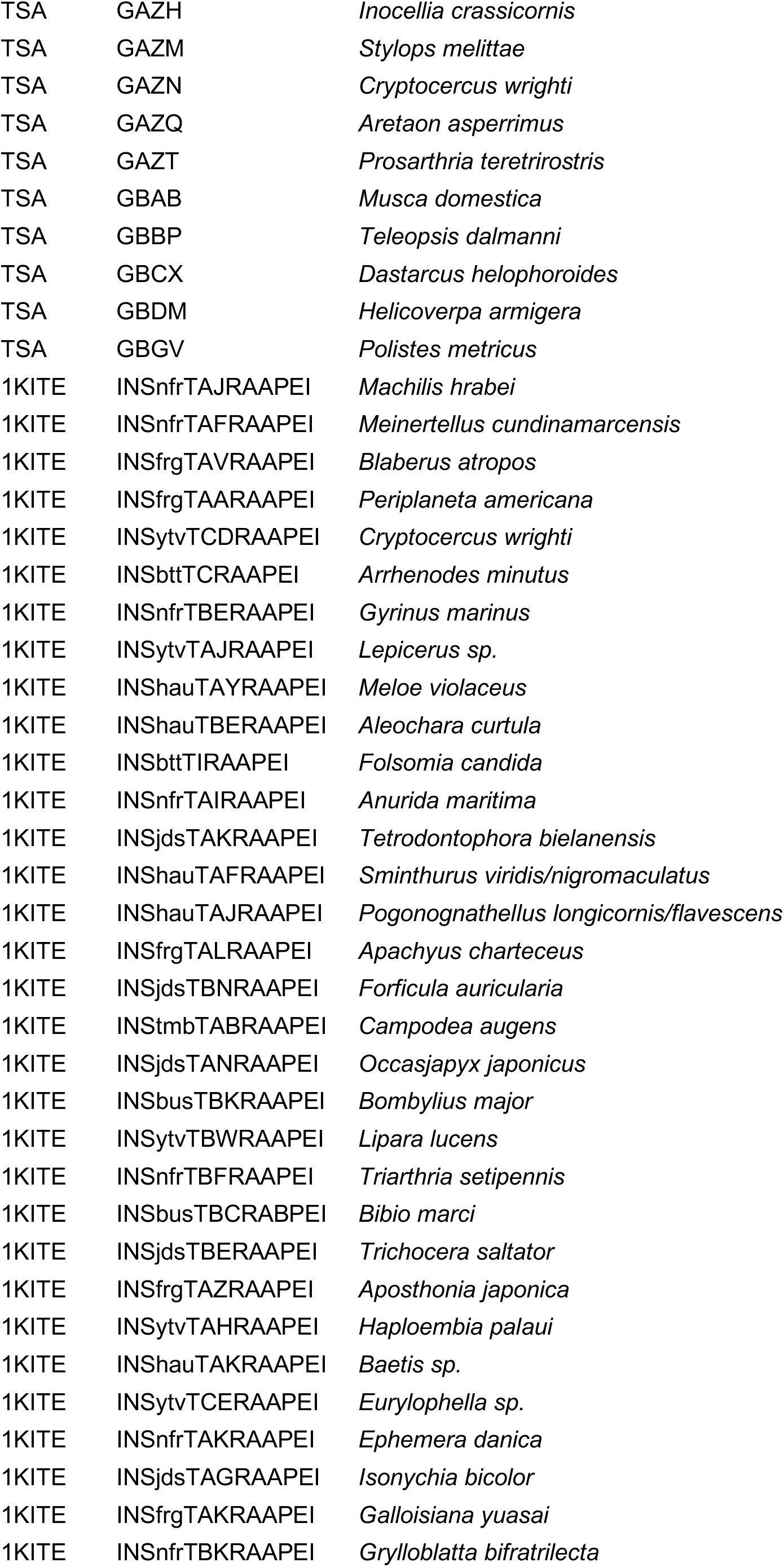
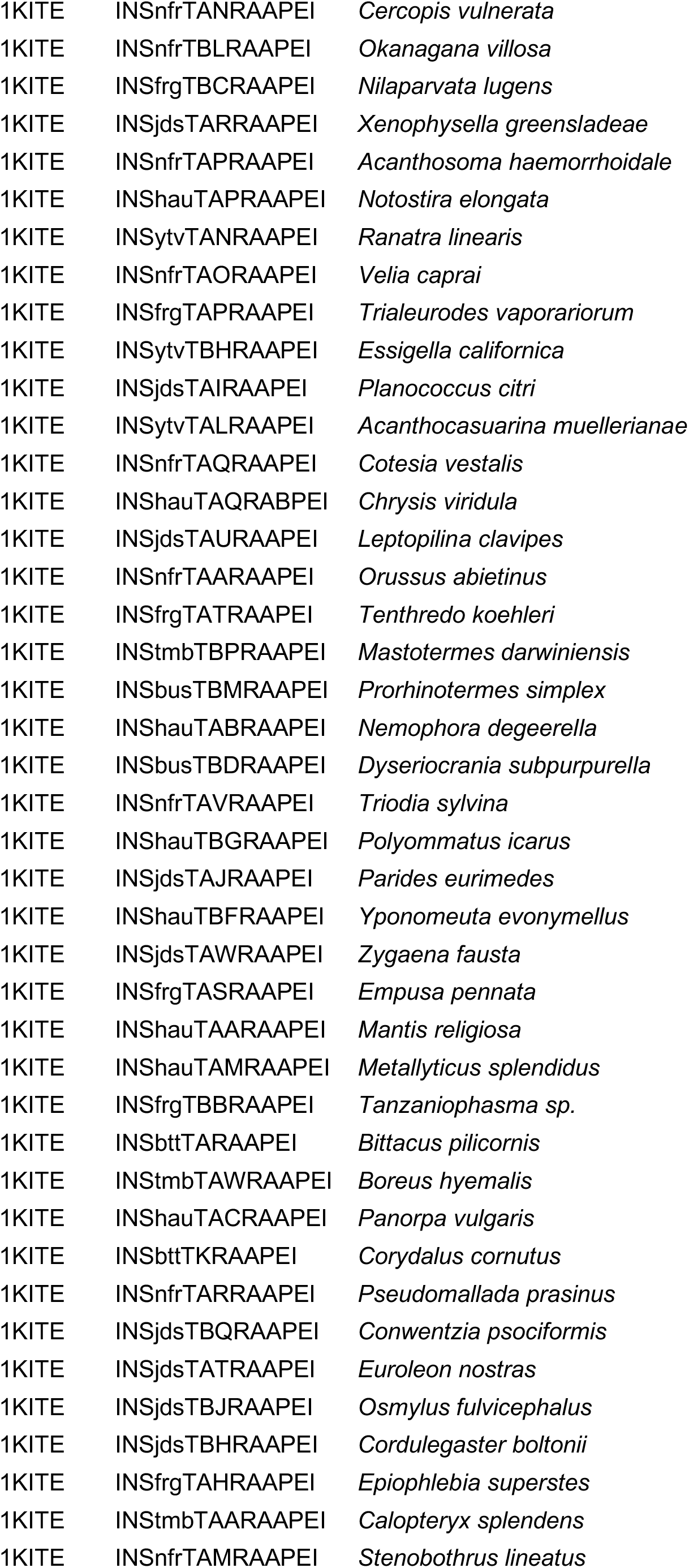
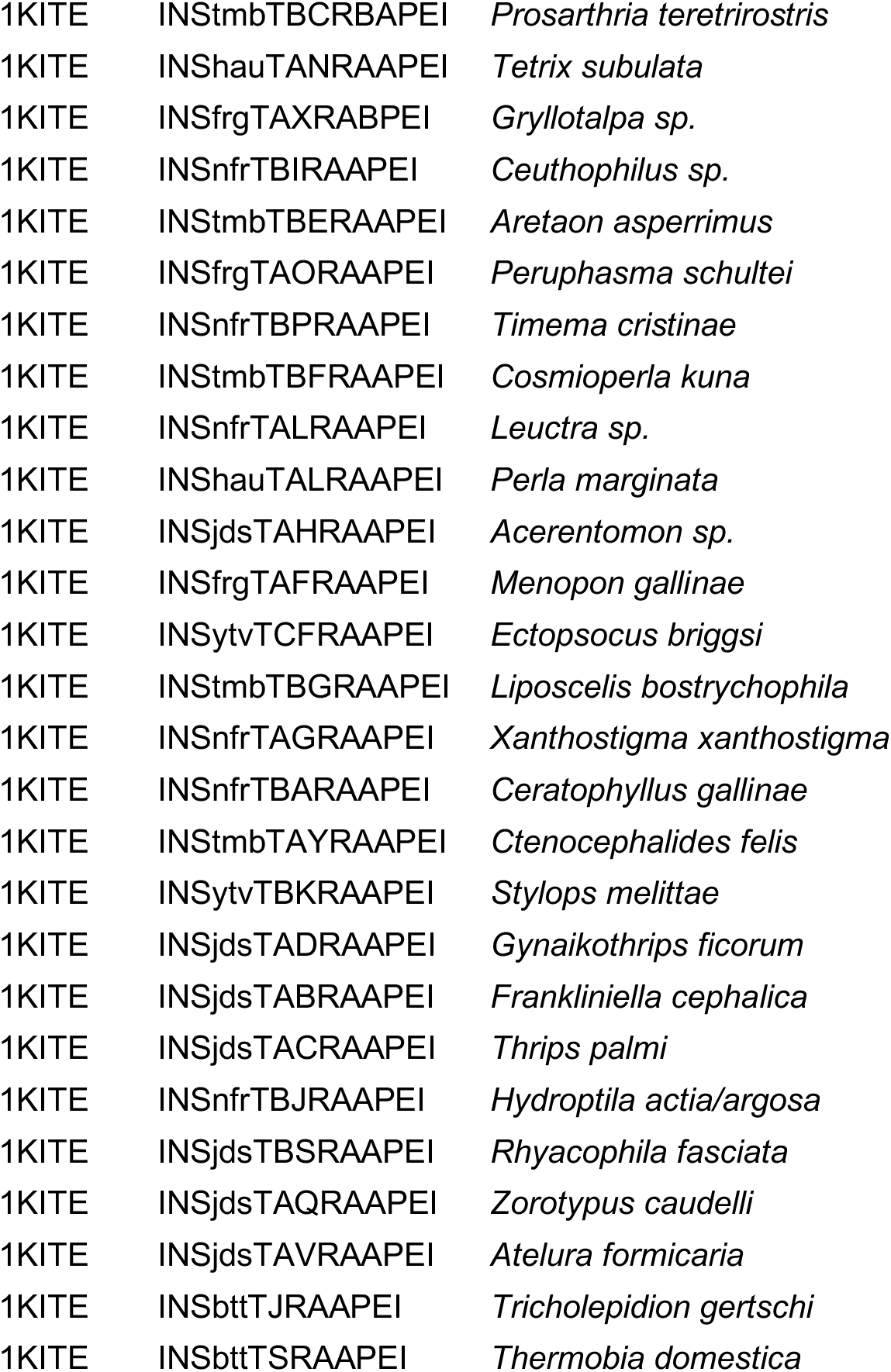
List of genomes and transcriptomes used for automated *oskar* search. List of genomes and transcriptomes that were downloaded, annotated, and searched for *oskar* sequences (*see “Hidden Markov Model (HMM) generation and alignments of the OSK and LOTUS domains*” in Methods). The table reports the database provenance (NCBI genome or TSA, or 1KITE database) and the accession number. The TSA accession ID can be searched using the NCBI TSA browser here: https://www.ncbi.nlm.nih.gov/Traces/wgs/?view=TSA.

**Supplementary Table S2:**
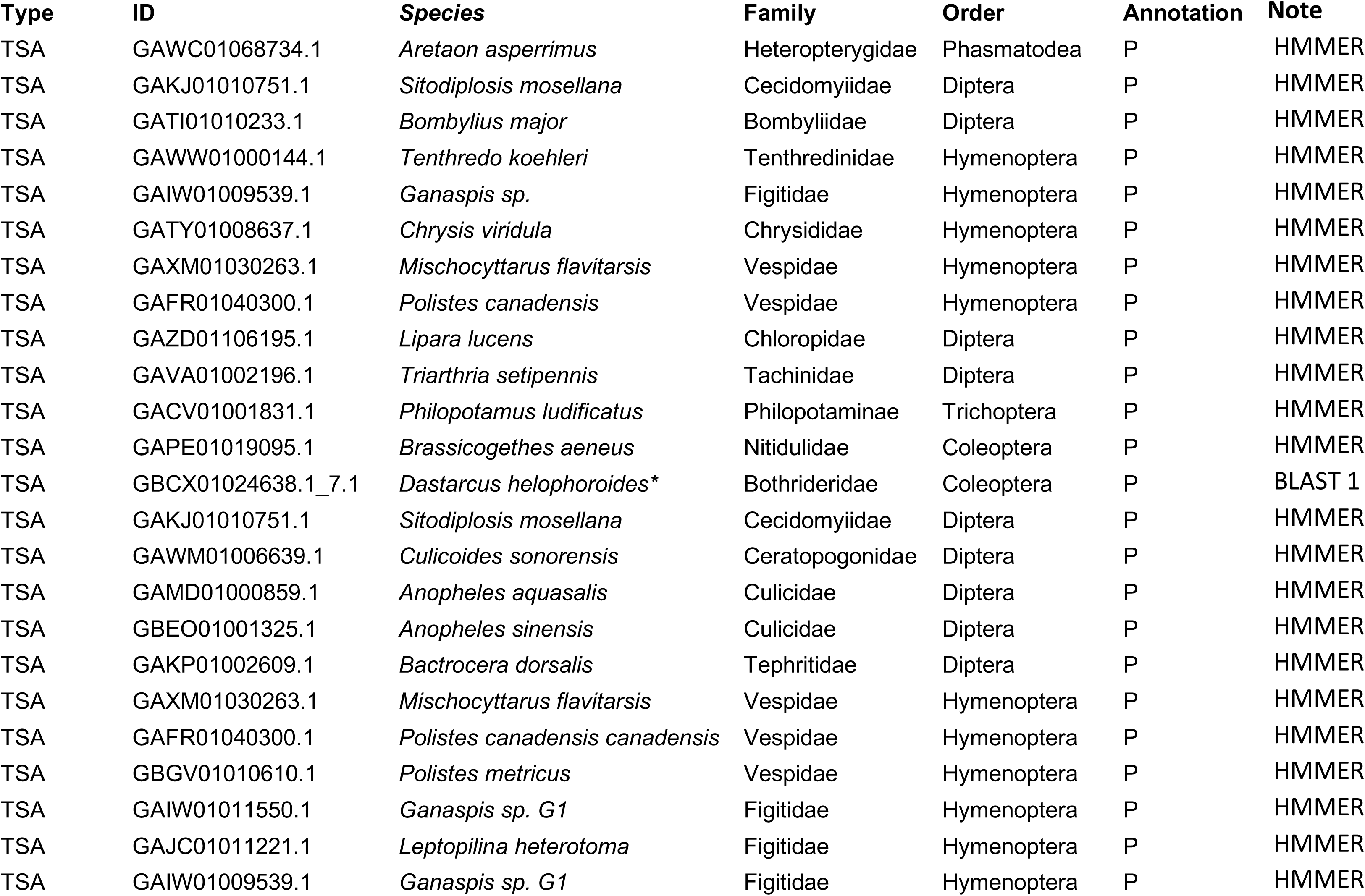
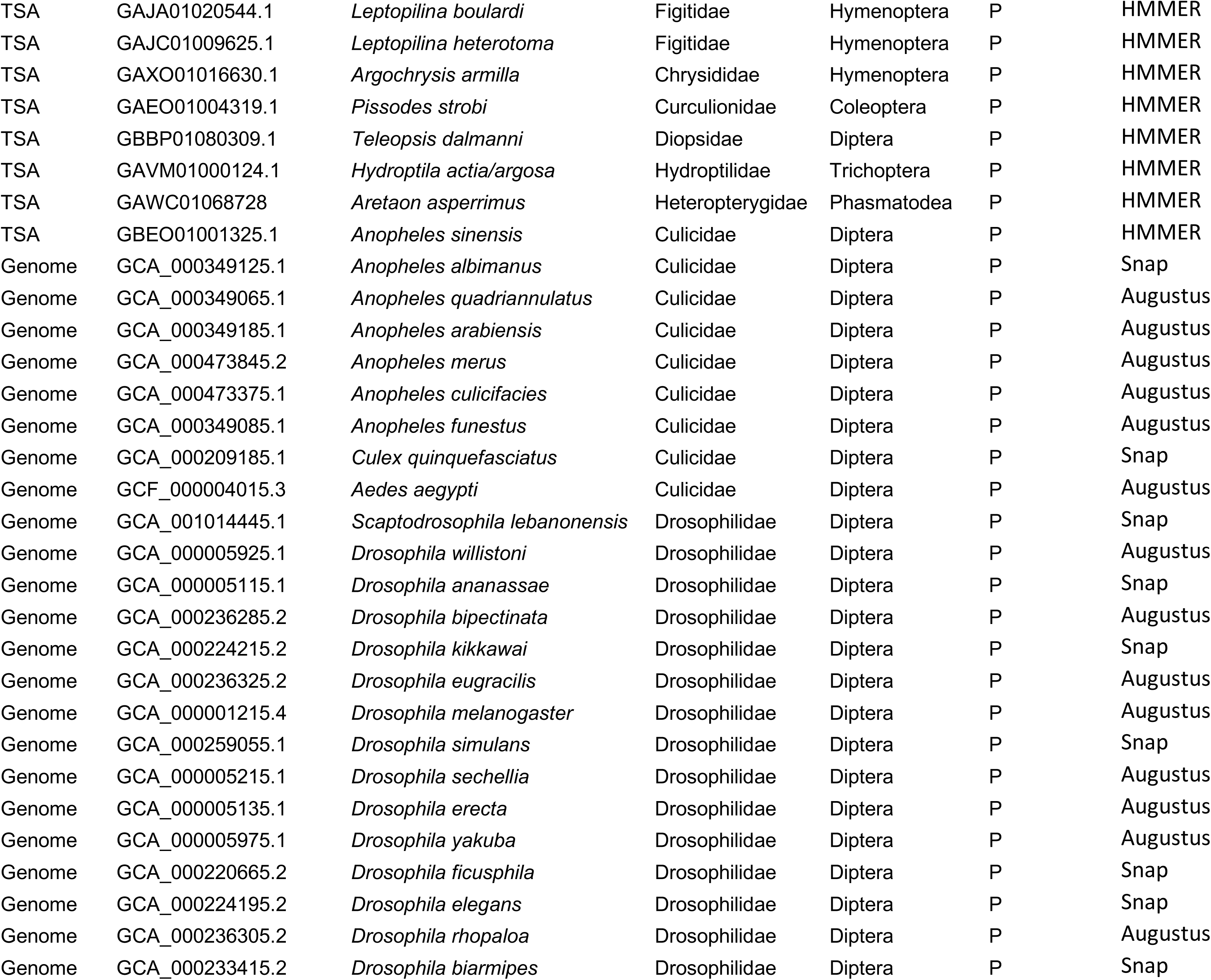
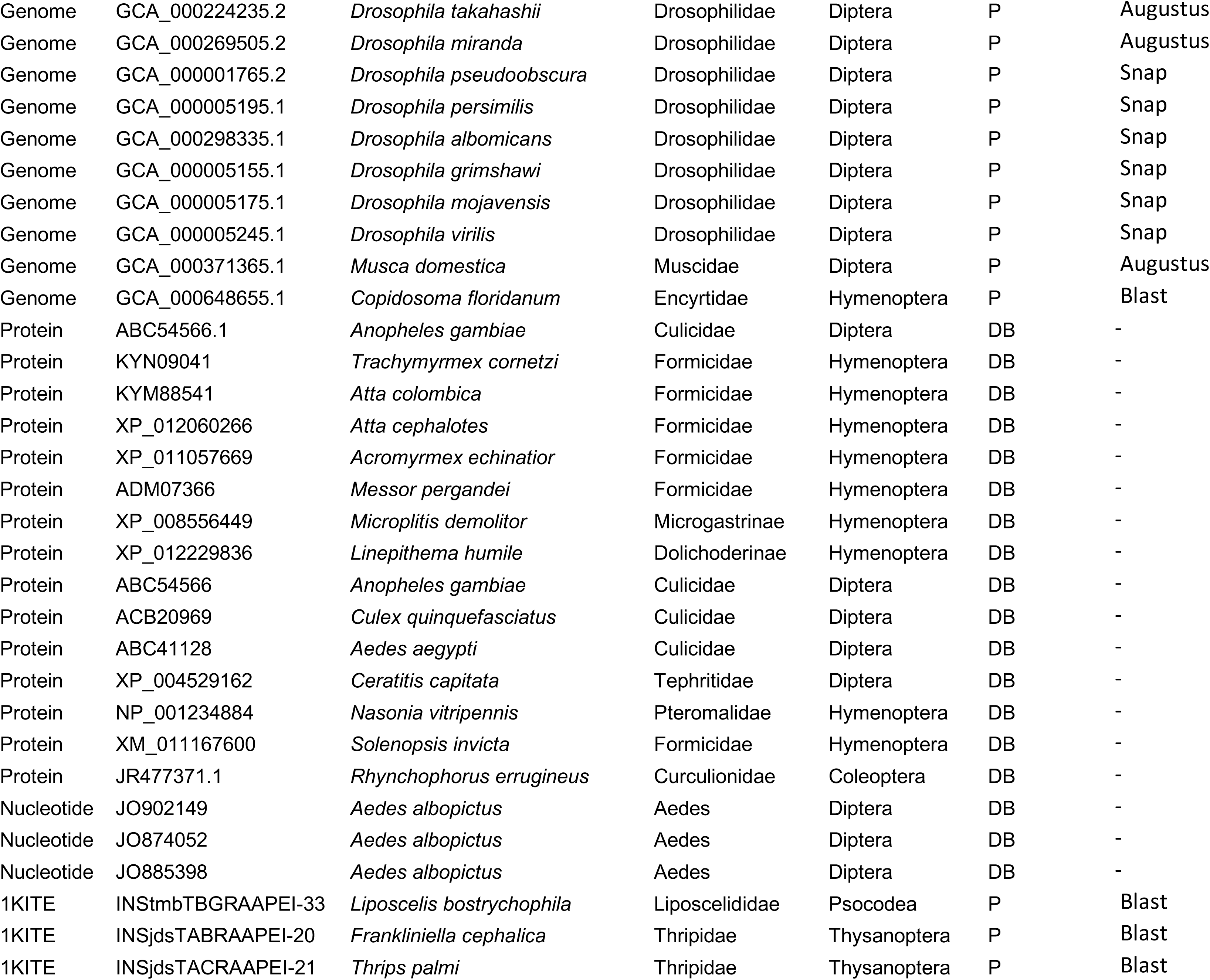
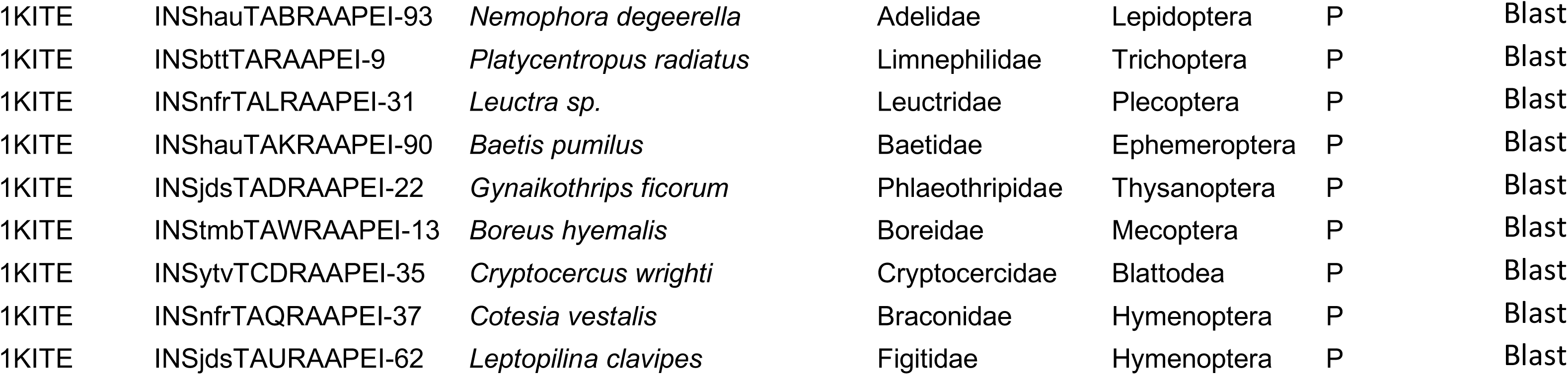
List of *oskar* sequences used in the final alignment. List of accession numbers and database provenance of the sequences used in the final alignments of Oskar analysed herein. The table contains the database provenance (*Type*), the database accession number (*ID*), the species, family and order, and extraction notes. In the “Annotation” Column, P = homolog identified by pipeline; DB = homolog identified by database annotation. *Sequence recomposed from two transcripts: GBCX01024638.1 and GBCX01024637.

**Supplementary Table S3:**
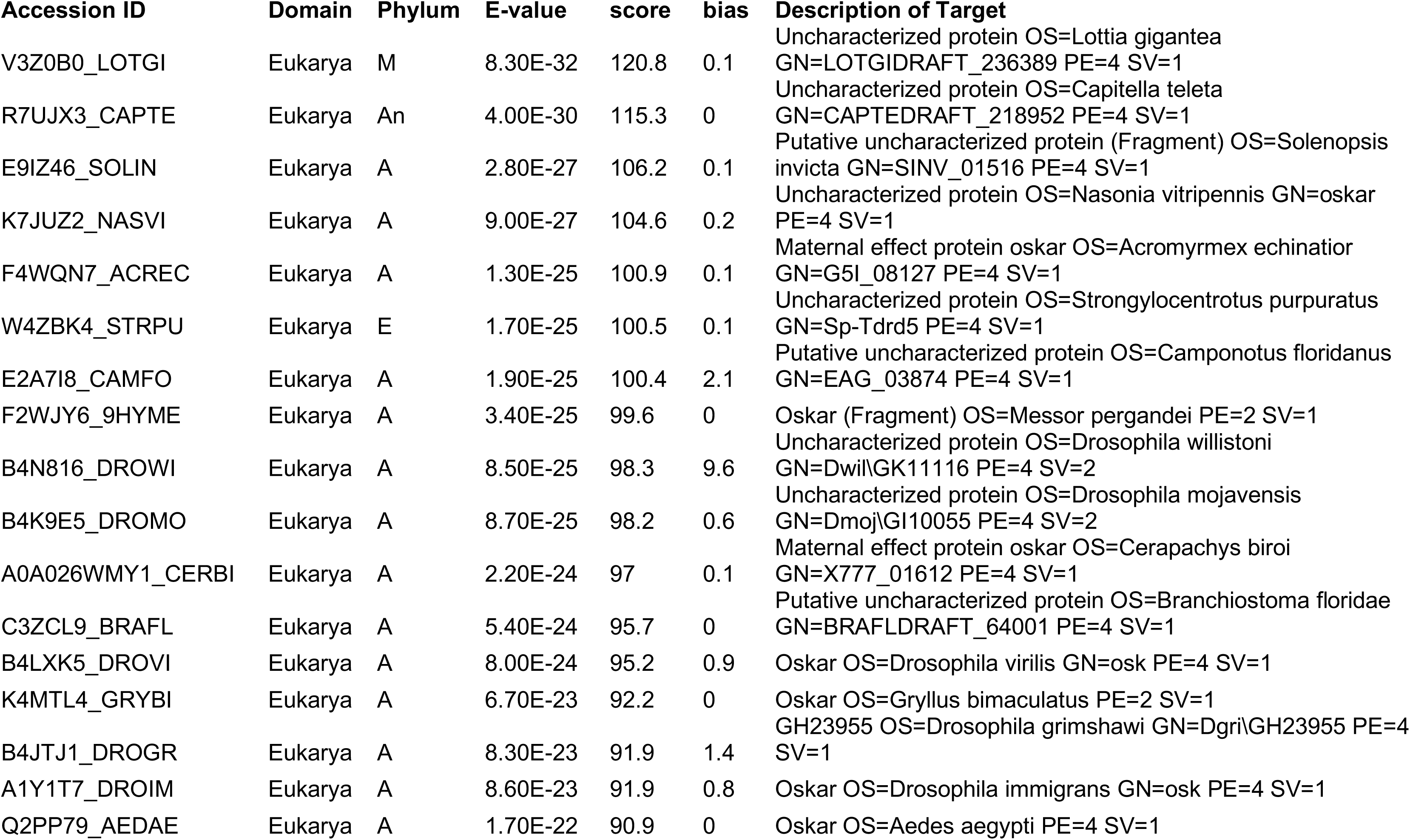
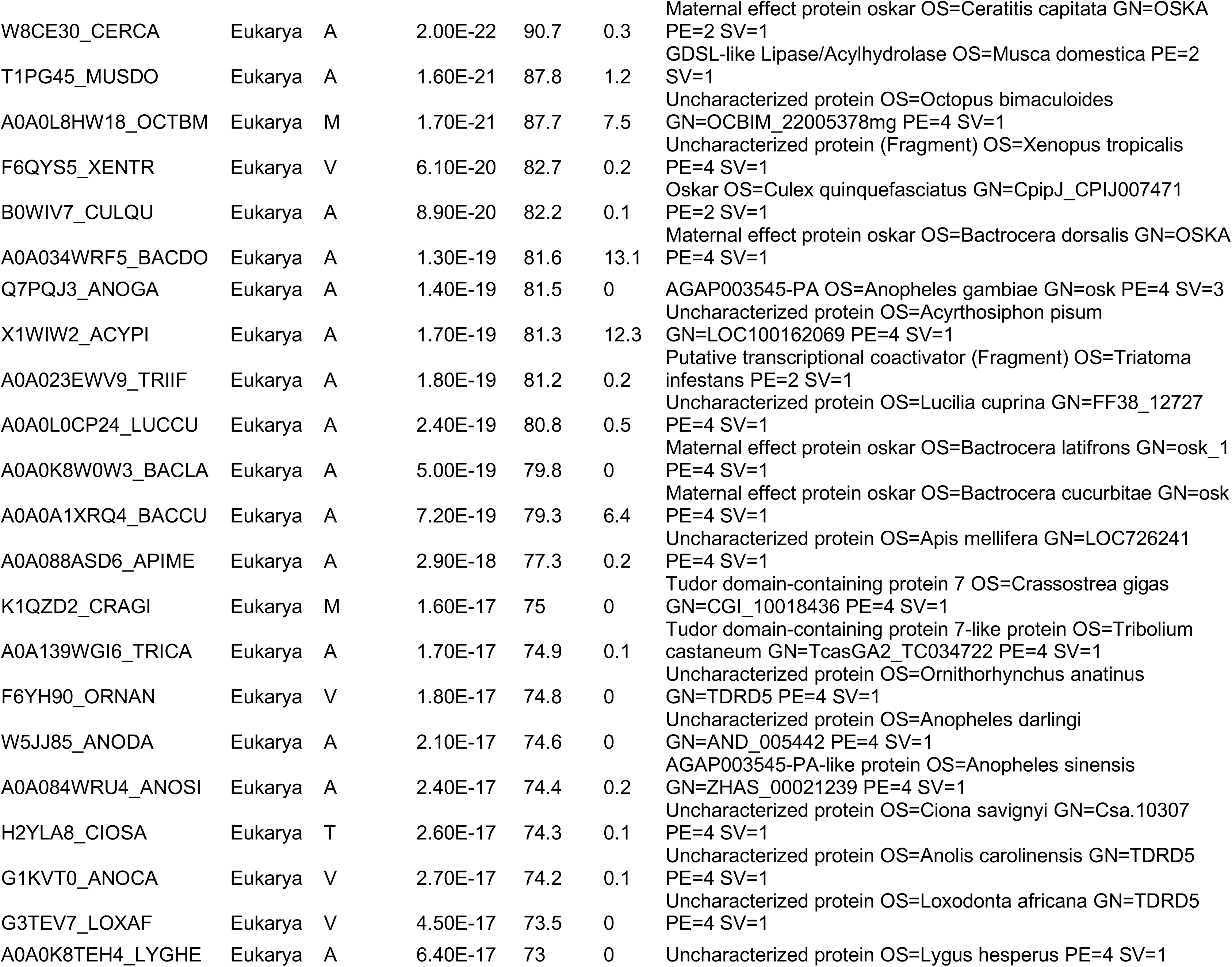
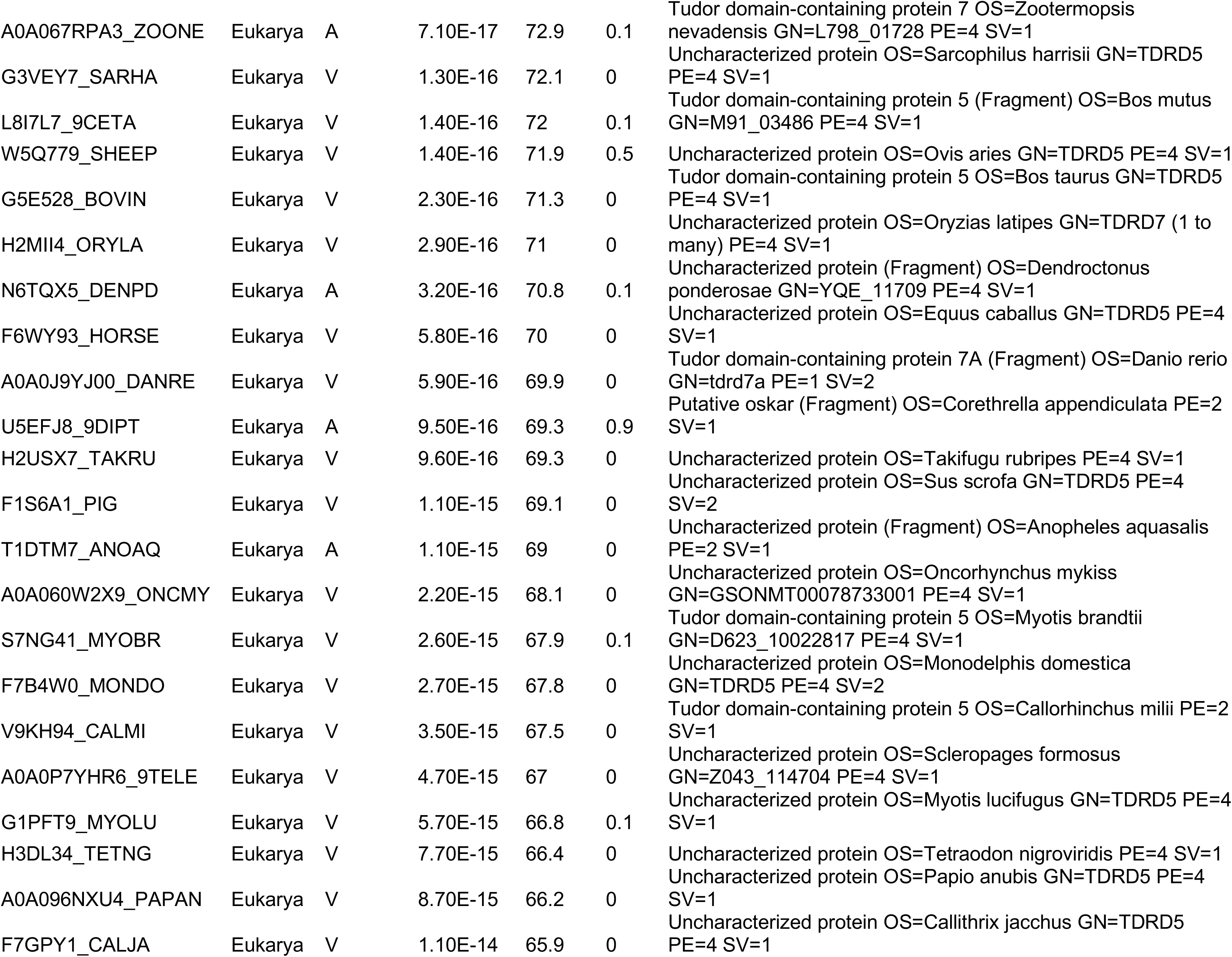
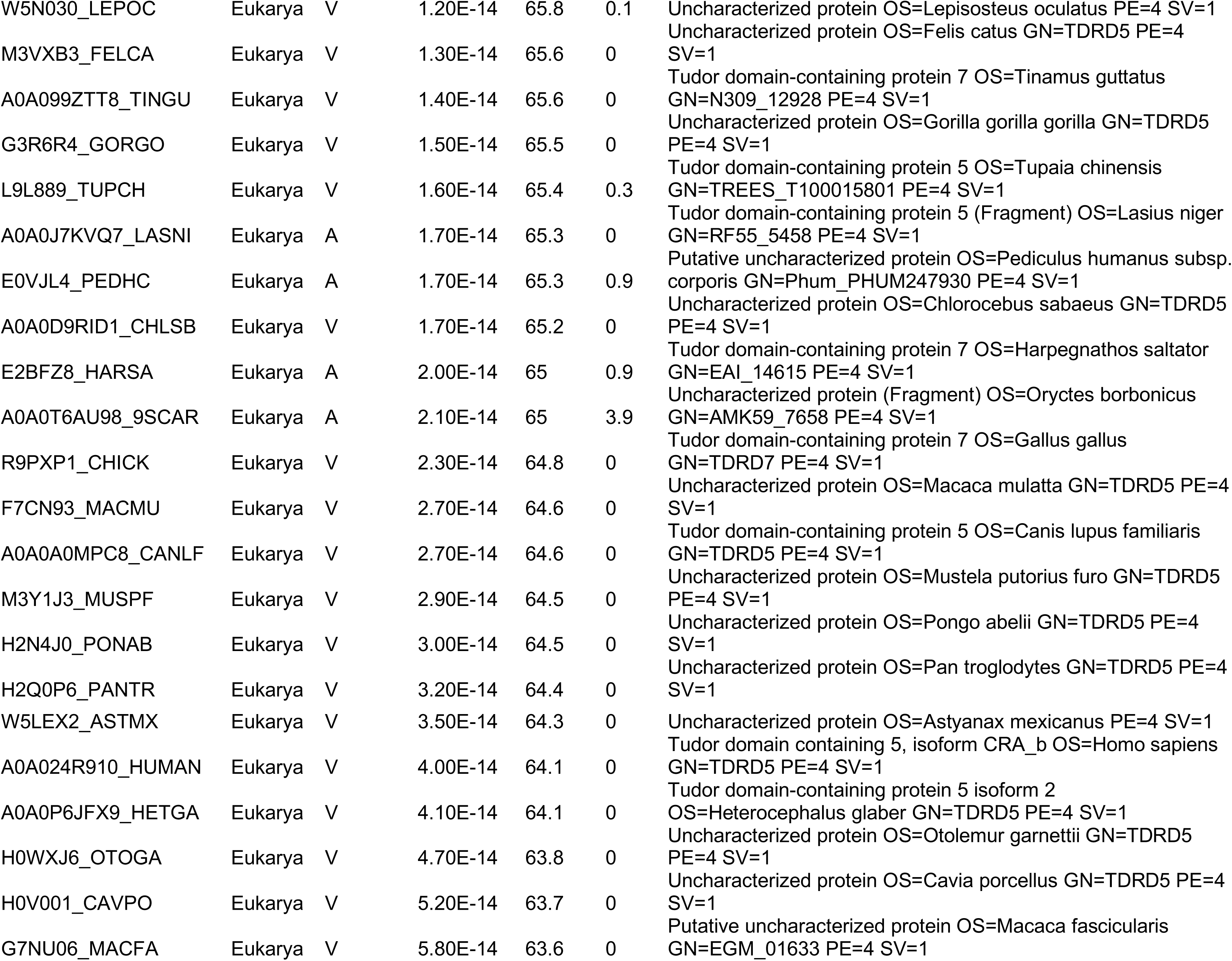
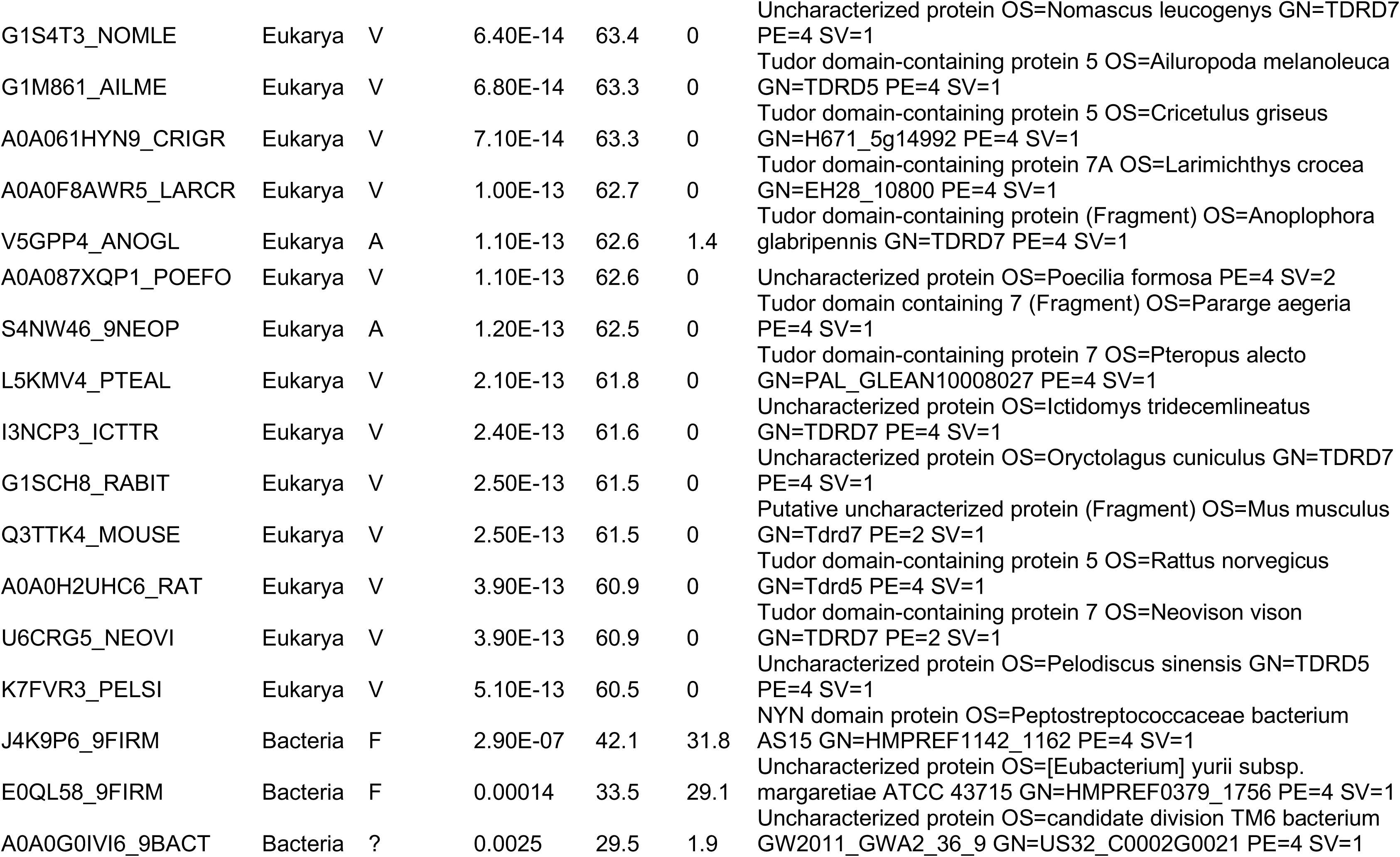
List of sequences and their BLAST results used for phylogenetic analysis of the LOTUS domain. The sequences were obtained by searching the TrEMBL database using hmmsearch and the final HMM generated for LOTUS (Supplementary files: HMM>LOTUS.hmm). Reported are the UniProtID (*Accession Number*), the Domain and Phylum origin of the sequence, the E-value, score and bias given by hmmsearch, and the description of the target from UniProt. To obtain sequences for each entry, either search UniProt directly (https://www.uniprot.org/) or consult the final alignment in Supplementary Files: Alignments>LOTUS_TREE.fasta. Phylum abbreviations: A = Arthropoda; An = Annelida; E = Echinodermata; F = Firmicutes; M = Mollusca; T = Tunicata; V = Vertebrata;= unclassified

**Supplementary Table S4:**
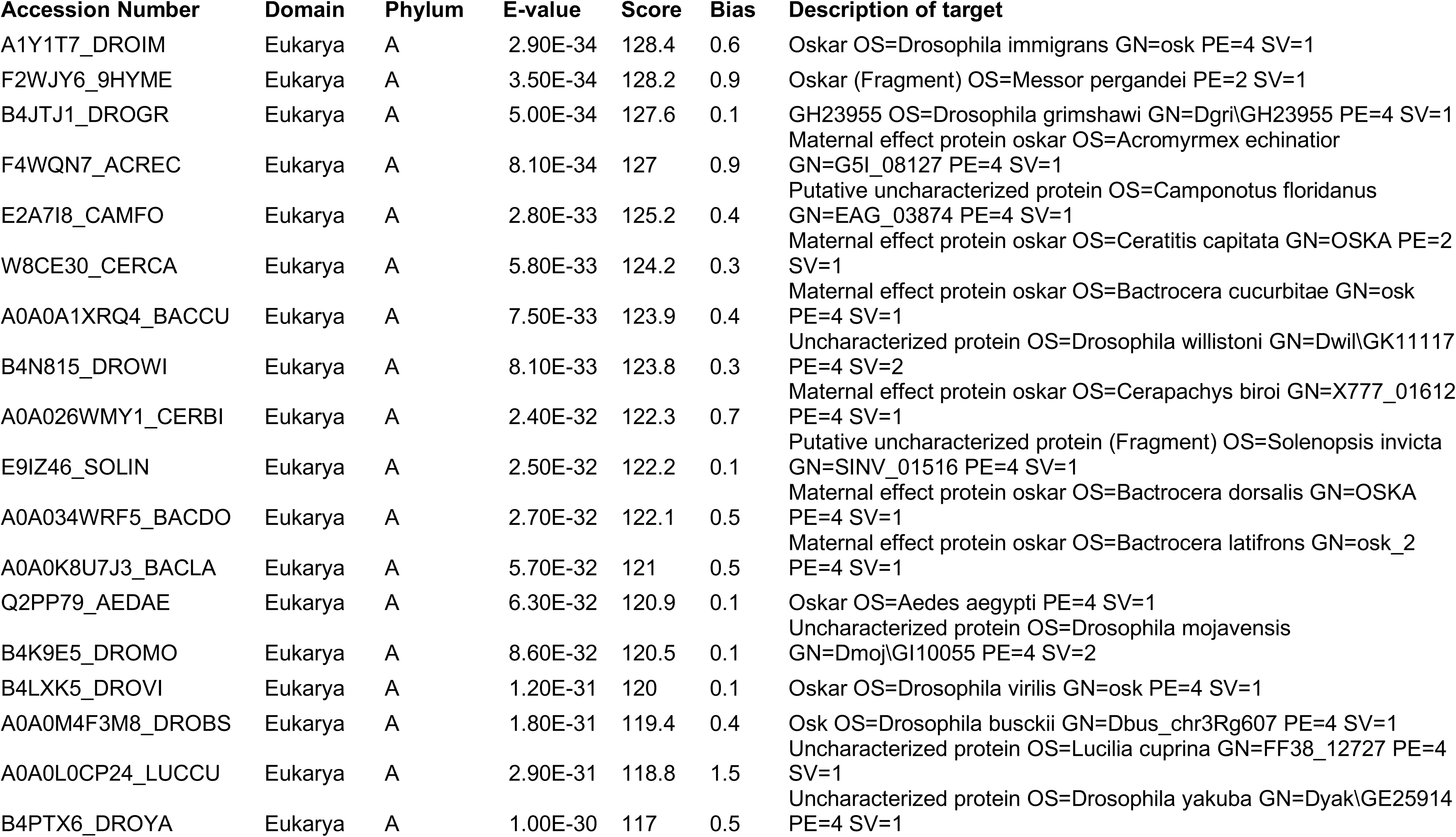
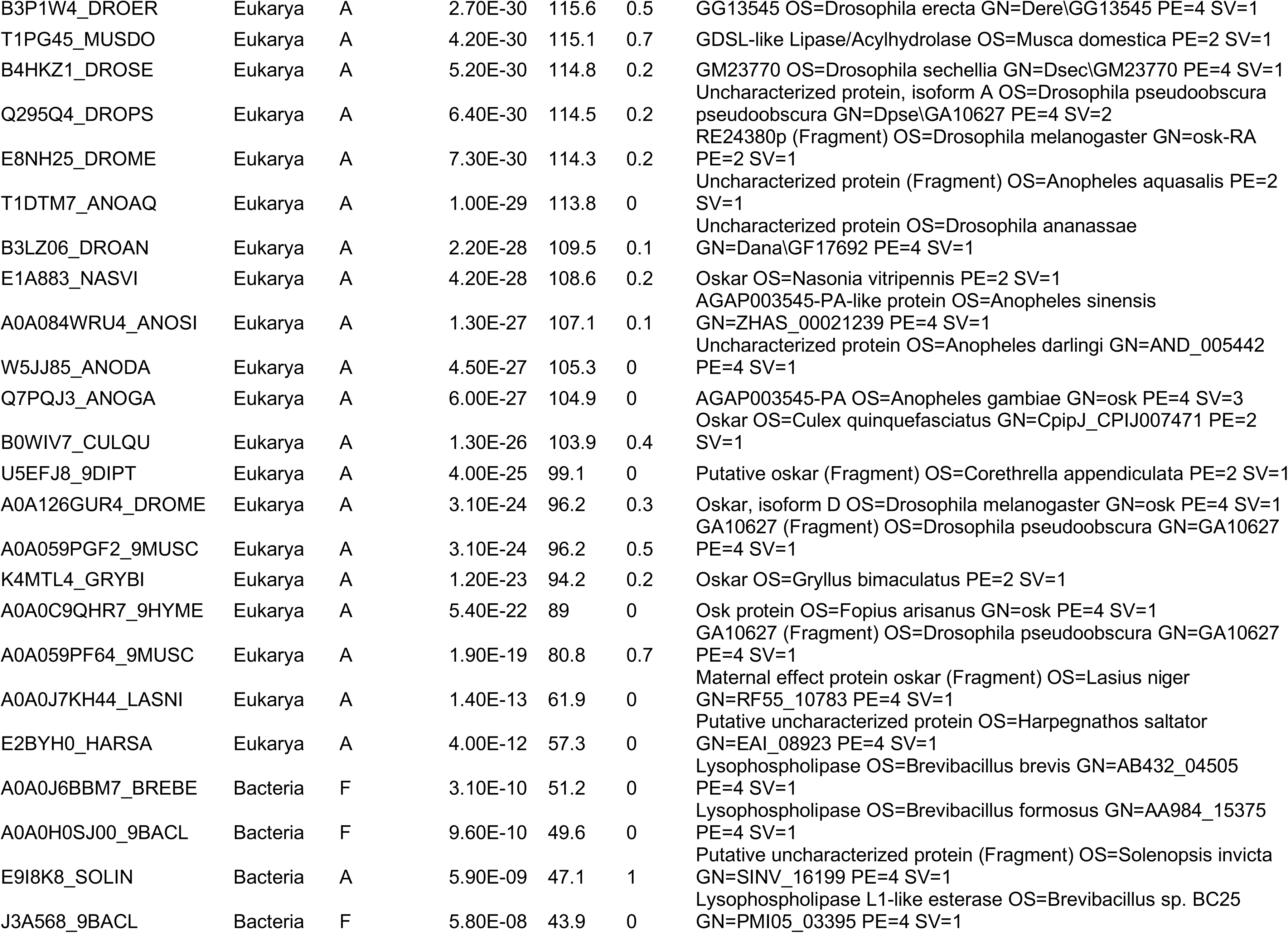
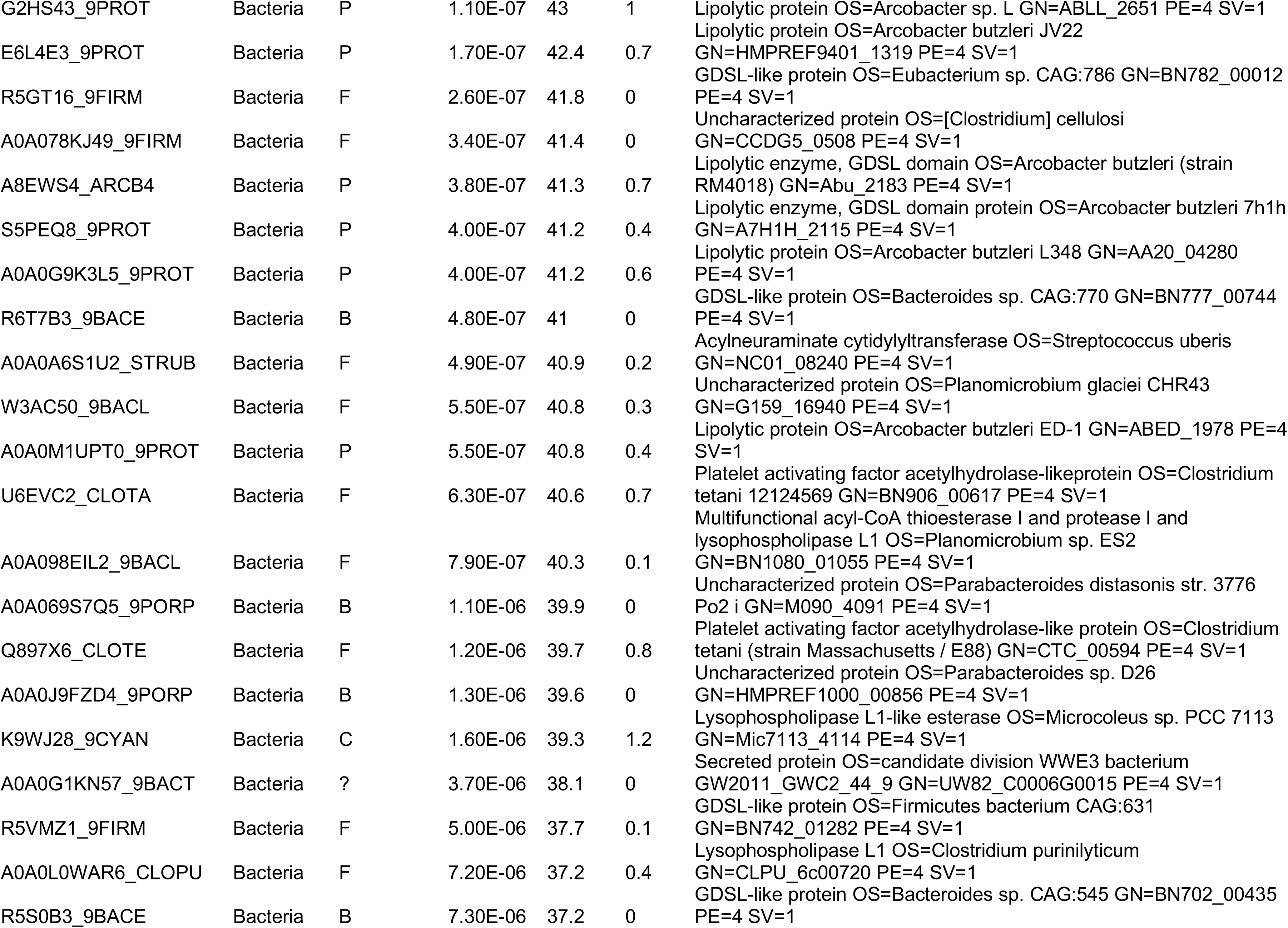
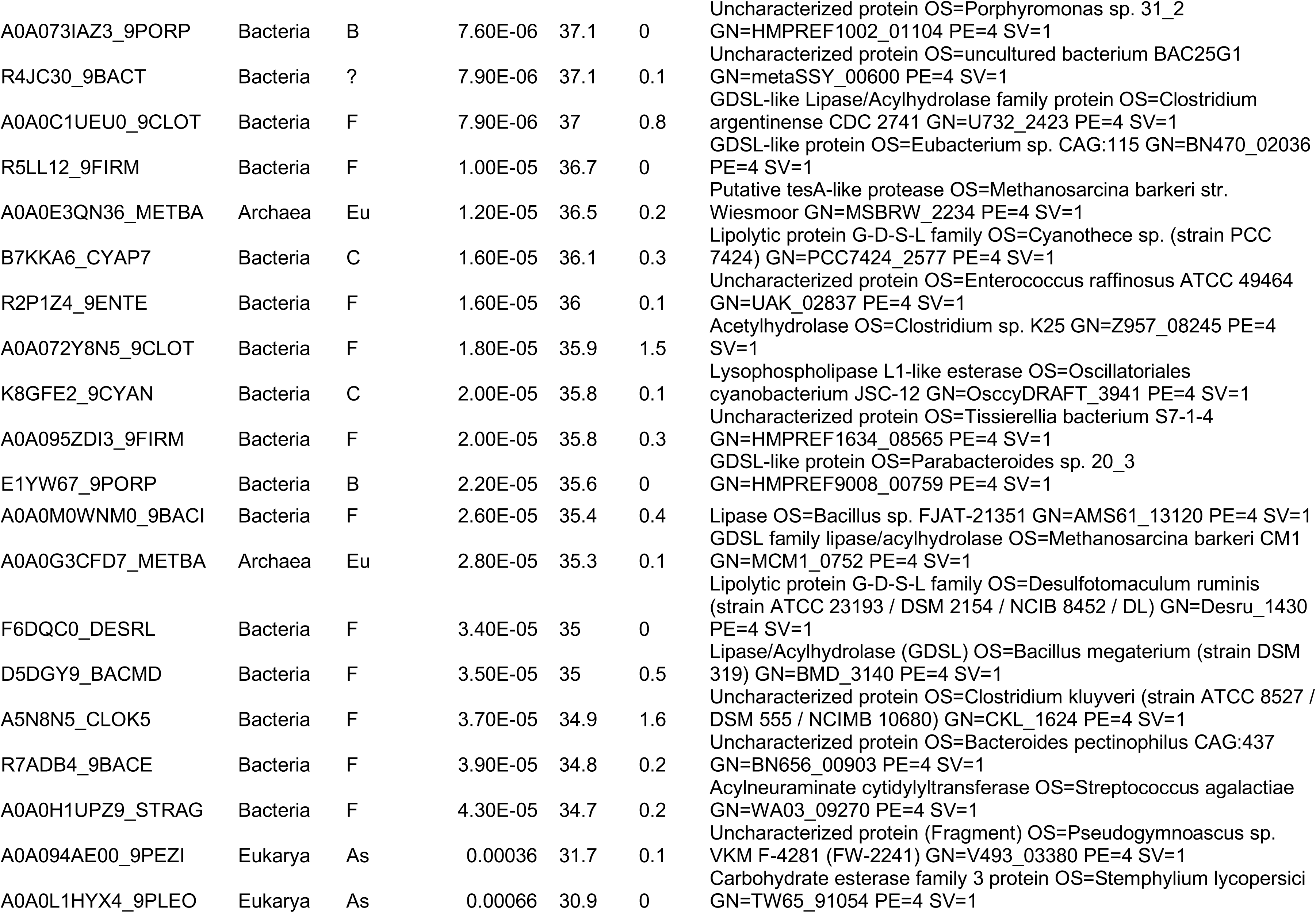
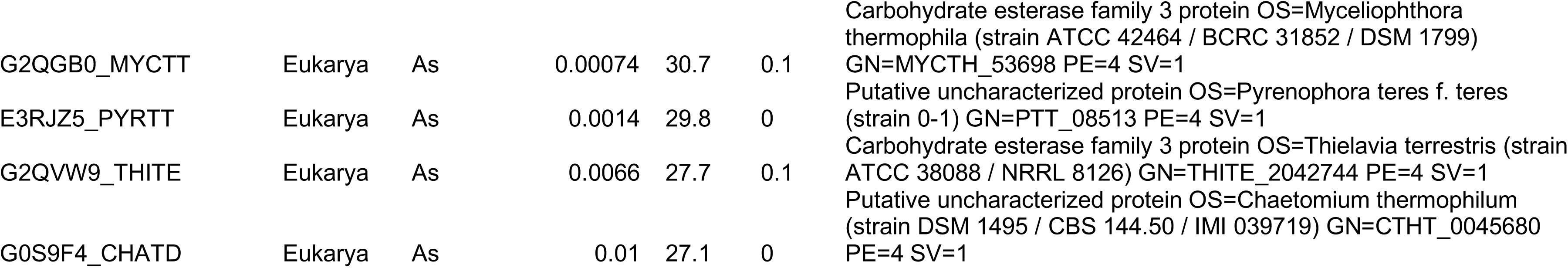
List of sequences and their BLAST results used for phylogenetic analysis of the OSK domain. The sequences were obtained by searching the TrEMBL database using hmmsearch and the final HMM generated for OSK (Supplementary files: HMM>OSK.hmm). Reported parameters are as described for Supplementary Table S3. To obtain sequences for each entry, either search UniProt directly (https://www.uniprot.org/) or consult the final alignment in Supplementary Files: Alignments>OSK_TREE.fasta. Phylum Abbreviations: A = Arthropoda; Ar = Archaea; As = Ascomycota; B = Bacteroidetes; C = Cyanobacteria; Eu = Euryarchaeota; F = Firmicutes; Fu = Fungi; P = Proteobacteria

**Supplementary Table S5:** List of genomes analyzed for codon use. This table lists the 17 genomes that were downloaded and analyzed for codon use as described in “*Selection of sequences for codon use analysis*” in Methods. All genomes can be downloaded from https://www.ncbi.nlm.nih.gov/genome/browse#!/overview/. The table lists the species name (*Species*), family (*Family*) and Order (*Order*), NCBI genome accession number (*Genome ID*), and the *oskar* NCBI Nucleotide accession number (*oskar Nucleotide ID*).

| Species | Family | Order | Genome ID | Oskar Nucleotide ID |
| --- | --- | --- | --- | --- |
| *Drosophila melanogaster* | Drosophilidae | Diptera | GCA_001014345.1 | NM_169248.4 |
| *Nasonia vitripennis* | Pteromalidae | Hymenoptera | GCA_000002325.2 | HM535628.1 |
| *Culex quinquefasciatus* | Culicidae | Diptera | GCA_000209185.1 | EU517695.1 |
| *Drosophila virilis* | Drosophilidae | Diptera | GCA_000005245.1 | L22556.1 |
| *Ceratitis capitata* | Tephritidae | Diptera | GCA_000347755.4 | LOC101450245 |
| *Musca domestica* | Muscidae | Diptera | GCA_000371365.1 | LOC101890691 |
| *Acromyrmex echinator* | Formicidae | Hymenoptera | GCA_000204515.1 | LOC105147973 |
| *Harpegnathos saltator* | Formicidae | Hymenoptera | GCA_000147195.1 | LOC105187957 |
| *Bactrocera dorsalis* | Tephritidae | Diptera | GCA_000789215.2 | LOC105232054 |
| *Fopius arisanus* | Braconidae | Hymenoptera | GCA_000806365.1 | LOC105267990 |
| *Athalia rosae* | Tenthredinidae | Hymenoptera | GCA_000344095.2 | LOC105692731 |
| *Orussus abietinus* | Orussidae | Hymenoptera | GCA_000612105.2 | LOC105696794 |
| *Stomoxys calcitrans* | Muscidae | Diptera | GCA_001015335.1 | LOC106086381 |
| *Bactrocera oleae* | Tephritidae | Diptera | GCA_001188975.2 | LOC106622417 |
| *Copidosoma floridanum* | Encyrtidae | Hymenoptera | GCA_000648655.2 | LOC106642594 |
| *Polistes canadensis* | Vespidae | Hymenoptera | GCA_001313835.1 | LOC106790143 |
| *Neodiprion lecontei* | Diprionidae | Hymenoptera | GCF_001263575.1 | LOC107223453 |

## References

1. J. S. Taylor, J. Raes, Duplication and divergence: the evolution of new genes and old ideas. Annual review of genetics 38, 615–643 (2004).

2. D. Tautz, T. Domazet-Loso, The evolutionary origin of orphan genes. Nat. Rev. Genet. 12, 692–702 (2011).

3. P. J. Wittkopp, G. Kalay, Cis-regulatory elements: molecular mechanisms and evolutionary processes underlying divergence. Nat. Rev. Genet. 13, 59–69 (2011).

4. H. E. Hoekstra, J. A. Coyne, The locus of evolution: evo devo and the genetics of adaptation. Evolution 61, 995–1016 (2007).

5. S. Chen, B. H. Krinsky, M. Long, New genes as drivers of phenotypic evolution. Nat. Rev. Genet. 14, 645–660 (2013).

6. W. Zhang, P. Landback, A. R. Gschwend, B. Shen, M. Long, New genes drive the evolution of gene interaction networks in the human and mouse genomes. Genome Biol. 16, 202 (2015).

7. Y. E. Zhang, P. Landback, M. Vibranovski, M. Long, New genes expressed in human brains: implications for annotating evolving genomes. BioEssays 34, 982–991 (2012).

8. S. Chen et al., Frequent recent origination of brain genes shaped the evolution of foraging behavior in Drosophila. Cell Reports 1, 118–132 (2012).

9. J. C. Dunning Hotopp et al., Widespread lateral gene transfer from intracellular bacteria to multicellular eukaryotes. Science (New York, NY) 317, 1753–1756 (2007).

10. R. Acuna et al., Adaptive horizontal transfer of a bacterial gene to an invasive insect pest of coffee. Proc Natl Acad Sci U S A 109, 4197–4202 (2012).

11. D. B. Sloan et al., Parallel histories of horizontal gene transfer facilitated extreme reduction of endosymbiont genomes in sap-feeding insects. Mol. Biol. Evol. 31, 857–871 (2014).

12. F. Husnik et al., Horizontal gene transfer from diverse bacteria to an insect genome enables a tripartite nested mealybug symbiosis. Cell 153, 1567–1578 (2013).

13. R. Lehmann, Germ Plasm Biogenesis--An Oskar-Centric Perspective. Curr. Top. Dev. Biol. 116, 679–707 (2016).

14. R. Lehmann, C. Nüsslein-Volhard, Abdominal Segmentation, Pole Cell Formation, and Embryonic Polarity Require the Localized Activity of *oskar*, a Maternal Gene in *Drosophila*. Cell 47, 144–152 (1986).

15. B. Ewen-Campen, J. R. Srouji, E. E. Schwager, C. G. Extavour, *oskar* Predates the Evolution of Germ Plasm in Insects. Curr. Biol. 22, 2278–2283 (2012).

16. C. G. Extavour, M. E. Akam, Mechanisms of germ cell specification across the metazoans: epigenesis and preformation. Development 130, 5869–5884 (2003).

17. J. A. Lynch et al., The Phylogenetic Origin of *oskar* Coincided with the Origin of Maternally Provisioned Germ Plasm and Pole Cells at the Base of the Holometabola. PLoS Genetics 7, e1002029 (2011).

18. B. Ewen-Campen, S. Donoughe, D. N. Clarke, C. G. Extavour, Germ cell specification requires zygotic mechanisms rather than germ plasm in a basally branching insect. Curr. Biol. 23, 835–842 (2013).

19. E. Abouheif, Evolution: oskar Reveals Missing Link in Co-optive Evolution. Curr. Biol. 23, R24–R25 (2012).

20. M. Jeske et al., The Crystal Structure of the *Drosophila* Germline Inducer Oskar Identifies Two Domains with Distinct Vasa Helicase- and RNA-Binding Activities. Cell Reports 12, 587–598 (2015).

21. M. Jeske, C. W. Muller, A. Ephrussi, The LOTUS domain is a conserved DEAD-box RNA helicase regulator essential for the recruitment of Vasa to the germ plasm and nuage. Genes Dev. 31, 939–952 (2017).

22. A. Ephrussi, R. Lehmann, Induction of germ cell formation by *oskar*. Nature 358, 387–392 (1992).

23. J. Kim-Ha, J. L. Smith, P. M. Macdonald, *oskar* mRNA is localized to the posterior pole of the *Drosophila* oocyte. Cell 66, 23–35 (1991).

24. N. Yang et al., Structure of *Drosophila* Oskar reveals a novel RNA binding protein. Proc. Natl. Acad. Sci. USA 112, 11541–11546 (2015).

25. A. H. Kirk-Spriggs, B. J. Sinclair, Eds., Manual of Afrotropical Diptera, (South African National Biodiversity Institute, Pretoria, South Africa, 2017), vol. 1.

26. R. S. Peters et al., Evolutionary History of the Hymenoptera. Curr. Biol. 27, 1013–1018 (2017).

27. N. F. Vanzo, A. Ephrussi, Oskar anchoring restricts pole plasm formation to the posterior of the *Drosophila* oocyte. Development 129, 3705–3714 (2002).

28. T. R. Hurd et al., Long Oskar Controls Mitochondrial Inheritance in *Drosophila melanogaster*. Dev. Cell 39, 560–571 (2016).

29. B. Misof et al., Phylogenomics resolves the timing and pattern of insect evolution. Science 346, 763–767 (2014).

30. D. L. Swofford, G. J. Olsen, P. J. Waddell, D. M. Hillis, in Molecular Systematics 2nd Edition, D. M. Hillis, C. Moritz, B. K. Mable, Eds. (Sinauer Associates, Inc., Sinauer, MA, 1996), chap. 11, pp. 407–453.

31. S. H. Church, J. F. Ryan, C. W. Dunn, Automation and Evaluation of the SOWH Test with SOWHAT. Syst. Biol. 64, 1048–1058 (2015).

32. D. Wheeler, A. J. Redding, J. H. Werren, Characterization of an ancient lepidopteran lateral gene transfer. PLoS ONE 8, e59262 (2013).

33. S. T. Chepkemoi et al., Identification of Spiroplasmainsolitum symbionts in Anopheles gambiae. Wellcome Open Research 2, 90 (2017).

34. E. Zchori-Fein, S. J. Perlman, S. E. Kelly, N. Katzir, M. S. Hunter, Characterization of a ‘Bacteroidetes’ symbiont in Encarsia wasps (Hymenoptera: Aphelinidae): proposal of ‘Candidatus Cardinium hertigii’. International Journal of Systematic and Evolutionary Microbiology 54, 961–968 (2004).

35. E. Zchori-Fein, S. J. Perlman, Distribution of the bacterial symbiont Cardinium in arthropods. Mol. Ecol. 13, 2009–2016 (2004).

36. T. Tuller, Codon bias, tRNA pools and horizontal gene transfer. Mobile Genetic Elements 1, 75–77 (2011).

37. P. M. Sharp, W. H. Li, The codon Adaptation Index--a measure of directional synonymous codon usage bias, and its potential applications. Nucleic Acids Res. 15, 1281–1295 (1987).

38. J. J. Li, G. L. Chew, M. D. Biggin, Quantitative principles of cis-translational control by general mRNA sequence features in eukaryotes. Genome Biol. 20, 162 (2019).

39. C. H. Yu et al., Codon Usage Influences the Local Rate of Translation Elongation to Regulate Co-translational Protein Folding. Mol Cell 59, 744–754 (2015).

40. S. Pechmann, J. Frydman, Evolutionary conservation of codon optimality reveals hidden signatures of cotranslational folding. Nat Struct Mol Biol 20, 237–243 (2013).

41. M. Zhou, T. Wang, J. Fu, G. Xiao, Y. Liu, Nonoptimal codon usage influences protein structure in intrinsically disordered regions. Mol. Microbiol. 97, 974–987 (2015).

42. P. Krishnakumar et al., Functional equivalence of germ plasm organizers. PLoS Genet 14, e1007696 (2018).

43. A. Jenny et al., A translation-independent role of oskar RNA in early Drosophila oogenesis. Development 133, 2827–2833 (2006).

44. M. Kanke et al., oskar RNA plays multiple noncoding roles to support oogenesis and maintain integrity of the germline/soma distinction. RNA 21, 1096–1109 (2015).

45. H. Jambor, C. Brunel, A. Ephrussi, Dimerization of oskar 3’ UTRs promotes hitchhiking for RNA localization in the Drosophila oocyte. RNA 17, 2049–2057 (2011).

46. H. Jambor, S. Mueller, S. L. Bullock, A. Ephrussi, A stem-loop structure directs oskar mRNA to microtubule minus ends. RNA 20, 429–439 (2014).

47. M. Zubradt et al., DMS-MaPseq for genome-wide or targeted RNA structure probing in vivo. Nat Methods 14, 75–82 (2017).

48. K. Bourtzis, T. A. Miller, Eds., Insect Symbiosis, (CRC Press, Boca Raton, FL, 2006), vol. 3, pp. 304.

49. S. Chen, Y. E. Zhang, M. Long, New genes in *Drosophila* quickly become essential. Science 330, 1682–1685 (2010).

50. G. Cornelis et al., Ancestral capture of syncytin-Car1, a fusogenic endogenous retroviral envelope gene involved in placentation and conserved in Carnivora. Proc. Natl. Acad. Sci. USA 109, E432–441 (2012).

51. F. Husnik, J. P. McCutcheon, Functional horizontal gene transfer from bacteria to eukaryotes. Nat Rev Microbiol 16, 67–79 (2018).

52. I. Di Lelio et al., Evolution of an insect immune barrier through horizontal gene transfer mediated by a parasitic wasp. PLoS Genet 15, e1007998 (2019).

53. N. Wybouw, Y. Pauchet, D. G. Heckel, T. Van Leeuwen, Horizontal Gene Transfer Contributes to the Evolution of Arthropod Herbivory. Genome Biol. Evol. 8, 1785–1801 (2016).

54. D. G. Quispe-Huamanquispe, G. Gheysen, J. F. Kreuze, Horizontal Gene Transfer Contributes to Plant Evolution: The Case of Agrobacterium T-DNAs. Front Plant Sci 8, 2015 (2017).

55. L. Boto, Horizontal gene transfer in the acquisition of novel traits by metazoans. Proc Biol Sci 281, 20132450 (2014).

56. A. C. Wilson, R. P. Duncan, Signatures of host/symbiont genome coevolution in insect nutritional endosymbioses. Proc Natl Acad Sci U S A 112, 10255–10261 (2015).

57. S. Lopez-Madrigal, R. Gil, Et tu, Brute? Not Even Intracellular Mutualistic Symbionts Escape Horizontal Gene Transfer. Genes (Basel) 8, (2017).

58. N. A. Provorov, O. P. Onischuk, Microbial Symbionts of Insects: Genetic Organization, Adaptive Role, and Evolution. Mikrobiologiya 87, 151–163 (2018).

59. M. Shelomi et al., Horizontal Gene Transfer of Pectinases from Bacteria Preceded the Diversification of Stick and Leaf Insects. Sci Rep 6, 26388 (2016).

60. Z. Zeng et al., Bacterial endosymbiont Cardinium cSfur genome sequence provides insights for understanding the symbiotic relationship in Sogatella furcifera host. BMC Genomics 19, 688 (2018).

61. D. C. Bublitz et al., Peptidoglycan Production by an Insect-Bacterial Mosaic. Cell 179, 703–712 e707 (2019).

62. X. Xu, J. L. Brechbiel, E. R. Gavis, Dynein-Dependent Transport of *nanos* RNA in *Drosophila* Sensory Neurons Requires Rumpelstiltskin and the Germ Plasm Organizer Oskar. J. Neurosci. 33, 14791–14800 (2013).

63. J. A. Rees, K. Cranston, Automated assembly of a reference taxonomy for phylogenetic data synthesis. Biodiversity Data Journal 5, e12581 (2017).

64. S. F. Altschul, W. Gish, W. Miller, E. W. Myers, D. J. Lipman, Basic local alignment search tool. J. Mol. Biol. 215, 403–410 (1990).

65. H. Aspöck et al. (2018).

66. M. A. Benton, N. J. Kenny, K. H. Conrads, S. Roth, J. A. Lynch, Deep, Staged Transcriptomic Resources for the Novel Coleopteran Models Atrachya menetriesi and Callosobruchus maculatus. PLoS ONE 11, e0167431 (2016).

67. R. C. Edgar, MUSCLE: a multiple sequence alignment method with reduced time and space complexity. BMC Bioinformatics 5, 113 (2004).

68. S. R. Eddy. (2007).

69. M. Stanke, R. Steinkamp, S. Waack, B. Morgenstern, AUGUSTUS: a web server for gene finding in eukaryotes. Nucleic Acids Res. 32, W309–312 (2004).

70. I. Korf, Gene finding in novel genomes. BMC Bioinformatics 5, 59 (2004).

71. A. Ahuja, C. G. Extavour, Patterns of molecular evolution of the germ line specification gene *oskar* suggest that a novel domain may contribute to functional divergence in *Drosophila*. Dev. Genes Evol. 222, 65–77 (2014).

72. U. Consortium, The Universal Protein Resource (UniProt) 2009. Nucleic Acids Res. 37, D169–174 (2009).

73. J. A. Gerlt et al., Enzyme Function Initiative-Enzyme Similarity Tool (EFI-EST): A web tool for generating protein sequence similarity networks. Biochimica et Biophysica Acta (BBA) - Proteins and Proteomics 1854, 1019–1037 (2015).

74. P. Shannon et al., Cytoscape: a software environment for integrated models of biomolecular interaction networks. Genome Res. 13, 2498–2504 (2003).

75. S. Capella-Gutierrez, J. M. Silla-Martinez, T. Gabaldon, trimAl: a tool for automated alignment trimming in large-scale phylogenetic analyses. Bioinformatics 25, 1972–1973 (2009).

76. A. Stamatakis, RAxML version 8: a tool for phylogenetic analysis and post-analysis of large phylogenies. Bioinformatics 30, 1312–1313 (2014).

77. J. P. Huelsenbeck, F. Ronquist, MRBAYES: Bayesian inference of phylogenetic trees. Bioinformatics 17, 754–755 (2001).

78. O. Robinson, D. Dylus, C. Dessimoz, Phylo.io: Interactive Viewing and Comparison of Large Phylogenetic Trees on the Web. Mol. Biol. Evol. 33, 2163–2166 (2016).

79. A. Loytynoja, Phylogeny-aware alignment with PRANK. Methods Mol. Biol. 1079, 155–170 (2014).

80. C. Notredame, D. G. Higgins, J. Heringa, T-Coffee: A novel method for fast and accurate multiple sequence alignment. J. Mol. Biol. 302, 205–217 (2000).

